# Bioprospection of culturable soil-borne bacteria with biotechnological potential for use in priming defense

**DOI:** 10.1101/2025.04.21.649657

**Authors:** Mariela González - Arriagada, Jaime Ortega, Jorge Torres, Yoelvis Sulbaran, Brynelly Bastidas, Patricia Morales-Montero, Sebastian Flores, Uri Aceituno-Valenzuela, Andree Álvarez, Rodrigo Contreras-Soto, Ernesto San Blas, Mauricio Latorre Mora, Lorena Pizarro Arcos

## Abstract

**Background:** The plant root microbiome is central to disease resistance and stress resilience. In intensive tomato production, prolonged agrochemical use disrupts microbial communities, reducing their protective functions and enabling pathogen establishment.

**Methods:** We integrated 16S rRNA amplicon sequencing with culture-dependent isolation to analyze microbiome shifts in tomato plants across healthy, asymptomatic, and symptomatic states in a nematode-infested field. Network analysis and machine learning were used to identify key taxa. Isolates were screened for plant growth-promoting rhizobacteria (PGPR) and nematicidal activity, and selected strains were evaluated in planta under pathogen challenge.

**Results:** Microbial diversity and community complexity declined with disease severity. *Gaiella occulta* emerged as a potential biomarker of plant health. From 223 isolates, 45 strains exhibited PGPR and nematicidal traits. Ten were tested in tomato plants, where treatments conferred systemic resistance to *Pseudomonas syringae* pv *tomato* without fitness cost. Four strains, primarily *Pseudomonas* and *Bacillus*, triggered immune priming, enhanced root development, and three of them were co-isolated from a single asymptomatic plant.

**Conclusions:** Our findings highlight the potential of targeted bacterial consortia to restore microbiome balance and activate immune responses in tomato. These results support the rational design of synthetic microbial communities (SynComs) for sustainable, microbiome-based crop protection.

## INTRODUCTION

Harnessing plant-associated microbiomes stands as a promising frontier in sustainable agriculture and biotechnology, offering tangible solutions to global food security, climate resilience, and the reduction of agrochemical use (Trivedi et al., 2021). Central to this approach is the intricate relationship between plants and soil microbes, shaped by evolutionary interactions that contribute significantly to plant resilience. Suppressive soils, for instance, provide compelling evidence of microbiomes actively defending plants against soil-borne pathogens through microbial competition or by priming the plant immune system to trigger systemic resistance (Berg et al., 2016; Pegg et al., 2019; Pietersen et al., 2024). Moreover, plants can selectively modulate their surrounding microbiome, enriching beneficial microbes to suppress disease (Bakker et al., 2018; Prigigallo et al., 2022).

Despite this potential, engineering natural microbiomes remains a challenge due to their complexity and the difficulty of maintaining community structure during cultivation (Ruan et al., 2024). Recent advances in systems biology, however, have improved our understanding of plant–microbe network interactions, paving the way for the development of synthetic microbial communities (SynComs) with the potential to replicate or enhance beneficial ecological functions, such as soil-borne disease suppression (Feng et al., 2019; Bakker et al., 2018). Technological innovations such as metabarcoding and high-resolution taxonomic identification have accelerated the discovery of novel bacterial and fungal taxa (Berg et al., 2020), fueling efforts to design SynComs based on laboratory-derived "benchtop" microbiomes (Berstein, 2019).

Two main strategies have emerged for building SynComs: the transplantation of whole microbial communities from suppressive soils, and the isolation and assembly of key functional taxa. The latter approach is particularly promising, as SynComs composed of 2–23 strains, often 2 to 5, can be tailored to perform targeted microbial functions, including biofilm formation, secondary metabolite production, and induction of plant resistance (Martins et al., 2023). However, careful selection of microbial candidates is critical to avoid loss of functionality.

To address the complexity of microbiome data, artificial intelligence has become an indispensable tool. Machine learning models trained on multidimensional phenotypic and genomic datasets, such as Random Forest classifiers, are being used to identify microbial biomarkers and differentiate key taxa within microbial communities (Huang et al., 2021; Chang et al., 2017; Khan et al., 2022). These approaches offer next-generation capabilities for microbial culturomics and SynCom design.

Despite these advances, rational SynCom design for robust and durable protection against pathogens remains an open challenge. Targeted bioprospecting of beneficial microbes represents a promising strategy, particularly for economically important crops like tomatoes. In this study, we used *Solanum lycopersicum* (tomato) as a model system to investigate immune-related microbial interactions. Our goal was to establish a standardized, reproducible framework for identifying bacterial candidates to priming the plant immune system. This work contributes toward the development of synthetic microbial communities designed to boost plant immunity and support sustainable crop production.

## RESULTS

### Bacterial root community shifts associated with phytosanitary statuses

The bacterial communities associated with tomatoes comprised 28,506 amplicon sequence variants (ASVs) across fifty-five samples, representing the three phytosanitary statuses: healthy (He, n=16), asymptomatic (As, n=19), and symptomatic (S, n=20).

Diversity analyses revealed that bacterial community composition remained largely conserved across phytosanitary statuses; the main difference was observed in species dominance, where the transition from He to As to S was marked by a decline in diversity and an increased prevalence of a few dominant ASVs (Fig S1). The Venn diagram further highlighted this trend, showing that 27,647 ASVs were shared across all conditions (Fig S2). Nevertheless, the shift in the phytosanitary statuses showed greater stability in the transition from AS to S, whereas a more subtle change was observed from He to As. Notably, healthy tomatoes harbored forty-one exclusive ASVs, whereas the asymptomatic and symptomatic plants exhibited fewer than twenty ASVs reinforcing their higher similarity. Despite these shared ASVs, beta diversity analysis revealed significant differences in microbial composition between groups (PERMANOVA, F = 5.77, p = 0.001), highlighting the effect of plant statuses on bacterial communities. The bacterial communities were dominated by four major phyla: Actinobacteria, Chloroflexi, Firmicutes, and Proteobacteria (Fig S3). Of particular interest was the enrichment of plant-growth-promoting rhizobacteria (PGPR), particularly within the Gammaproteobacteria and Bacilli class (Fig S4 and S5). Notably, a progressive increase in the relative abundance of *Pseudomonas* spp. was observed with disease progression, while Bacillaceae remained stable, constituting over 63% of the bacterial community in all statuses.

The networks comprised 439 taxa from healthy, asymptomatic, and symptomatic tomato plants, representing the core microbial community after filtering. The microbiome reveals a progressive disruption of bacterial community structure (Table 1). In asymptomatic plants, network fragmentation and loss of key ASVs suggest a weakening microbiome stability. Symptomatic plants exhibit increased modularity and a rise in positive interactions, indicating a shift towards a reorganized microbial community potentially dominated by opportunistic bacteria. These findings suggest that microbiome alterations precede symptom development, highlighting its potential as an early indicator of plant health decline.

**Table 1.**
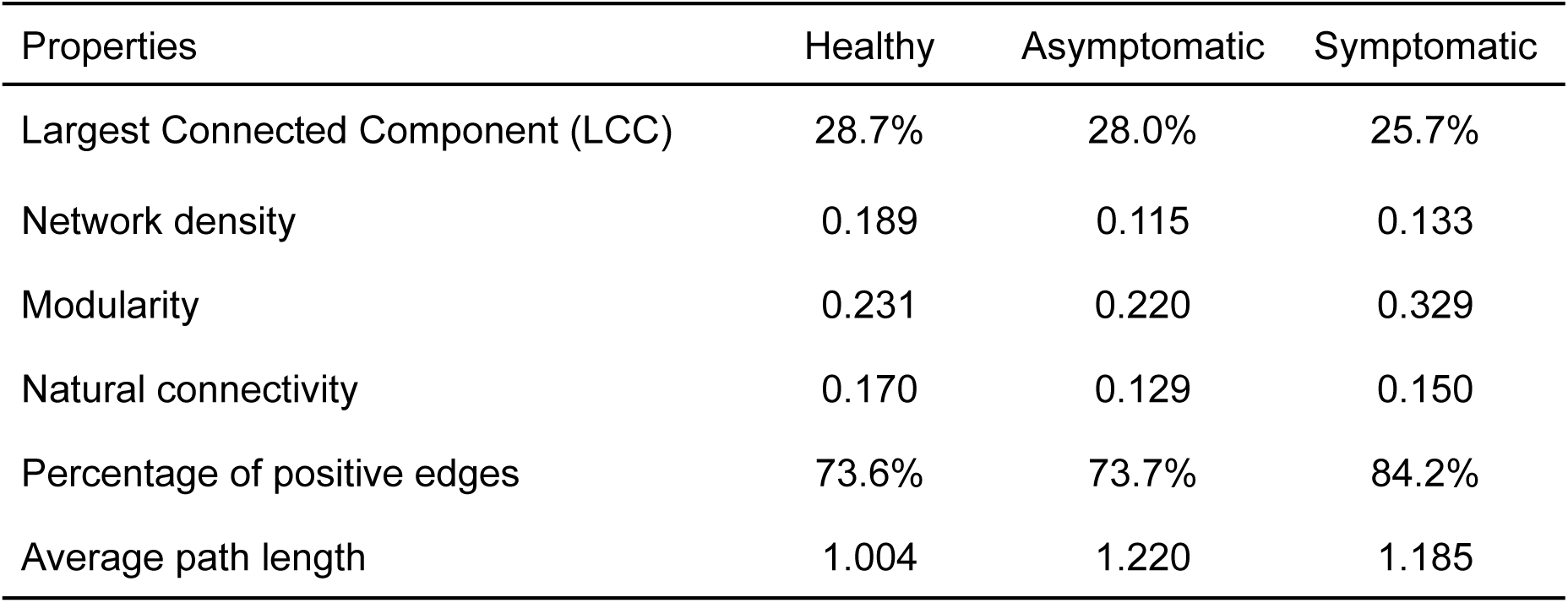
Topological properties of the networks comparing the phytopathogenic statuses.

| Properties | Healthy | Asymptomatic | Symptomatic |
| --- | --- | --- | --- |
| Largest Connected Component (LCC) | 28.7% | 28.0% | 25.7% |
| Network density | 0.189 | 0.115 | 0.133 |
| Modularity | 0.231 | 0.220 | 0.329 |
| Natural connectivity | 0.170 | 0.129 | 0.150 |
| Percentage of positive edges | 73.6% | 73.7% | 84.2% |
| Average path length | 1.004 | 1.220 | 1.185 |

The Random Forest model was built using 55 samples and 1,792 ASVs, yielding an out-of-bag (OOB) error rate of 25.45%, suggesting good model performance. Cross-validation was performed to assess model reliability, yielding a classification accuracy of 77% with a kappa value of 0.64. The top 20 most important bacterial ASVs contributing to classification predominantly belonged to Actinobacteria (67%), Proteobacteria (27%), and Firmicutes (0,3%); the class Thermoleophilia (36%) and Alphaproteobacteria (23%) were the most frequent. The PCA performed with these 20 main ASVs (Fig 1) showed that asymptomatic and symptomatic samples formed an overlapping cluster, while healthy samples were distinctly separated.

**Fig 1.**
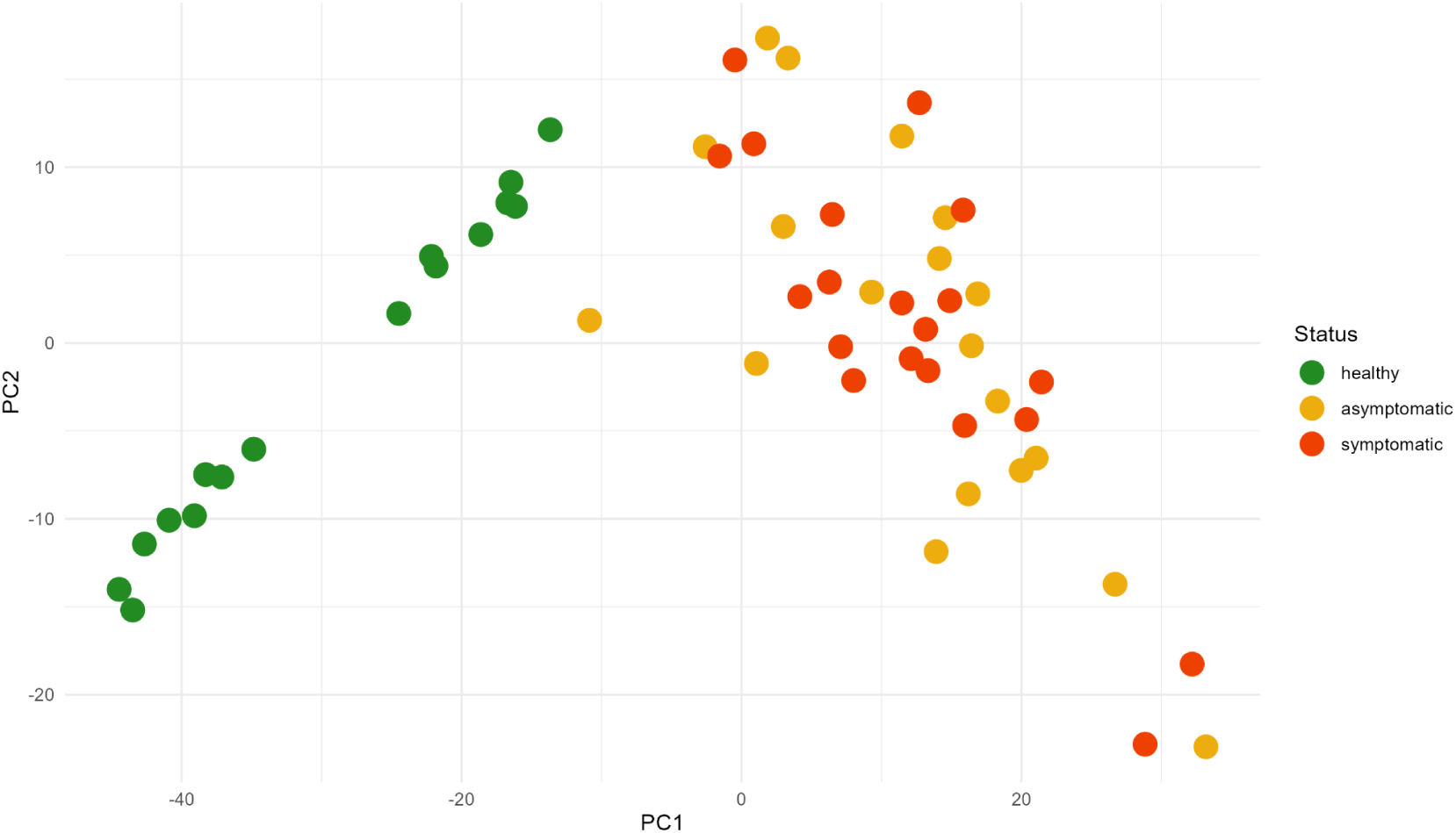
Principal component analysis (PCA) based on the 20 most important amplicon sequence variants (ASVs) identified by the Random Forest model. Each dot represents a tomato soil sample colored according to its phytopathogenic status. The first principal components (PC1 and PC2) explained 74% and 15% of the total variance.

### Plant growth-promoting rhizobacteria profile

A total of 223 bacterial strains (Table S1) were isolated from the rhizosphere of tomato plants, with 42% derived from healthy (S) plants and 58% from asymptomatic (As) individuals, reflecting a diverse microbial reservoir potentially linked to plant health status. Functional screening revealed plant growth-promoting rhizobacteria (PGPR) traits across several strains. Indole-3-acetic acid (IAA) production was detected in 21 strains, predominantly from asymptomatic plants (71%), surpassing the IAA production threshold of *Pseudomonas putida* ATCC 12633, indicating a potential correlation between auxin biosynthesis and host condition (Fig S6, Table S1). ACC deaminase activity, a key trait in stress alleviation, was observed in 22 strains, with distinct temporal patterns of enzymatic activity; strains from healthy plants responded earlier (48 h), whereas those from asymptomatic plants exhibited a delayed yet sustained response (96 h) (Fig S7). Osmotic stress resistance, tested via PEG_6000_ exposure, was observed in 70% of the isolates, suggesting a widespread capacity for drought resilience, with a slight predominance in strains from asymptomatic plants (Fig S8). Nitrogen fixation (Fig S9) and phosphate solubilization (Fig S10) were less common traits, present in 5 and 8 strains respectively, yet showed distinct distribution patterns: phosphate-solubilizing strains were more frequently isolated from healthy plants, while nitrogen-fixing bacteria were evenly distributed. Collectively, these findings reveal a functional mosaic of PGPR traits among rhizosphere-associated bacterial communities, with asymptomatic and healthy plants harboring strains with differential capacities that may contribute to growth promotion and stress mitigation.

### Oxidative burst measurement (ROS)

Twenty-one bacterial strains were identified as inducers of oxidative bursts (Fig S11), suggesting their potential role in triggering induced systemic resistance (ISR) in plants. Among these strains, only one was isolated from healthy plants, while the remaining twenty were obtained from asymptomatic plants. When defining *Pst* DC3000 as a reference (1), five strains (AS60, AS77, AS85, AS89, and AS109) exhibited a 2-fold increase in relative light unit (RLU) emission, and three strains (AS87, AS104, and AS115) showed a 3-fold increase. These results indicate that specific bacterial strains from asymptomatic plants may strongly contribute to ISR activation.

Forty-five bacterial strains were selected based on different criteria (above) for the profile of a *Plant growth-promoting rhizobacteria*

### Nematicide activity

Out of the 45 bacterial strains tested (Fig S12), eight demonstrated comparable or superior efficacy to the commercial nematicide (BAFEX-N®) for controlling *Meloidogyne* spp. second-stage juveniles (J2) after 72 hours of incubation. Among these, three strains—S05, AS66, and AS85, all belonging to the Bacillaceae family—showed statistically significant differences compared to the commercial product (BAFEX-N®), with higher nematicidal activity.

### Pathogenicity assay

Five days post-inoculation, 13 bacterial strains induced symptoms of yellowing and necrosis at the edges of the leaf disks, resembling those caused by *Pst*. In contrast, 32 bacterial strains exhibited slight oxidation at the edges or no symptoms. Leaf disks inoculated with the mock treatment showed no symptoms.

### Assign taxonomy

Forty-five bacterial strains were sequenced using the 16S SSU rRNA gene. Taxonomic identification based on NCBI BLAST nucleotide similarity revealed that the isolates belonged to the phyla Bacillota, Pseudomonadota, and Bacteroidota. Among them, the most frequently isolated genera, *Bacillus* and *Pseudomonas*, each representing 31% (14 strains), and *Calidifontibacillus erzurumensis* with 15% (7 strains) were the most prevalent species. In total, 25 distinct taxa were identified. Phylogenetic relationships were inferred using Maximum Likelihood (ML) and Bayesian Inference (BI) analyses, based on 1,411 amino acid sites, including 560 constant sites, 505 parsimony-informative sites, and 849 distinct patterns. The best-fit evolutionary model, determined by the Bayesian Information Criterion (BIC), was GTR. The phylogenetic tree was rooted and revealed two well-defined clades: clade one corresponding exclusively to *Pseudomonas, Aquipseudomonas,* and *Ectopseudomonas*, and another, clade two, containing *Bacillus*, *Peribacillus,* and *Calidifontibacillus*.

Functional trait analysis showed that phosphate solubilization and nitrogen fixation were traits characteristic of *Pseudomonas* strains in this study, while ACC deaminase activity, osmotic stress resistance, and induction of systemic resistance were distributed across multiple taxa throughout the phylogenetic tree (Fig 2)

**Figure 2.**
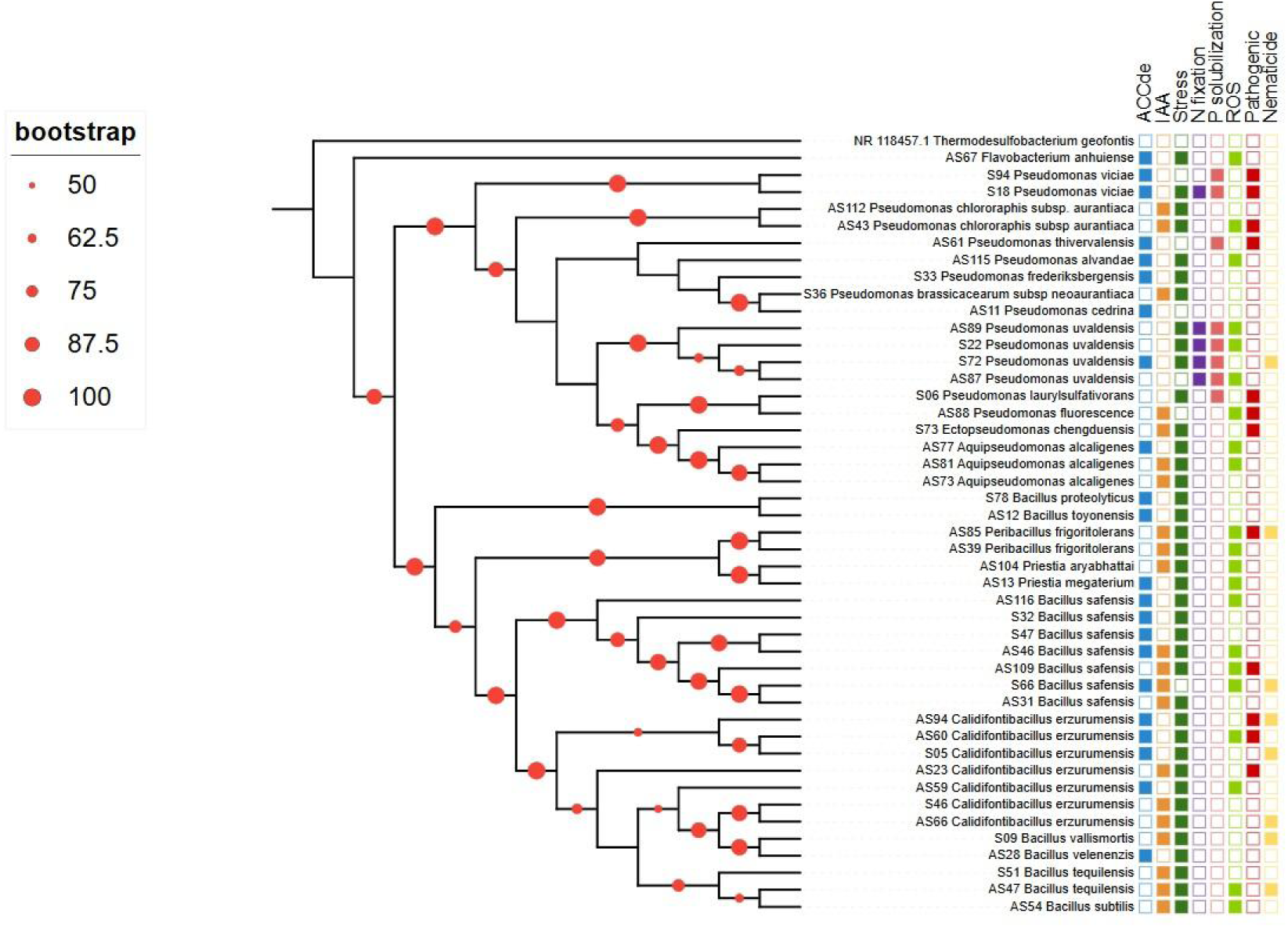
Phylogenetic tree of 45 root-associated bacterial strains isolated from healthy (S) and asymptomatic (AS) tomato plants. The tree was constructed using the maximum likelihood (ML) method, with *Thermosulfobacterium geofontis* (NR 118457.1) as the outgroup. Bootstrap support values (>50%) are indicated by dots on the branches. The heatmap represents the functional profile of each strain for key plant growth-promoting (PGP) traits and other activities: 1-aminocyclopropane-1-carboxylic acid (ACC) deaminase activity (ACCde), indole acetic acid (IAA) production, osmotic stress resistance (Stress), nitrogen fixation (N fixation), phosphate solubilization (P solubilization), oxidative burst response (ROS), pathogenicity, and nematicidal activity. Bacterial strains are grouped into major phylogenetic clades, with taxonomic classifications indicated. Raw data are available in Table SXXX.

### Bacterial strain performance

Ten bacterial strains (Table 2 were selected based on the presence of 3 to 5 tested traits to be tested for pathogen resistance and growth promotion.

**Table 2.** Ten bacterial strains (ID) selected from plants in different statuses were tested in plants for their growth-promoting activity.

| Treatment | Plant | ID |
| --- | --- | --- |
| T1 | C13 | *Pseudomonas uvaldensis* |
| T2 | C3 | *Pseudomonas laurylsulfativorans* |
| T3 | C18 | *Pseudomonas brassicacearum* subsp. *neoaurantiaca* |
| T4 | C17 | *Pseudomonas frederiksbergensis* |
| T5 | C10 | *Bacillus safensis* |
| T6 | A16 | *Bacillus safensis* |
| T7 | A17 | *Pseudomonas chlororaphis* subsp. *aurantiaca* |
| T8 | A17 | *Pseudomonas alvandae* |
| T9 | A17 | *Bacillus safensis* |
| T10 | A9 | *Peribacillus frigoritolerans* |

Tomatoes infected with *Pst* showed dark brown-to-black spots 5 to 7 days post-inoculation, and differences in severity scale were visible to the naked eye. Statistical analysis revealed significant differences in disease severity across six treatments (T3, T5, T6, T8, T9, and T10), which notably reduced CFU/g of leaf tissue compared to natural infection with *Pst* (Fig S14). The negative control without *Pst* infection did not exhibit any symptoms.

Tomato plants inoculated with *Pst* exhibited significant differences in root growth when subjected to treatments T3, T4, T5, T6, T7, T8, and T9 (Table S2). Treatments T4, T7, T8, and T9 demonstrated superior performance in at least four out of the six root parameters tested, including projection area, surface area, average diameter, and root volume, being T4 and T9 exhibiting the most frequent and pronounced improvements. Notably, three treatments (T7, T8, and T9) with the best results originated from the same asymptomatic plant, A17, and corresponded to two distinct bacterial genera. Using the K-means clustering method (Fig 3), most repetitions of treatments T4, T7, and T9 were grouped into cluster 3. In contrast, treatment T8 was partitioned into three distinct clusters, likely due to variations in performance across the different measurements. A scatter plot based on root volume and average diameter of root showed how *Pseudomonas* and *Bacillus* were associated with the high performance of those variables (Fig S15) and corresponded to T4, T7, and T9 treatments.

**Fig 3:**
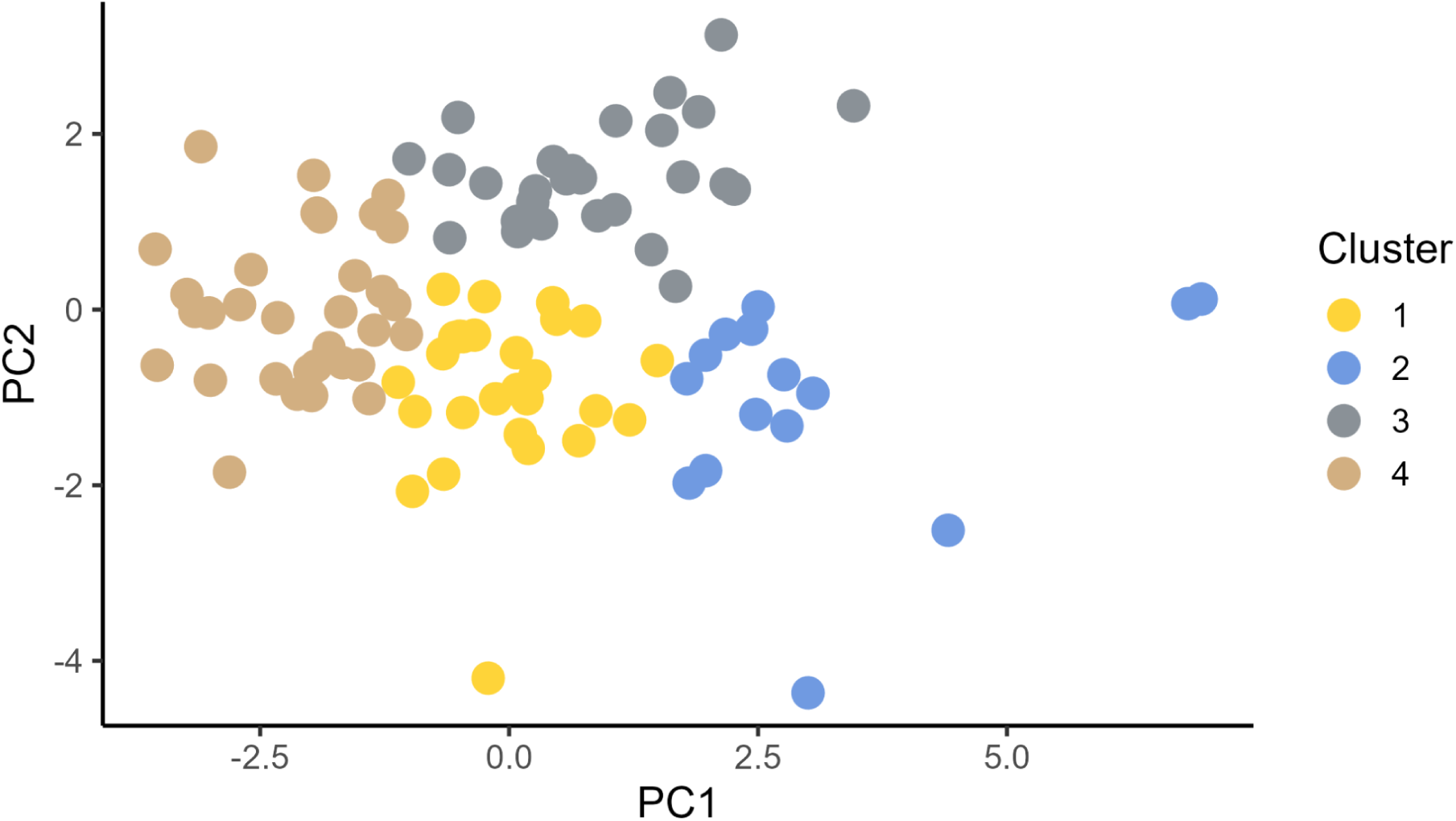
K-means clustering analysis of bacterial strains, based on root measurements in response to pathogen resistance against *Pseudomonas syringae* pv. *tomato* inoculation. Strains are color-coded according to their assigned clusters, with the optimal number of clusters determined using the elbow method

In terms of biostimulant activity without infection, the only treatment that exhibited a significant difference compared to the control was T2, which resulted in negative effects. Specifically, T2 was associated with reduced root length, surface area, and projection area, indicating that this treatment may inhibit root growth rather than promote it.

## DISCUSSION

Plant-associated root microbiome enhances disease resistance (Berendsen et al., 2018; Yin et al., 2020). However, in intensive greenhouse tomato cultivation, which relies heavily on pesticides and fertilizers, indiscriminate chemical use can significantly alter the composition and functionality of root-associated microbiota (Lamelas et al., 2020). These modifications are closely linked to intensifying the disease control strategies, potentially disrupting beneficial microbial communities and their protective effects on plant health (Saleem et al., 2019). In this research, the Inverse Simpson’s Index decreased progressively as tomato plants transitioned from healthy to symptomatic states, indicating a reduction in microbial diversity and a shift towards community dominance by fewer taxa. Similar results were observed in Wolfgang et al. (2019) when studying the root disease process. The decline of a stable microbiome may be associated with the loss of beneficial bacteria, asymptomatic plants, and reduced community resilience, creating an environment that favors opportunistic or pathogenic bacteria proliferation. As symptoms develop, the microbial network undergoes a structural reorganization, becoming dominated by a different set of bacteria. The interaction tomato-soil-*Meloidogyne spp* microbiome reveals complex microbial dynamics (Topalović et al., 2020a). While no specific taxonomy cluster was identified to distinguish among the three phytopathological states, however, distinctions were observed between healthy and asymptomatic-symptomatic plants showed modifications in he shift of healthy to asymptomatic status; a similar pattern of compartmentalization has been reported in other studies among healthy and rot knot disease (Lamelas et al., 2020), leads to a restructured root microbiome specialized in the gall tissue (Tian et al., 2015). The co-occurrence network analysis revealed a progressive disruption of bacterial community structure, consistent with findings from endosphere microbiomes of healthy, diseased, and severely diseased samples affected by *Clavibacter michiganensis* subsp. *michiganensis.* In that study, networks associated with healthy samples exhibited greater robustness than those from diseased conditions (Yin et al., 2020; Choi et al., 2020). Conversely, disease-suppressive soils often emerge following disease outbreaks, fostering the assembly of a protective root-associated microbiome that enhances plant resistance (Berendsen et al., 2018).

From the microbiome obtained from tomato plants grown in a field heavily infested with *Meloidogyne* spp, 28,506 ASVs were identified with more frequency in phyla Actinobacteria, Chloroflexi, Firmicutes, and Proteobacteria. Within the phylum Actinobacteria, the *Microbacteriaceae* family was highly abundant and has been associated with bacteria that adhere to the *Meloidogyne* spp. cuticle (Topalović et al., 2020a). Additionally, *Gaiella occulta* emerged as a key taxon for predicting the phytosanitary status in our random forest model. However, this species is known to exhibit poor growth on standard agar media (Albuquerque et al., 2011), which may explain the absence of cultured isolates in our study. Notably, *G. occulta* has been described as having biotechnological potential, including phosphate solubilization, organic matter decomposition, and antibiotic production for microbial inhibition (Severino et al., 2019), identifying its ecological role within the tomato microbiome is essential. On the other hand, Firmicutes and Proteobacteria are recognized by their PGPR activity. From culture-dependent, a total of 223 isolates were obtained from healthy and asymptomatic soil. These isolates were subsequently characterized for bioestimulant potential (Gaete et al., 2020; Maza et al., 2019; Ali and Khan, 2021). Interestingly, among the 45 bacterial strains exhibiting a PGPR profile and nematicidal activity, a predominance of two bacterial families, *Pseudomonadaceae* and *Bacillaceae,* was coincident with the results obtained from microbiome characterization.

Eight bacterial strains from the *Bacillaceae* family exhibited nematicidal activity comparable to or exceeding that of a commercial product when tested against infective *Meloidogyne* J2 juveniles. The biocontrol of root-knot focused on preventing infection, and several strategies have been identified in *Bacillus* spp. as a direct mechanism, including their ability to form biofilms on roots, which act as physical barriers against nematode penetration. Indirect mechanisms include modifying root exudates, thereby disrupting nematode recognition, altering the development of feeding sites, influencing to female-to-male sex ratio within root tissues, or promoting plant growth (Mhatre et al., 2019). In this research, analysis of the samples revealed the presence of antagonistic microbial consortia commonly associated with suppressive soils, including *Pasteuria*, *Bacillus*, *Pseudomonas*, *Rhizobium*, *Streptomyces*, *Arthrobacter*, *Lysobacter*, and *Variovorax*, consistent with previous reports from RKNs suppressive (Topalović et al., 2020b).

Ten bacterial strains were selected to evaluate their performance in tomato plants; half were subjected to a foliar pathogen, *Pst*. Interestingly, non-inoculated tomato plants showed non-significant differences compared to the control. On the other hand, *Pst*-inoculated plants that received bacterial treatment remained healthy, primarily exhibiting leaf abscission as a active protective mechanism (Patharkar et al., 2017; Kong and Yang, 2023), without a fitness cost associated to the root development related to growth-defense trade-off (Pietersen et al., 2014; He et al., 2022), which could be associated to a phenomenon that growth–defense trade-offs can occur in an organ- or tissue-specific, effectively minimizing conflicts at the whole-plant level (Conrath et al., 2015; He et al., 2022). This response was significantly more pronounced in treated plants than in those inoculated with *Pst* without treatment or in healthy controls. Similar results have been reported in *Arabidopsis*, where foliar infection with a biotrophic pathogen triggers systemic signals to the roots, promoting the growth of specific microbial species in the rhizosphere; this consortia not only induce resistance but also enhances plant growth (Berendsen et al., 2018). Tomato plants pre-treated with beneficial biocontrol agents exhibited increased resistance to infection by RKNs associated with the activation of defense genes (PR-1, PR-3, PR-5, ACO) until 12d after inoculation (Molinari and Leonetti, 2019). Induced systemic resistance (ISR) is a response stimulation of the host’s immune system modulated by root-associated microorganisms or interaction between microorganisms (Pietersen et al., 2014; Berendsen et al., 2018; Topalović et al., 2020b). This mechanism enhances the plant’s protection against a broad spectrum of pathogens (Walter et al., 2013), and it is associated with defense priming, an adaptive strategy that prepares the plant to respond more rapidly and effectively to future attacks, minimizing damage in hostile environments (Pietersen et al., 2014), adding to the fitness costs of priming are lower than those of constitutively activated defenses (Conrath et al., 2015). Leaves of *Arabidopsis* infected with *Pst* generate an attraction of *Bacillus subtilis* FB17 on the root, and trigger ISR protection on non-infected parts of the plant, also enhancing stomatal closure, delaying the disease progression (Rudrappa et al., 2008)

Molecules with microbial origin, such as flagellin, chitin, lipopolysaccharides, among others, known as microbe-associated molecular patterns, act as biotic inducers of ISR (Reglinski et al., 2023), and are released in pathogen attack (Boutrot and Zipfel, 2017). Later, pattern recognition receptors (PRRs) in the plasma membrane of the cell recognize the inducer, and a complex biochemical signaling cascade is triggered, activating the transcriptional reprogramming defense mechanism (Reglinski et al., 2023), which includes stomatal closure, production of antimicrobial components, cell-wall fortifications, production of toxic ROS (Li et al. 2020; Topalović et al., 2020b). This could be induced through the biotic inducer application, resulting in a “primed” effect against the pathogen (Conrath et al. 2015) which persists, in general, for 2 - 4 weeks after application (Walters et al. 2013).

Among the ten bacterial strains, four exhibited highly significant and promising performance in pathogen interaction, with representatives from the genera *Pseudomonas* and *Bacillus;* both are well-known for their mutualistic association with roots, where they prime the plant immune system, enhancing defense mechanisms without triggering energy-intensive responses (Pieterse et al., 2014). Upon root colonization, *Pseudomonas* and *Bacillus* alter the root architecture, show an increase in root hair length, plant biomass production, and abundant lateral root formation due to the auxin-dependent root developmental program (Ali and Khan, 2021). These strains displayed key characteristics such as ACC deaminase production, IAA synthesis, and ROS activation. Notably, three of these four strains were isolated from the same asymptomatic plant, which could be indicative of a potential for these strains to conform to the synthetic community with high performance. Different plant species release distinct root exudates that selectively attract specific PGPR strains, and the chemical composition of these exudates can vary not only between species but also among cultivars within the same species (Tripathi et al., 2024). This suggests the relevance of the asymptomatic plant’s exudate profile in fostering the selective recruitment of high-performing strains. The specific set of microbes in the soil product of a selective enrichment during a pathogen invasion due to the “cry for help” strategy from the plants as a defensive tactic (Bakker et al., 2018; Liu et al., 2024) results in the formation of long-lasting disease suppressive soil (Mendes et al., 2011), for that reason, creating a synthetic microbial community that accurately mimics the natural microcosm is a challenging task, and developing a robust network of interactions holds promise as a potential strategy for controlling disease (Yin et al., 2020). Also, SynCom is a useful tool as a biosensor, using real-time PCR, to detect specific microorganisms present in soil, once their ecological role in the microbiome is predicted (Yuan et al., 2023).

The major societal challenge of producing more food with reduced fertilizer and agrochemical inputs in crop protection has heightened awareness of the critical role played by the root microbiome in maintaining plant health under current agricultural and horticultural practices. In this context, strategies aimed at activating plant innate immunity represent a promising, environmentally friendly approach to pest management, offering a sustainable and economically viable solution for farmers. Systems biology approaches are a useful tool for the development of next-generation inducers by leveraging an increasingly detailed understanding of plant recognition receptors and defense signaling pathways.

## CONCLUSION

This study highlights the significant potential of leveraging plant-associated microbiomes for sustainable crop protection and plant health management. By focusing on the intricate interactions between plants and their root microbiomes, we demonstrate that microbial communities can enhance disease resistance through mechanisms such as immune system priming and pathogen suppression. The identification and characterization of beneficial microbes, particularly those with plant growth-promoting and nematicidal activities, provide a promising strategy for developing synthetic microbial communities (SynComs) that mimic natural disease-suppressive soils. Our findings emphasize the importance of selecting key microbial taxa to build effective SynComs, capable of enhancing plant immunity without imposing fitness costs. Moreover, the use of machine learning models and advanced biotechnological tools for microbiome analysis offers a robust framework for future bioprospecting efforts. As agricultural practices shift towards more sustainable, environmentally friendly approaches, the integration of SynComs and microbiome-based technologies promises to play a critical role in improving crop resilience, reducing reliance on agrochemicals, and fostering long-term agricultural sustainability.

## METHODS

### Sampling area

A tomato (*Solanum lycopersicum* XX) cv. "Alamina" grafted onto "Maxifort" greenfield was visited in a location in Pichidegua (−34°18’07.8’’S −71°21’57.8’’W) during the mid-production season (January 2024). The area was previously recorded as infested with root-knot nematodes, *Meloidogyne* spp. Soil samples (500 g) were collected from near the roots of tomato plants, with the top 2 cm of soil removed. Ethanol was applied to the shovel between samples to ensure sterile conditions. Tomato plants were classified as healthy, asymptomatic, or symptomatic based on the presence of root-knot nematodes, and their phytosanitary status was noted. Plants classified as healthy showed no visible symptoms and no presence of root-knot nematodes. Asymptomatic plants exhibited a healthy phenotype but tested positive for the presence of root-knot nematodes. Symptomatic plants displayed visible disease symptoms and confirmed presence of root-knot nematodes. Twenty samples were collected per condition. The samples were stored at −20 °C for further analysis.

### Bacterial root community shifts associated with phytosanitary statuses

#### Metabarcoding

0.3 g of soil, from samples of the three phytosanitary statuses, was taken for DNA extraction using the *Quick*-DNA Fecal/Soil Microbe DNA Miniprep Kit (Zymo Research) following the manufacturer’s protocol. DNA quality was assessed using the 260/280 nm ratio in a NanoDrop® One spectrophotometer (Thermo Fisher Scientific), and the concentration was quantified with a Qubit 2.0 fluorometer (Thermo Fisher Scientific) using the dsDNA HS assay kit. The samples were stored at −20°C until sequencing.

DNA samples were subjected to high-throughput sequencing by Mr. DNA using the Illumina MiSeq platform. The V4 hyper-variable region of the 16S SSU rRNA gene was amplified with 515f-806r primers, and sequencing was performed in both forward and reverse directions via paired-end technology.

#### Preprocessing data

Sequencing data were analyzed using a customized bioinformatics pipeline primarily based on the VSEARCH tool v.2.14 (Edgar and Flyvbjerg, 2015). Raw reads obtained from the Illumina platform were deposited in the National Center for Biotechnology Information (NCBI) and are accessible under Bioproject accession number PRJNA1238788. During preprocessing, PhiX sequences and primer fragments were removed using Bowtie2 v.2.5.5 (Martin, 2011) and Cutadapt v.3.4 (Langmead and Salzberg, 2012). Paired-end reads were subsequently merged with the fastq_mergepairs algorithm, and low-quality sequences were filtered by applying a maximum expected error threshold of 1 using fastq_filter in VSEARCH. The merged high-quality sequences were then dereplicated using the derep_fullength algorithm in VSEARCH. Amplicon Sequence Variants (ASVs) were inferred using the UNOISE algorithm (Edgar, 2016), and potential chimeric sequences were identified and removed using the uchime algorithm in VSEARCH (Edgar et al., 2011). To confirm that the retained ASVs corresponded to the 16S rRNA gene, they were screened using Metaxa2 v.2.2.3 (Bengtsson-Palme et al., 2015). The final ASV table was generated by mapping the filtered sequences against validated ASV centroids using the usearch_global algorithm in VSEARCH. Taxonomic classification was performed using the naïve Bayesian classifier Syntax (Edgar, 2016), aligning the ASVs against the Silva v.132 databases for 16S rRNA (Quast et al., 2013). Finally, the ASV table and taxonomic assignments were integrated into a phyloseq object, incorporating metadata on the phytosanitary statuses of the ASVs, using the *phyloseq* v.1.5 packages (McMurdie & Holmes., 2013) in RStudio (R Core Team., 2024).

#### Bacterial communities

Diversity indices, including alpha diversity metrics (Chao1, Shannon, Inverse Simpsons, and evenness Pielou), were calculated using the *alpha* function in microbiome v.1.16.0 (Lahti and Shetty, 2022) to assess within-sample diversity. Dis(similarities) in the composition of ASVs across phytosanitary statuses were visualized through Venn diagrams generated with the *ps_venn* function in the MicEco v.0.9.19 (Russel, 2022). The relative abundances of phyla and families were estimated according to tomato statuses. To evaluate community composition differences between phytosanitary statuses, Permutational Analysis of Variance (PERMANOVA; Anderson, 2001) was performed using the *adonis* function in vegan v.2.6-8 package (Oksanen et al., 2024), with Bray-Curtis matrix distance as the dissimilarity metric. A comparative family-level network analysis was performed using NetCoMi v.1.0.2 (Peschel, 2021) with the SparCC algorithm, applying a 0.85 threshold and centered log-ratio (CLR) transformation within the *netConstruct* function. For classification, Random forest analysis (Breiman, 2001) was employed to identify root-associated bacterial communities that distinguish between tomato phytosanitary statuses (Khan et al., 2022) using the RandomForest v.4.71.2 package (Liaw et al., 2022). ASV counts were filtered across all datasets to remove noise, retaining only those with a relative abundance >0.0001. The *randomForest* function used to classify the model was set with ntree 1000 and mtry =10, which refer to the number of trees and features considered in each tree split, respectively. Cross-validation was performed using the *train* function in caret v.7.0-1 (Kuhn, 2008) to estimate model performance. The *importance* function was used to determine the predictive contribution of each ASV in the classification process. Principal Component Analysis (PCA) was performed using the *prcomp* function to reduce the dataset dimensionality and explore the relationships between the phytosanitary statuses and bacterial community composition.

### Plant growth-promoting rhizobacteria profile

Samples obtained from soils of asymptomatic and healthy tomatoes were employed. 10 g of soil was mixed with 10 mL of sterile Phosphate Buffered Saline (PBS) 1X, and incubated in a shaker for 1 h. The suspension was centrifuged at 500 rpm for 10 min. 100 µL of supernatant was plated onto two different culture media: Luria-Bertani (LB) (Maza et al., 2019) and King’s B media (King et al., 1954). The strains were incubated at 30 °C for 48 h. Colonies with distinct morphological characteristics were isolated, maintained in their respective culture media, and stored in glycerol at −80 °C.

Several tests were conducted to define the bacteria’s potential as biostimulants (Gaete et al., 2020; Maza et al., 2019; Ali and Khan, 2021). Bacterial cultures were grown overnight in LB medium supplemented with 0.2% tryptophan at 30°C under agitation at 150 rpm. The assay for indole acetic acid production, ACC deaminase activity, osmotic stress resistance, nitrogen fixation, and phosphate solubilization was performed using the OT-2 Robot (OpenTrons, New York, USA).

#### Indole acetic acid (IAA) production

IAA production was evaluated using a Salkowski colorimetric method (Gordon and Weber, 1951). Isolates were cultured in an LB medium supplemented with 0.2% L-tryptophan as a precursor to AIA. The cultures were centrifuged at 4000 rpm for 10 min (Rotor A-2-MTP centrifuge Eppendorf 5430 R). 60 µL of the supernatants were mixed with 30 µL of Salkowski’s reagent, and the cultures were incubated for 20 min at room temperature. The absorbance at 535 nm was measured before and after incubation. The LB medium turned from yellow to violet according to the concentration of AIA in the medium. The positive control was *Pseudomonas putida* ATCC 12633, recognized for the capacity to produce IAA (Liffourrena and Lucchesi, 2018), and the negative control was *Enterococcus faecalis* OG1RF, a non-produced IAA. The experiment was performed in triplicate for each bacterial strain. Bacteria were selected based on the production of IAA equal to or greater than the positive control.

#### 1-aminocyclopropane-1-carboxylic acid (ACC) deaminase activity

Strains were cultured in media with ACC as a unique nitrogen source (Penrose and Glick, 2003). The enzymatic activity of 1-aminocyclopropane-1-carboxylic acid deaminase (ACCD) converts ACC into α-ketobutyrate (α-KB) and ammonium. A 20 µL volume of the culture was seeded in 180 µL of DF (Dworkin and Foster) minimal medium supplemented with 3mM of ACC as the sole N source (Janati et al., 2023) for 4 d at 30°C. The control consisted of DF medium without ACC. Growth was evaluated at 0, 48, and 96 h using a spectrophotometer at an absorbance of 600 nm (Vega-Celedón et al., 2020), with the 0-hour time point as the control blank. The experiment was performed in triplicate for each bacterial strain. Bacteria exhibiting significant growth differences in the DF+ACC medium were selected for their capacity to synthesize ACC deaminase.

#### Osmotic stress resistance

To simulate drought stress conditions, polyethyleneglycol (PEG) 6000 was added as an osmotic stress inducer (Bouremani et al., 2024). A 20 µL of each bacterial strain was seeded into LB medium as a control for natural conditions, and LB medium was supplemented with 15% (wv^-1^) PEG6000, corresponding to an osmotic pressure of −0.49 MPa (Eswaran et al., 2024). The growth of bacterial strains was measured at an absorbance of 600 nm at 0, 48, and 96 h, comparing stress conditions to the control without PEG. The experiment was performed in three replicas for each bacterial strain. Bacterial strains that did not show significant growth differences under drought stress were selected.

#### Nitrogen fixation

The essay allows the identification of chemoheterotrophic bacteria capable of fixing atmospheric nitrogen (Grobelak et al., 2015). 5 µL of each bacterial culture was seeded into a Norris glucose nitrogen-free medium (Norris and Wulff, 1969) and incubated at 30°C for 2 d. The formation of a halo around the seed spot was considered indicative of a positive bacterial strain for nitrogen fixation.

#### Phosphate solubilization

To determine the capability of bacteria to hydrolyze inorganic and organic insoluble phosphorus to soluble forms (Kaur et al., 2024), each bacterial strain was inoculated with 5 µL in triplicate onto Pikosvkaya (PVK) medium and incubated at 30°C. After 2d, the formation of halos around the colonies was used as an indicator of phosphate solubilization by the bacterial strains.

### Oxidative burst measurement (ROS)

ROS measurement was performed as previously described (Leibman-Markus et al., 2017; Pizarro et al., 2022). Bacterial strains were grown overnight in LB medium at 28° C with shaking to 200 rpm and adjusted to an OD600 of 0.2. *Pseudomonas syringae* pv. *tomato* DC3000 (*Pst*) was used as the positive control, and the negative control was *Agrobacterium tumefaciens* GV3101(At). Leaflets from the 4th to 6th leaves of 6-7 week-old *S. lycopersicum* cv. Moneymaker plants were superficially disinfected with 70% ethanol. Leaf disks (0.6 cm in diameter) were excised, bisected, and placed in a white 96-well plate (SPL Life Sciences, Korea) pre-filled with 250 μL distilled water. Samples were incubated at room temperature for 4–6 h, ensuring the abaxial side remained in contact with the well bottom. After incubation, the water was removed, and the ROS measurement reaction was initiated by adding a 1.5X luminol-based chemiluminescent solution (HRP 37.5 µg/mL and luminol 375 µM in 200 mM KOH). Each well received 100 µL of bacterial suspension and 100 µL of the luminol solution. Luminol-induced light emission was immediately recorded using a luminometer (Turner BioSystems Veritas, California, USA), with 4s readings over 18 cycles. Total ROS production was determined by summing the average of each six technical replicates for each measurement cycle. The oxidative burst triggered by *Pst* was set as 1; this provided a maximum reference value, against which the values of all bacterial strains were compared

### Nematicide activity

Second-stage juveniles (J2) of *Meloidogyne* were extracted from infested host plant roots using the Baermann funnel method (Hussey and Barker, 1973). Identification was based on morphological characteristics observed under a stereoscopic microscope, and only motile individuals were selected for the assay. Bacterial strains were cultured in LB medium at 28°C with shaking at 150 rpm for 24 h. The bacterial strains were adjusted to OD₆₀₀ of 0.05, corresponding to approximately 10^6^ CFU/mL. The assay was performed in 24-well plates, with each well containing 600 µL of the bacterial suspension, and approximately 20 *Meloidogyne* (j2). Each bacterial strain was tested in four independent replicates, randomly distributed across the wells. *Agrobacterium tumefaciens,* LB medium, and distilled water were employed as negative controls, and two *Bacillus* spp. isolates with known nematicide activity (FB25M and FB37BR; Aballay et al., 2020), and the commercial biological nematicide BAFEX-N® (Bio Insumos Nativa SpA, 2025), were used as a positive control. Nematodes were incubated in darkness at 20°C for 72h. Nematode viability was assessed under a stereoscopic microscope, considering individuals immobile and unresponsive to mechanical stimuli like death. Mortality was expressed as a percentage of non-viable nematodes relative to the total.

### Pathogenicity assay

Bacteria strains were grown in a TSB medium (Sambrook and Russell, 2001) for 24 h at 30°C with shaking at 150 rpm. Bacterial strains were adjusted to an OD_600_ of 0.6 for inoculation. Leaf disk infection was performed as described by Lienqueo et al. (2024); the fully expanded leaflets from 8-week-old *Solanum lycopersicum* cv. "M82" plants were employed. Leaflets were superficially disinfected using 70% ethanol and air-dried. Then, a cork borer was immersed in the bacterial suspension OD_600_ 0.6 for ten seconds, and 10 leaf disks were collected from 5 different leaflets, with 2 disks taken from each leaf. This procedure resulted in 10 replicates per treatment. Leaf disks were placed on agar-agar medium and incubated for 5 d at 25°C. *Pseudomonas syringae* pv. *tomato* DC3000 (*Pst*) was used as a positive control, and as a negative control, *Agrobacterium tumefaciens GV3101* (At). Photographs were taken 5 days after inoculation. Disease incidence on the leaf disks was evaluated as either positive (1) or negative (0).

### Assign taxonomy

Bacterial cultures were grown overnight in LB medium, followed by centrifugation at 5000 rpm for 10 min. A 20 µL aliquot of culture was mixed with 100 µL of PBS, incubated at 90°C for 10 min, and centrifuged at 10000 rpm for 1 min. The supernatants were transferred to a clean tube, resuspended in 200 µL of sterile Milli-Q water, and stored at −20 °C until use. The 16S rDNA region was amplified using the 27F (5′-AGAGTTTGATCCTGGCTCAG-3′) and 1492 R (5′-GGTTACCTTGTTACGACTT-3′). The PCR reaction mix comprised 40 µL containing 8 µL of buffer, 3.2 µL of MgCl₂, 0.8 µL of dNTPs, 0.5 µL of forward primer, 0.5 µL of reverse primer, 0.5 µL of GoTaq Flexi polymerase (Promega, Madison, WI), and 25 µL of nuclease-free water. Thermal cycling conditions were as follows: 5 min at 94°C, followed by 35 cycles of 94°C for 30 s, 55°C for 30 s, and 72°C for 60 s, and a final extension at 72°C for 7 min. PCR products were kept at 4°C until further use. PCR products were visualized by electrophoresis. PCR product sequencing was performed by Macrogen (Santiago, Chile). Consensus sequences were obtained using Bioedit v 7.7.1 (Hall, 1999). The sequences were aligned using the NCBI Genbank (https://www.ncbi.nlm.nih.gov/genbank/) database in the nucleotide blast tool, and the sequences were identified based on the similarity output results and taxonomy assignments. The sequences were aligned using MAFFT (https://mafft.cbrc.jp/alignment/server/) (Katoh et al., 2017), and edited in MEGA v 11.0.13 (Kumar et al., 2016). The phylogenetic tree was generated using the maximum likelihood tree method in the IQTree web server (http://iqtree.cibiv.univie.ac.at/) (Trifinopoulos et al., 2016), using the model with the best BIC score as determined by ModelFinder (Kalyaanamoorthy et al., 2017). The tree was rooted with *Thermosulfobacterium geofontis* (NR 118457.1) and visualized and edited with iTOL v.7 (https://itol.embl.de/), incorporating PGPR profile metadata using binary data (presence/absence).

### Bacterial performance

Ten bacterial strains were selected to evaluate pathogen resistance and plant growth-promoting rhizobacteria (PGPR) activity (Table xx). Bacteria were cultured for 24 h at 28°C in a TSB medium with agitation and adjusted to an optical density (OD_600_) of 0.2 for subsequent assays. Two-week-old *Solanum lycopersicum* v. "Alamina" seedlings were root-dipped in the corresponding bacterial suspension for 30 s, transplanted into individual pots containing a 1:1 peat-to-sand substrate, and maintained in a growth chamber under controlled conditions (25°C, 16:8 h light/dark photoperiod) throughout the experiment. Three days post-transplantation, seedlings were reinoculated by drenching with 10 mL of the same bacterial suspension (Yu and Macho, 2021), while mock-treated plants and *Agrobacterium tumefaciens* served as negative controls, with irrigation maintained at 10 mL of water every two days. To assess both induced immunity and root development, each bacterial treatment group (n = 7 per group) was divided into two subgroups, where half of the plants were challenged with *Pseudomonas syringae* pv. *tomato* (*Pst*) DC3000, is the causative agent of the bacterial speck disease (Morris et al., 2008), two weeks post-inoculation—when priming was expected to be activated—to evaluate pathogen resistance, while the remaining plants continued growing under the same conditions to assess root development. *Pst* DC3000 was grown for 48 h in TSB medium, and seedlings were sprayed with a *Pst* suspension at 2 × 10⁸ CFU mL⁻¹ until runoff, followed by polyethylene bag coverage for 24 h to maintain high humidity (Basim et al., 2004), with sterile distilled water as a negative control. Disease symptoms, including water-soaked lesions and dark brown-to-black spots, were monitored, and seven days post-inoculation, six-leaf disks from two symptomatic leaflets per plant (1 g of tissue) were suspended in PSB buffer, homogenized using D1000 hand-held (Sigma-Aldrich), and serially diluted (10⁻⁹ and 10⁻¹⁰) before plating on TSA supplemented with rifampicin, as *Pst* DC3000 is rifampicin-resistant (Xin and He, 2013). The bacterial population per gram of fresh leaf tissue was quantified after 24 h using a stereoscopic microscope (Motic Smz-171). Seedlings were maintained under identical conditions for one month, and root system development was evaluated in both *Pst*-challenged and non-inoculated plants to determine whether bacterial treatments influenced root architecture.

Before analysis, plants were carefully removed from the substrate, and roots were thoroughly rinsed with tap water to eliminate residual soil particles. Root morphology was assessed using a high-resolution flatbed scanner (Epson Perfection V800 Photo & V850 Pro, Seiko Epson Corp., Japan; resolution: 6400 dpi), and key root parameters—including root length (RL), projection area (PA), surface area (SA), mean root diameter (MRD), root volume (RV), and root tip number (RT)—were quantified using WinRhizo 2019a software (Regent Instruments Inc., Canada) (Contreras-Soto et al., 2022). This experimental setup ensured that all plants were exposed to identical growth conditions, with the only variable being bacterial inoculation treatments, and subsequent *Pst*-challenge in a subset of plants. For the statistical analysis of the Pst-inoculation assay, ten bacterial treatments, along with water and *Agrobacterium tumefaciens* controls, were inoculated with *Pst* to assess both disease progression (via CFU per gram of leaf tissue) and root development, comparing induced immunity and pathogen control. For the PGPR assay, the bacterial treatments were compared to the water control (mock) to evaluate their impact on root development. In both assays, the Kruskal-Wallis test was used to assess overall differences between treatments, followed by Wilcoxon pairwise comparisons to determine significant differences relative to the controls. Additionally, K-means clustering was used to classify bacterial isolates based on their effects on root growth, identifying distinct response profiles. The optimal number of clusters was determined using the elbow method. A scatterplot was generated to visualize the relationship between root variables and the bacterial genera of the isolates. This approach allowed for an evaluation of bacterial strain effects on pathogen resistance, immune activation, and root growth promotion

## Supporting information

Supplementary images and tables

