## Supplementary images and tables for "Bioprospection of culturable soil-borne bacteria with biotechnological potential for use in priming defense"

**Supplementary Material**

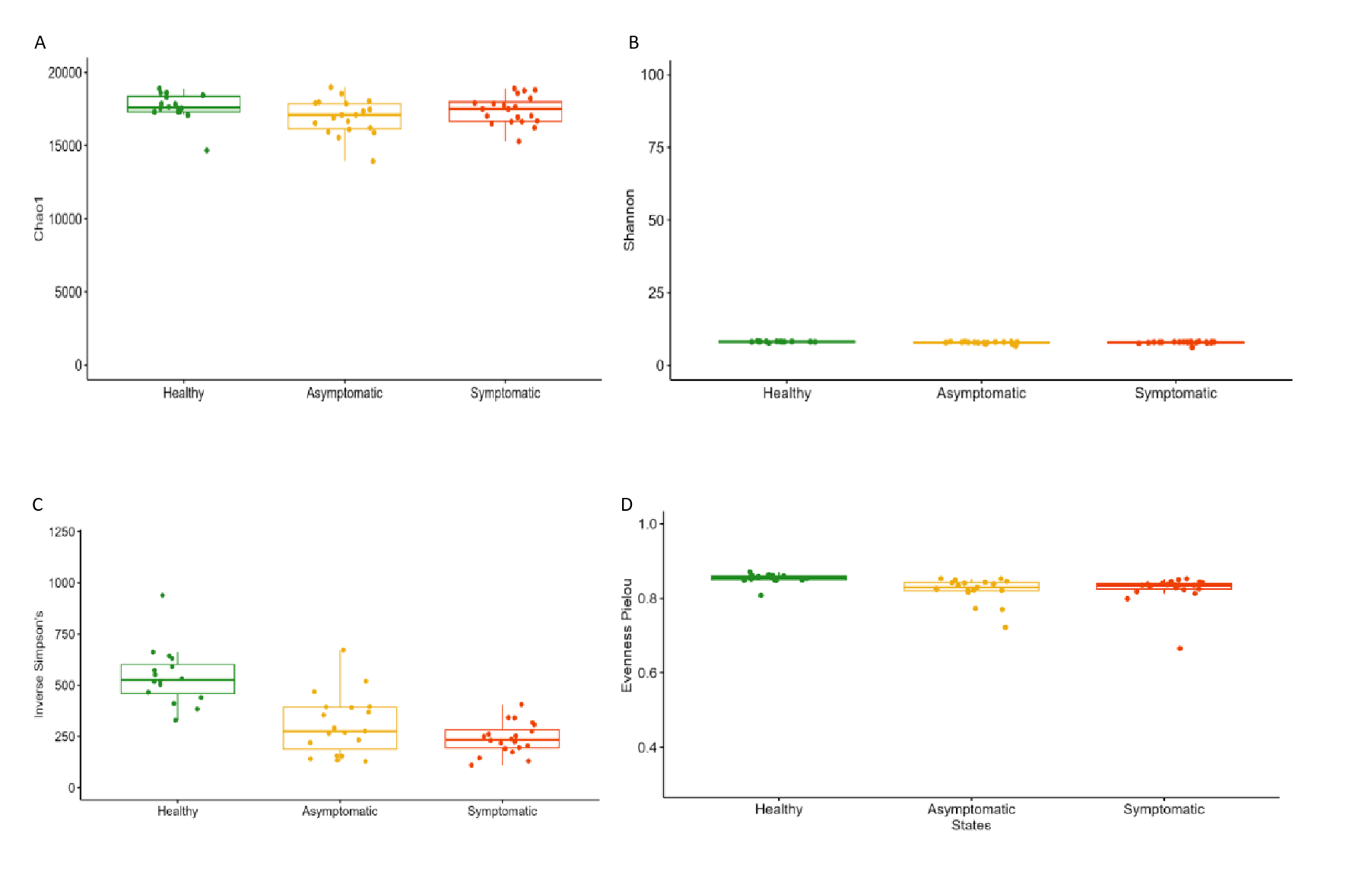
**Figure S1:** Diversity indices were estimated for the phytopathogenic status of tomato plants. Chao 1 diversity index (a); Shannon (b); Inverse Simpson index (c), and Pielou’s evenness index (d) estimated bacterial communities. Boxplot color represents phytopathogenic status.

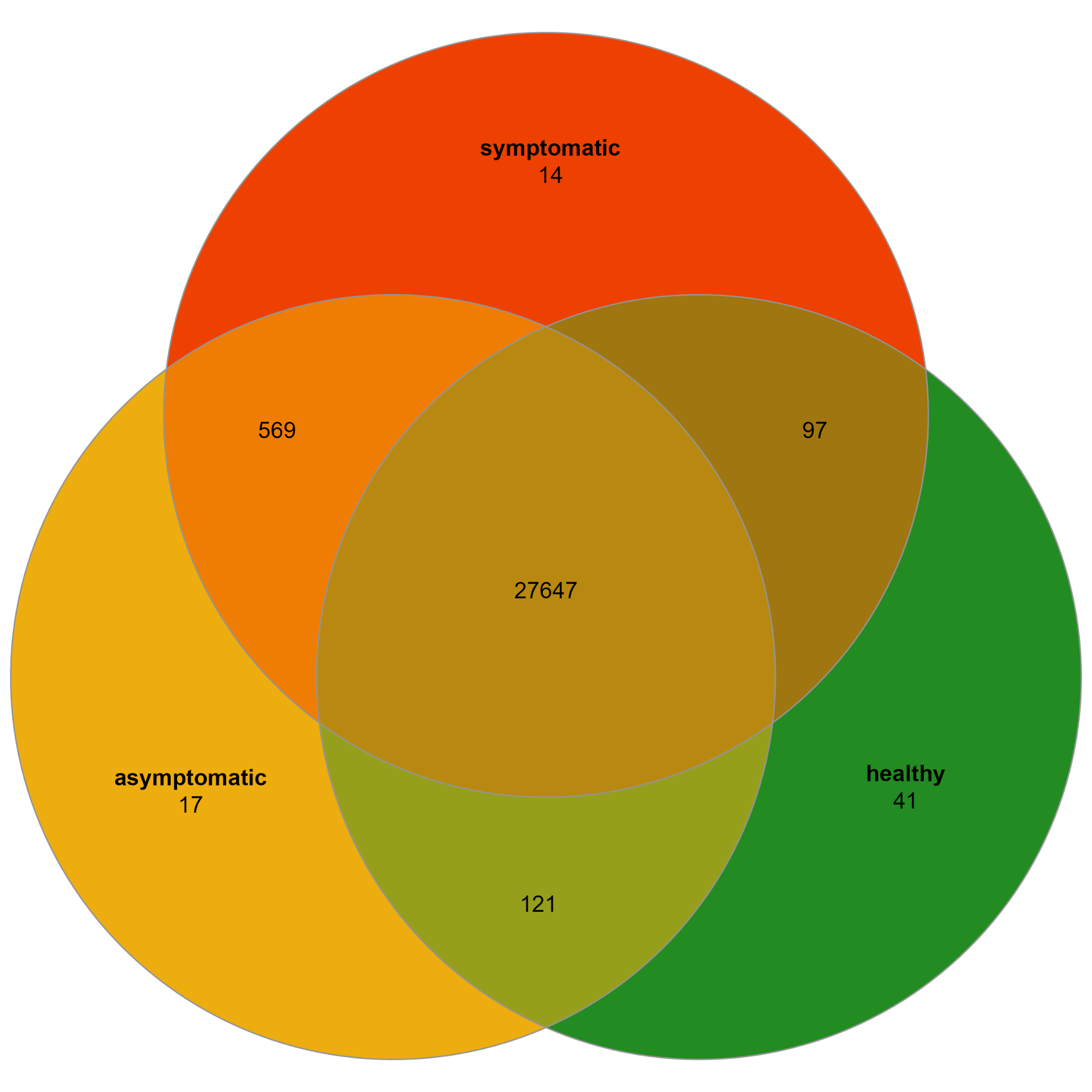

**Figure S2:** Venn diagram based on ASVs composition of bacterial communities associated with the phytosanitary status of tomato plants

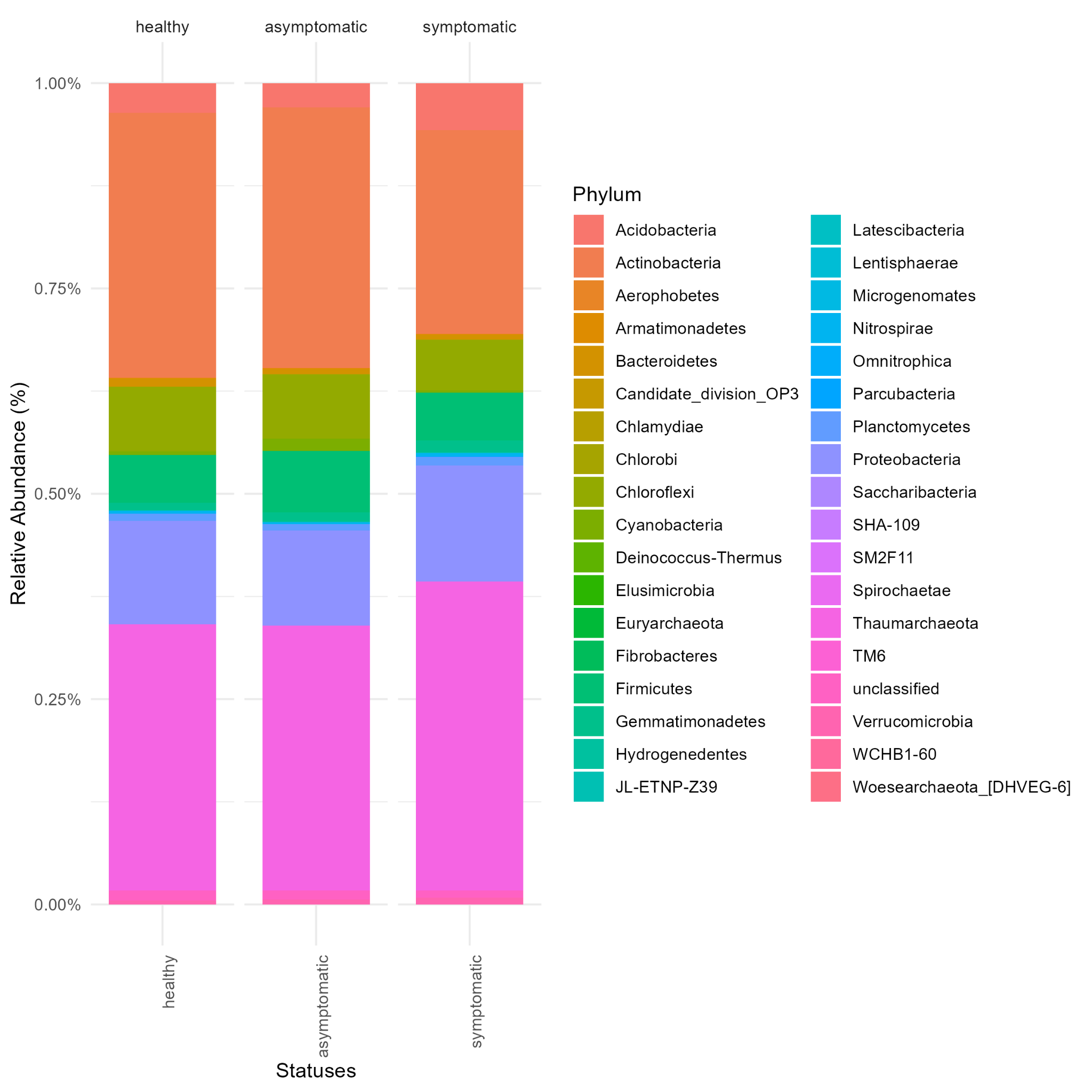

**Figure S3:** Barplot of relative abundance of the phyla based on bacterial communities associated with the phytosanitary status of tomato plants

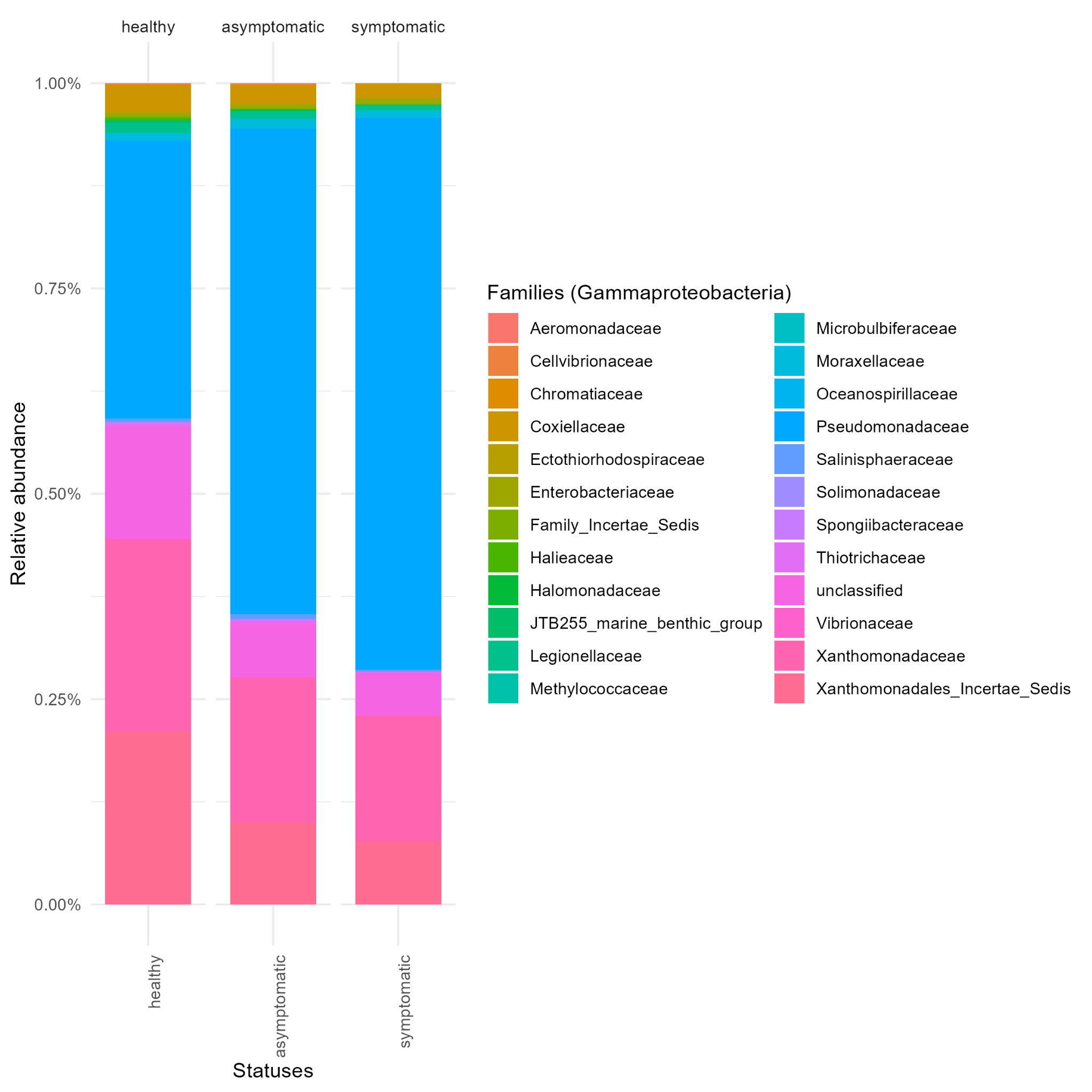

**Figure S4:** Barplot of relative abundance of the families in Gammaproteobacteria class based on bacterial communities associated with the phytosanitary status of tomato plants

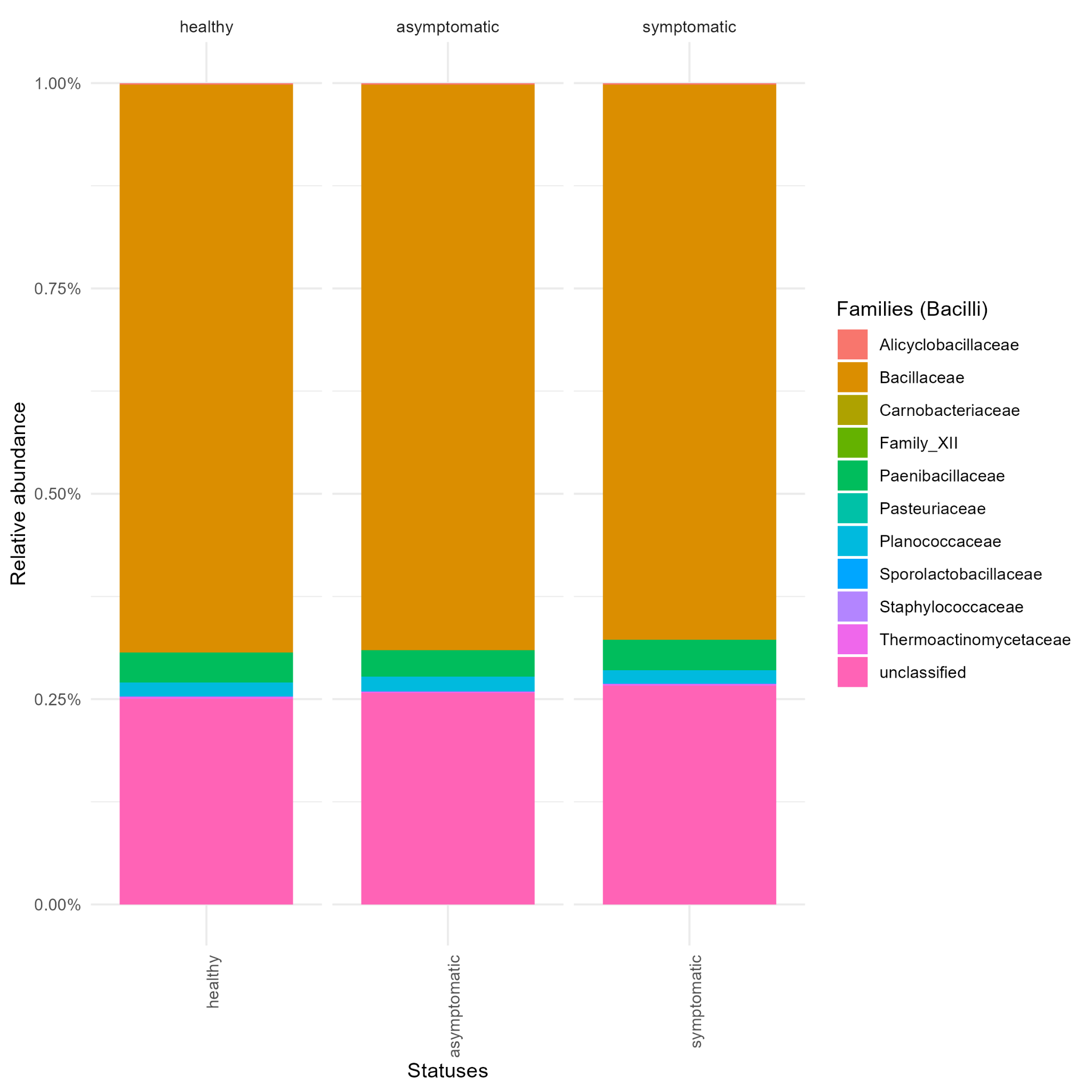

**Figure S5:** Barplot of relative abundance of the families in the Bacilli class based on bacterial communities associated with the phytosanitary status of tomato plants

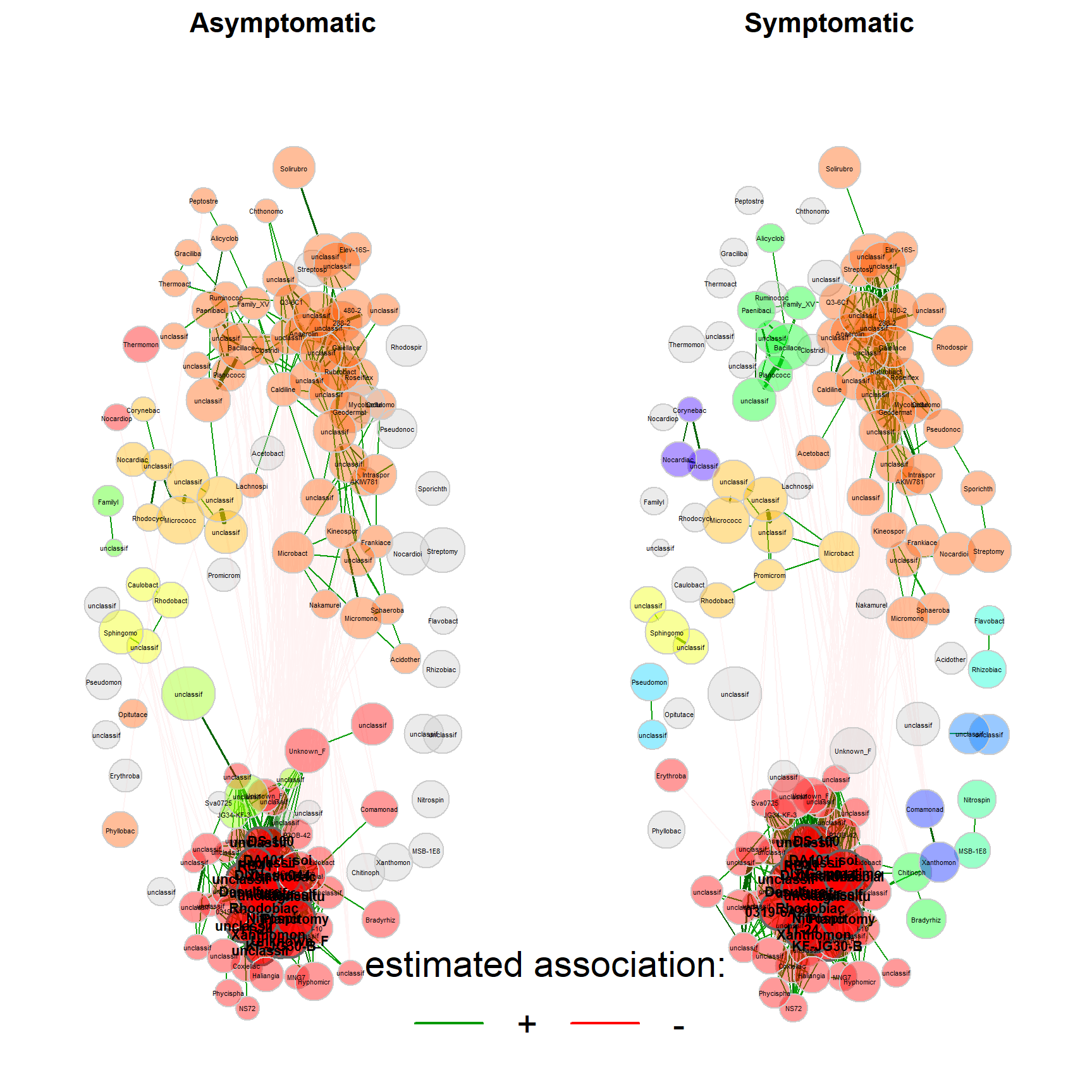

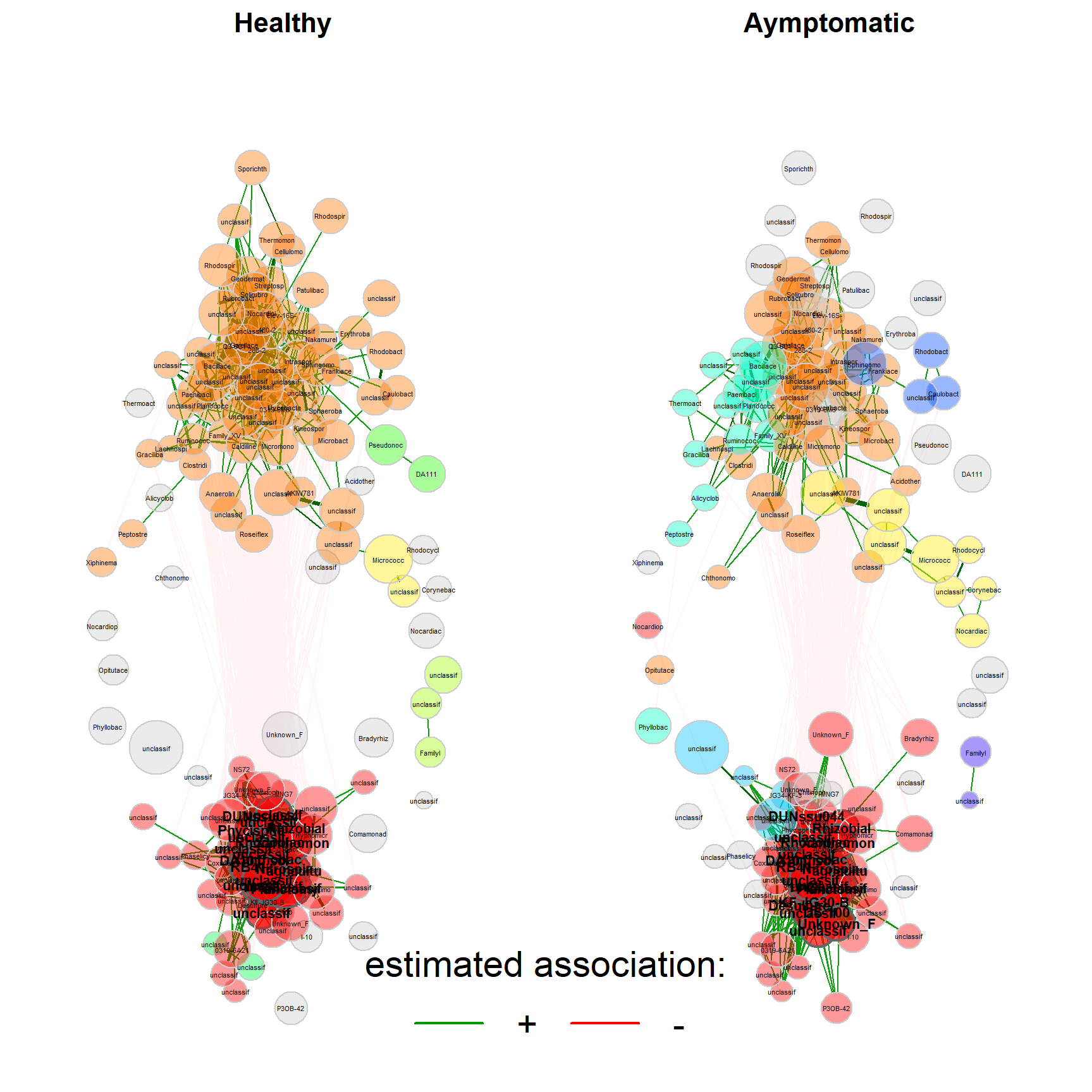

**Random forest**

**Figure S6:** Network of dissimilarities between families of bacterial communities associated with the phytosanitary status of tomato plants. Positive (green) and negative (red) correlation based on the SparCC algorithm with a threshold of absolute association value of 0.85 between taxa.

**PGPR profile**

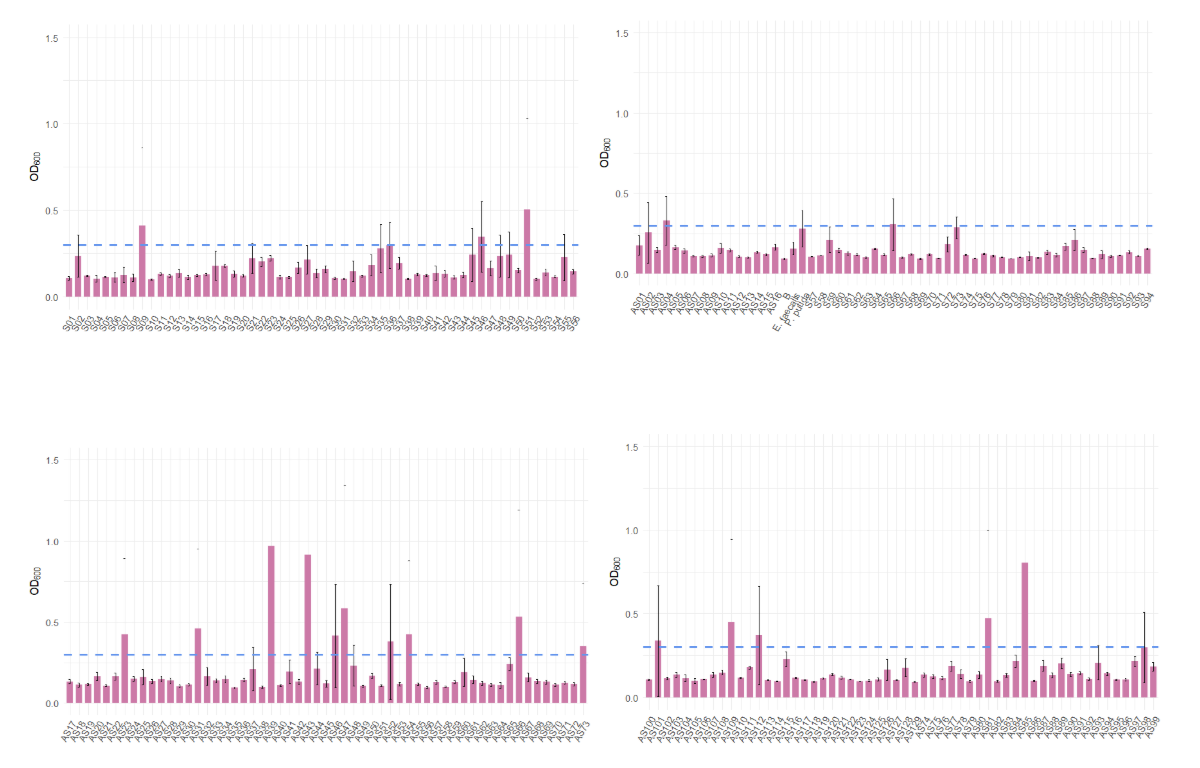

**Figure S6:** Barplot showing indole-3-acetic acid (IAA) production by bacterial strains isolated from soils associated with healthy (S) and asymptomatic (As) tomato plants. The blue line indicates the threshold absorbance (OD₅₃₅) corresponding to the reference strain *Pseudomonas putida* ATCC 12633*. Enterococcus faecalis* was included as a negative control.

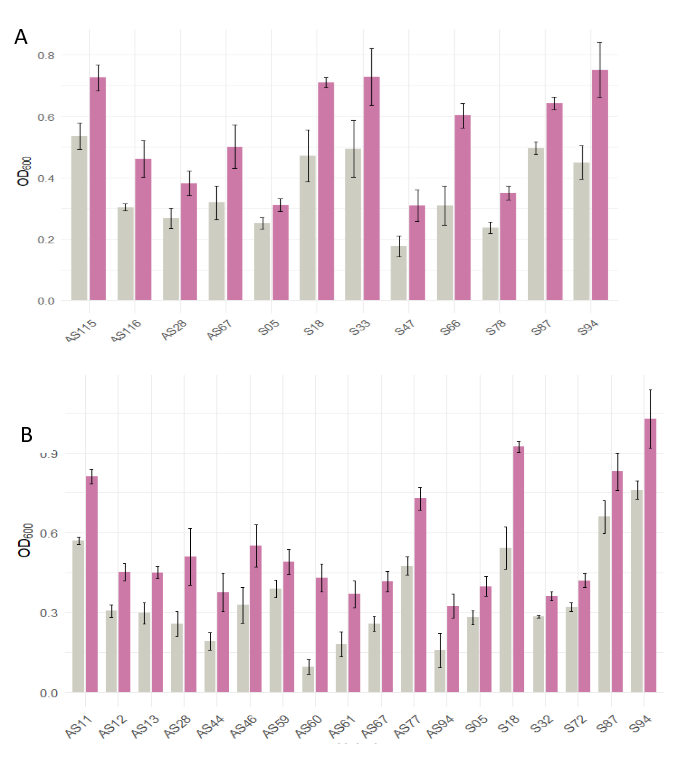

**Figure S7:** Barplot showing 1-aminocyclopropane-1-carboxylic acid (ACC) deaminase activity by bacterial strains isolated from soils associated with healthy (S) and asymptomatic (As) tomato plants. A: measurement at 48 hours; B: measurement at 96 hours. Gray bars represent growth in DF medium without ACC (negative control), while purple bars indicate growth in DF medium supplemented with ACC.

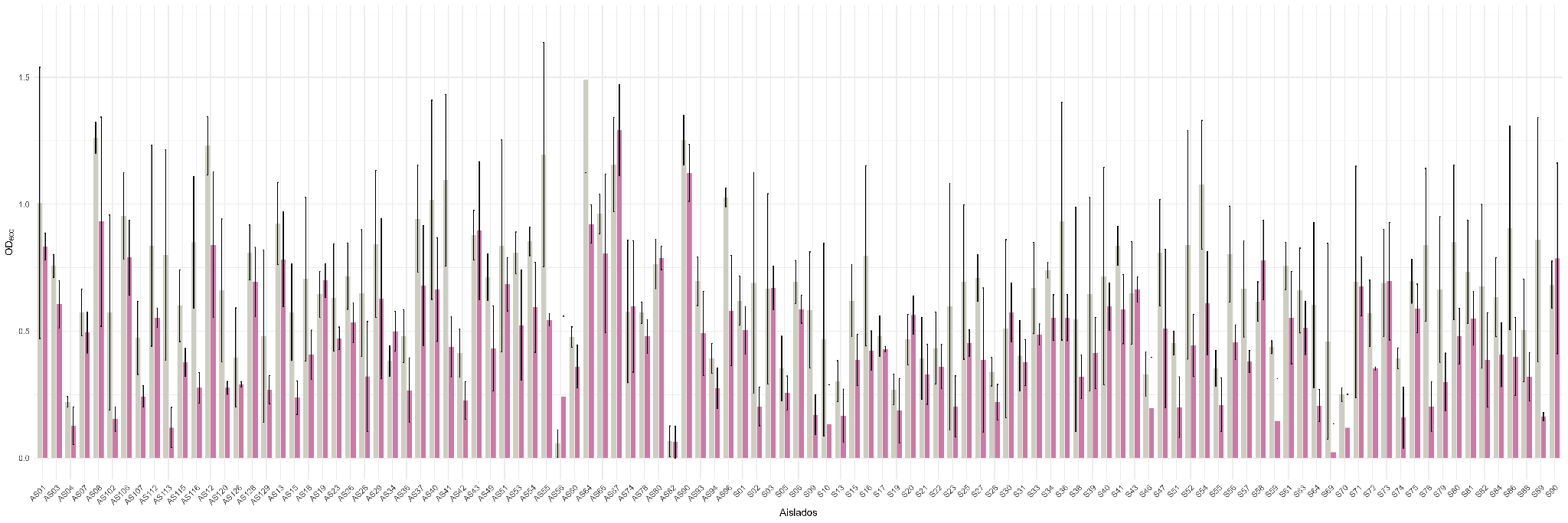

**Figure S8:** Barplot showing osmotic stress resistance of bacterial strains induced by PEG6000*.* Bacterial growth (OD600) at 96 hours is shown for different strains under non-stress (grey bars) and osmotic stress conditions (purple bars). Bars represent the mean ± standard error. Strains that did not exhibit significant growth differences between control and stress conditions were considered tolerant to osmotic stress.

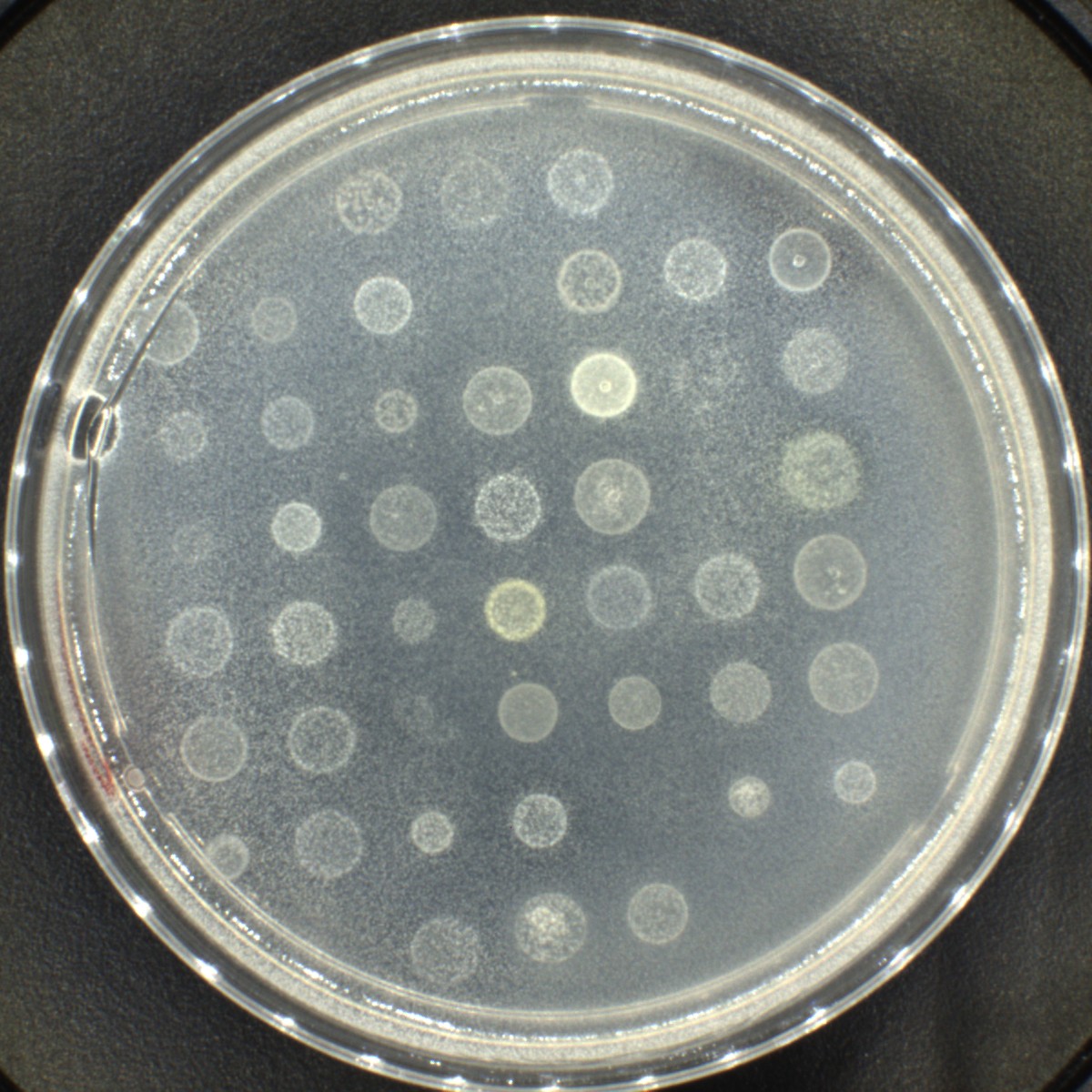

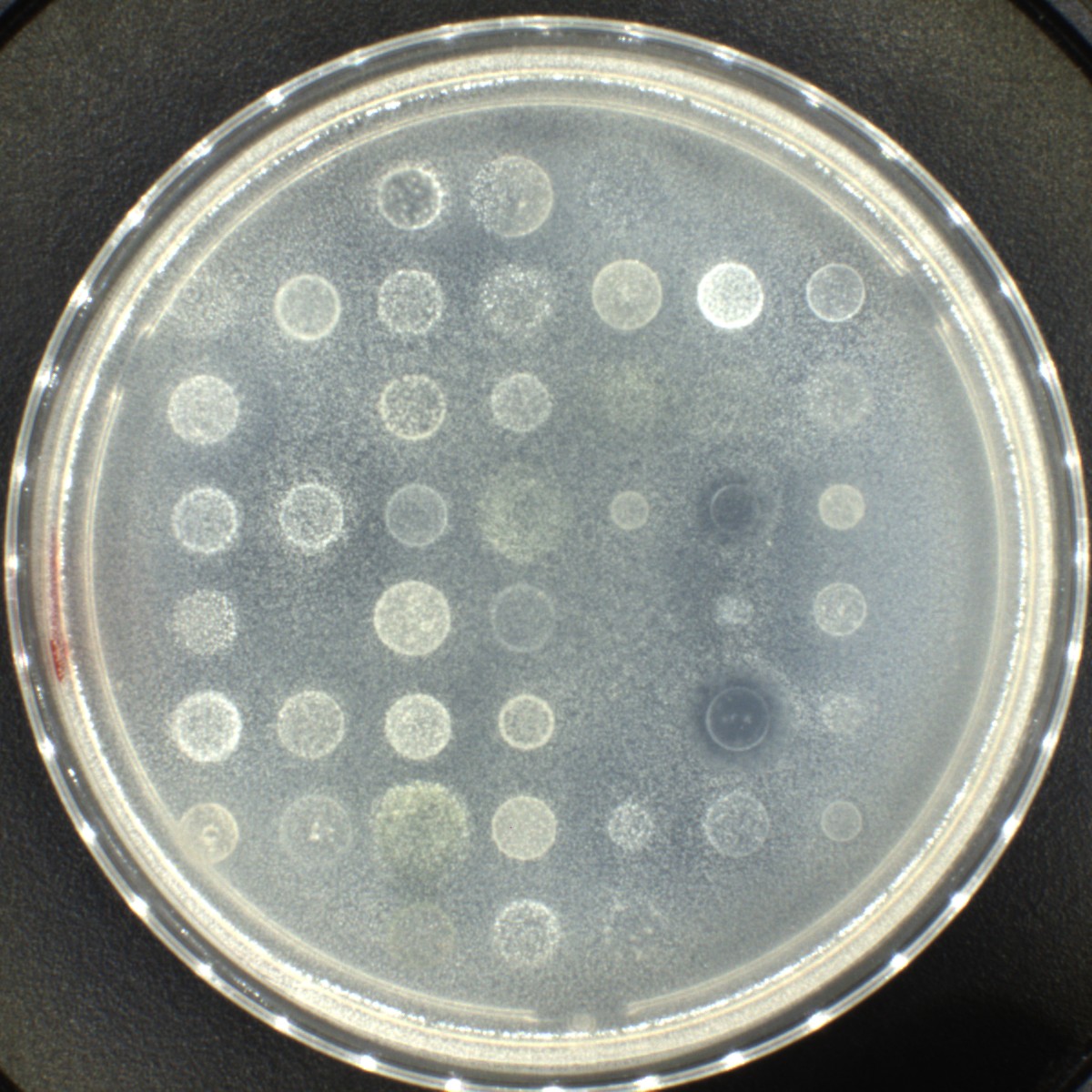

**Figure S9:** Nitrogen fixation capacity of bacterial strains evaluated on Norris Glucose Nitrogen-Free medium. Halo formation indicates potential nitrogen-fixing activity. Assessment performed after 48 hours of incubation.

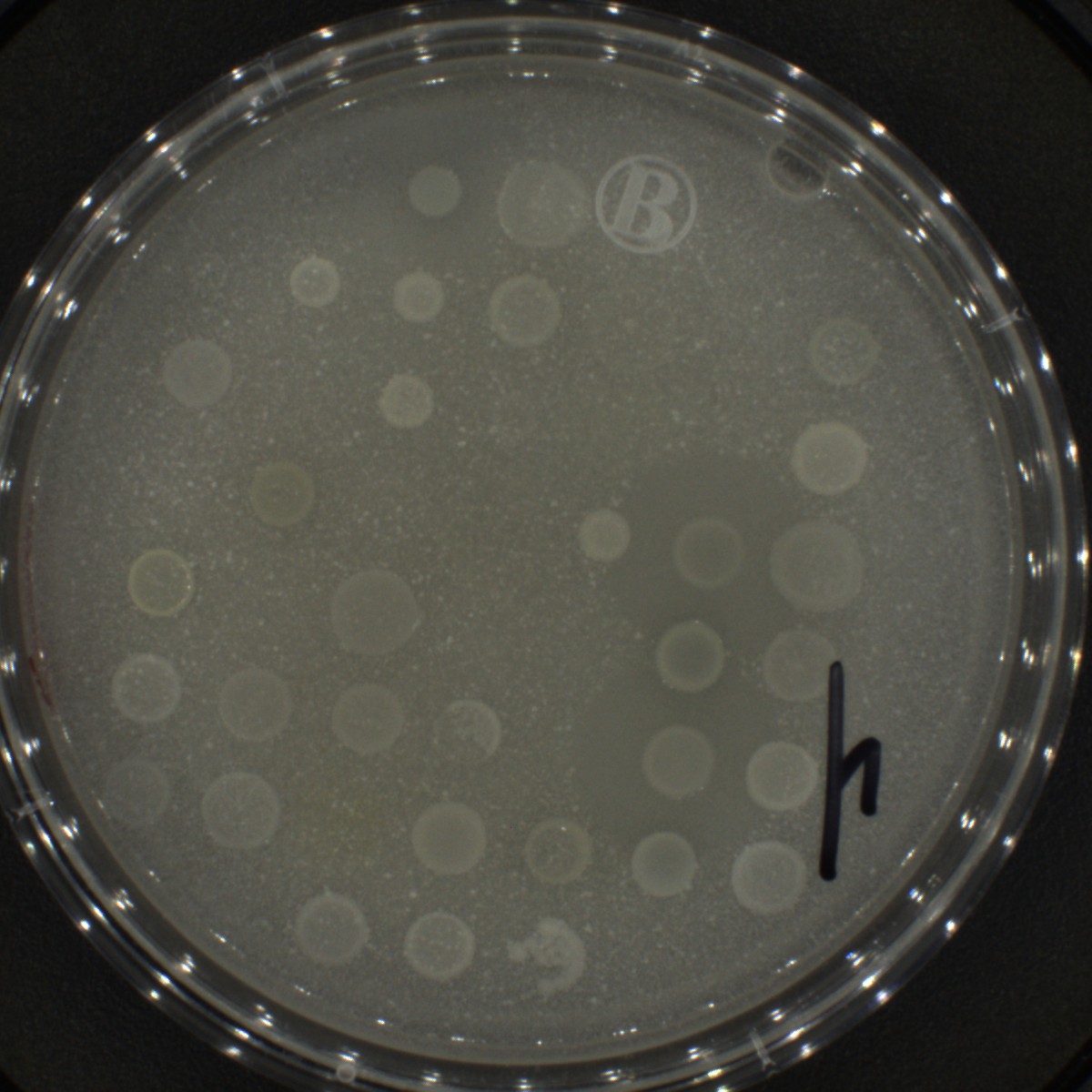

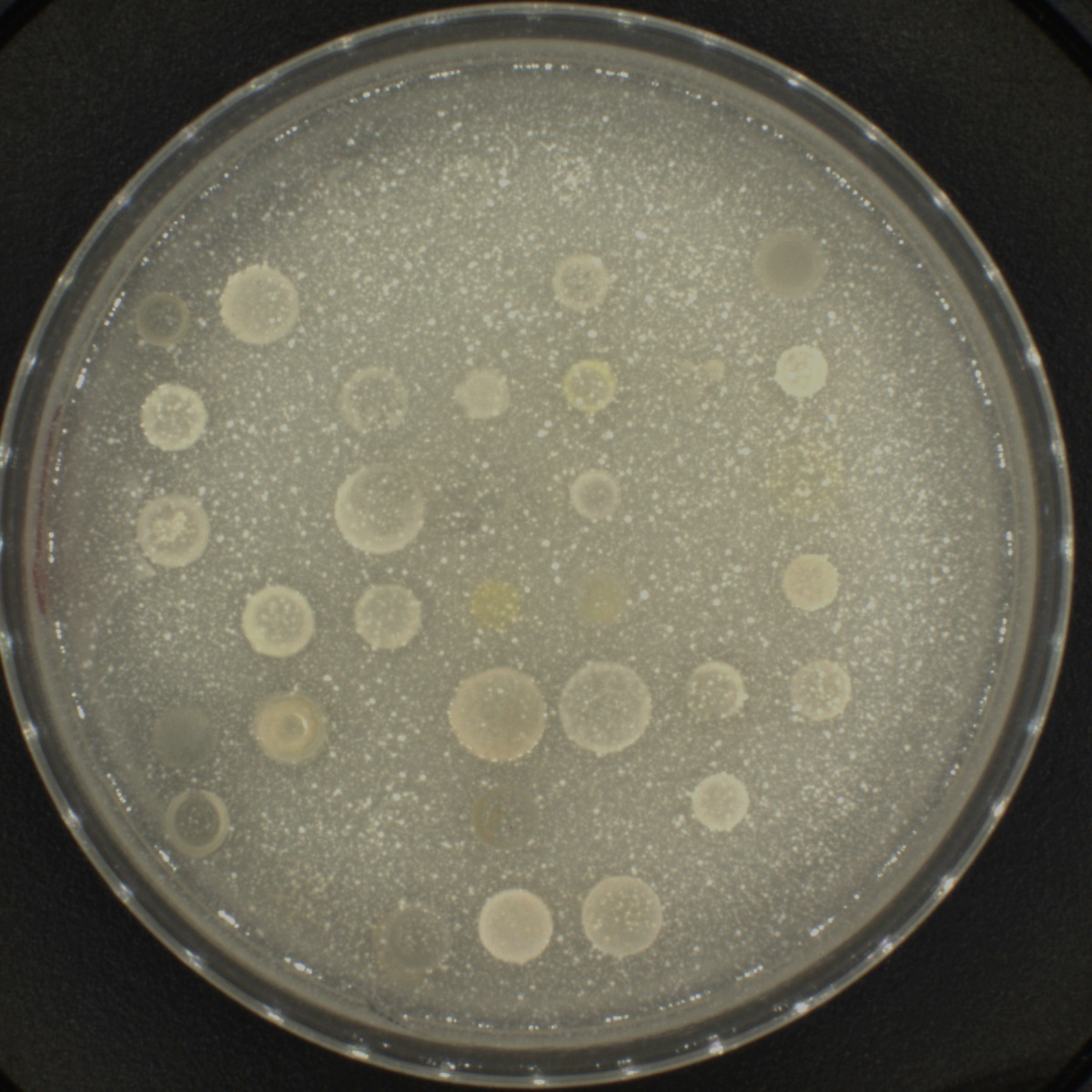

**Fig S10.** The phosphate solubilization capacity of bacterial strains was evaluated using Pikovskaya’s medium. Halo formation indicates the ability to solubilize inorganic or organic insoluble phosphorus into bioavailable forms. Assessment was conducted after 48 hours of incubation.

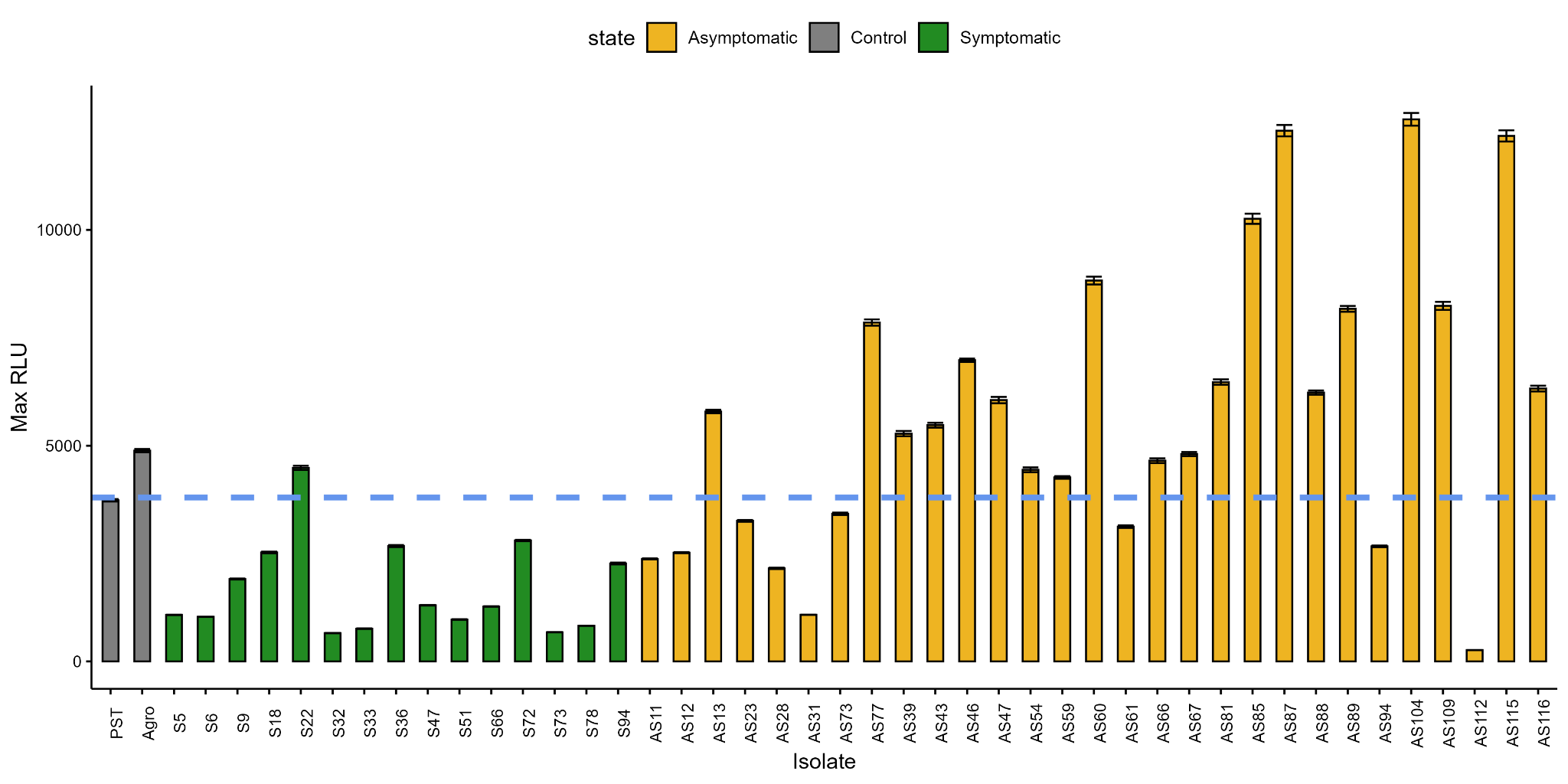

**Fig S11:** Maximum Relative Light Units (Max RLU) of bacterial strains isolated from soils associated with healthy (S) and asymptomatic (As) tomato plants. Reactive oxygen species production was monitored every 4 seconds over 18 cycles using the HRP-luminol chemiluminescence assay. *Pseudomonas syringae* pv. *tomato* (PST) was used as a positive control. The blue line represents the Max RLU threshold for PST. Values are expressed as mean ± SEM from six technical replicates.

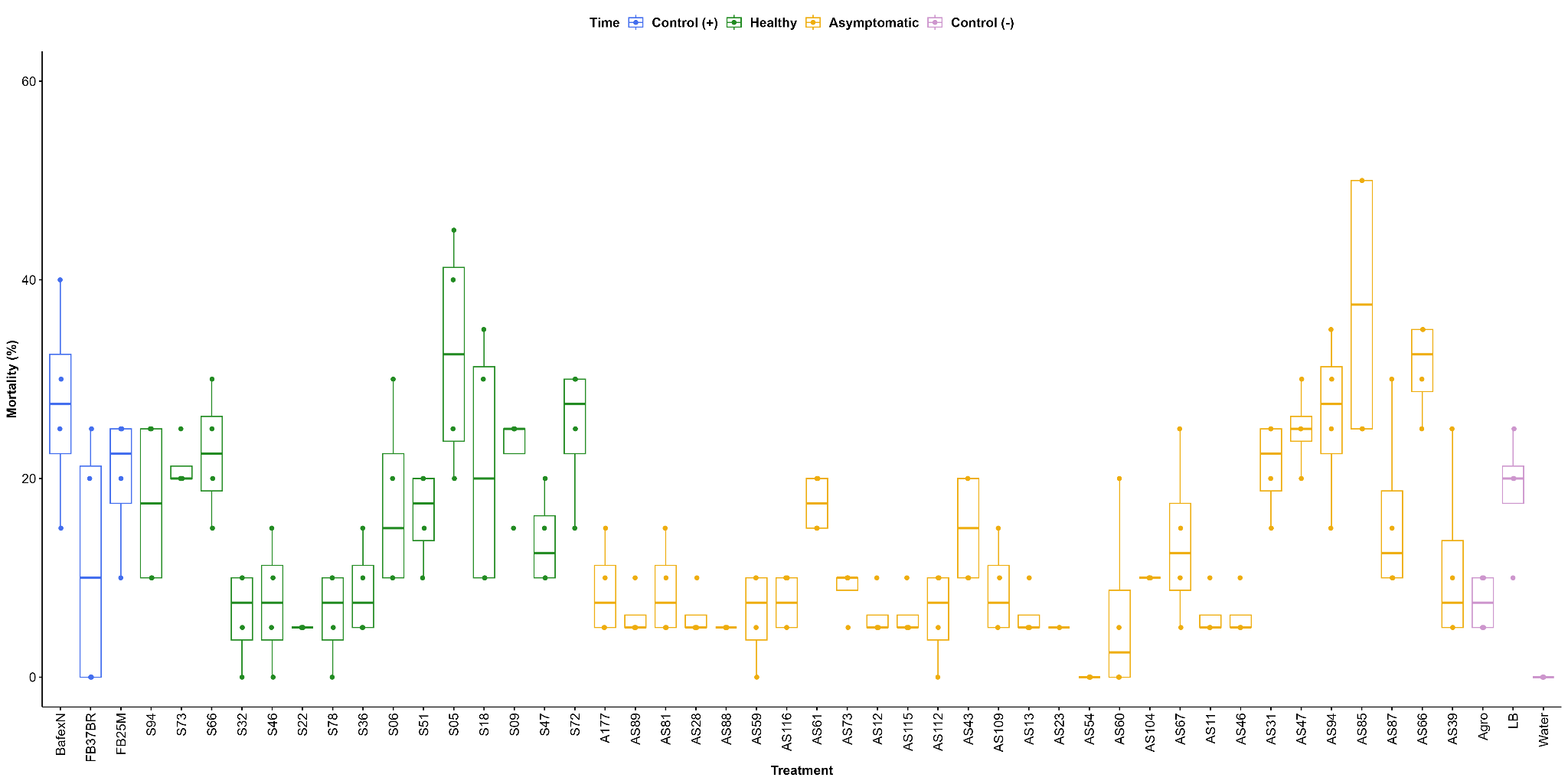

**Fig S12**: Barplot showing the percentage of mortality(%) of second-stage juveniles (J2) of *Meloidogyne* spp*.* by bacterial strains isolated from soils associated with healthy (S) and asymptomatic (As) tomato plants*. Agrobacterium tumefaciens (agro),* LB medium, and distilled water were employed as negative controls, and two *Bacillus* spp. isolates with known nematicide activity (FB25M and FB37BR), and the commercial biological nematicide BAFEX-N® as a positive control. The assay was evaluated 72 h before the application of treatments. Each treatment was evaluated in four independent replicates.

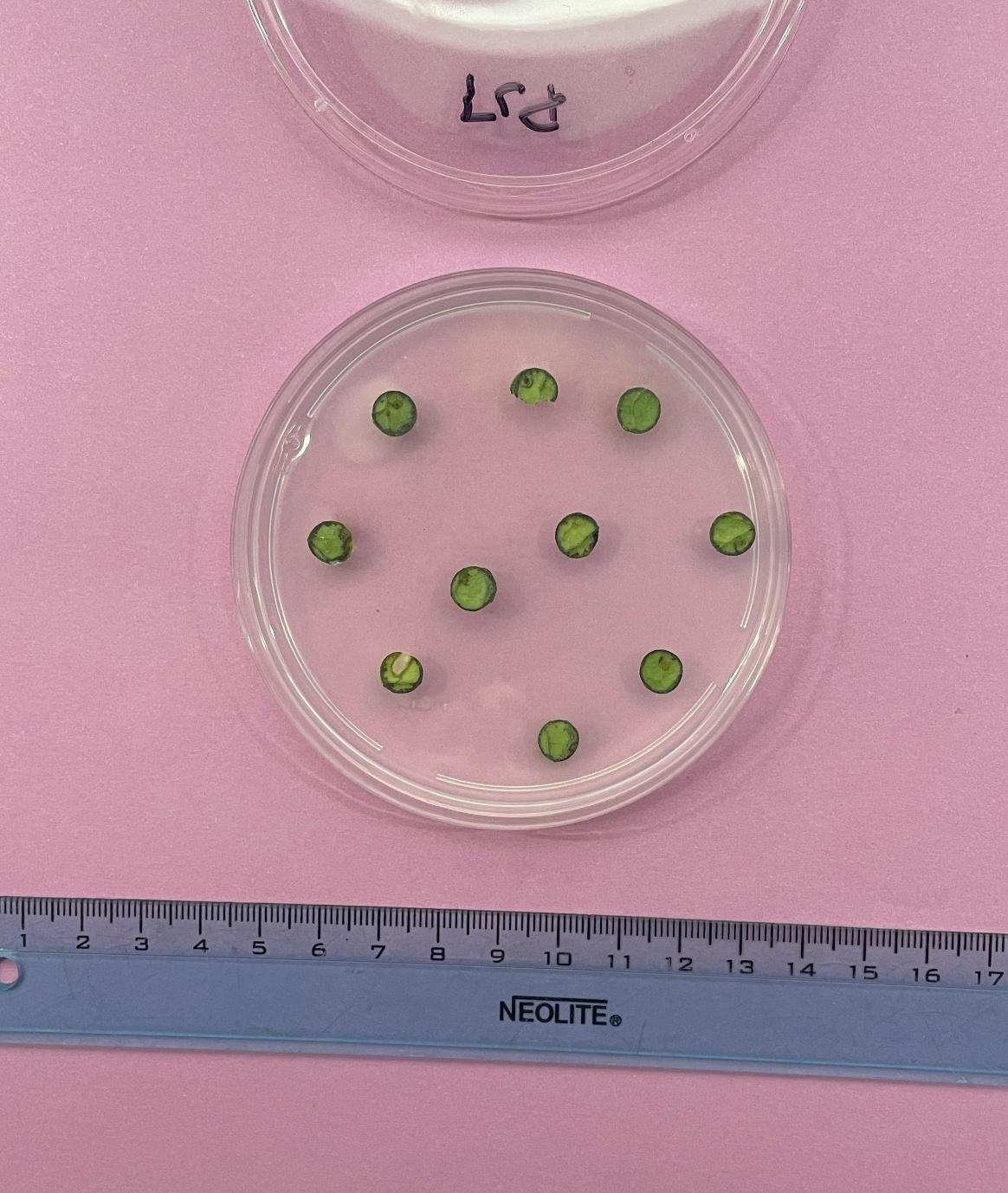

**PST – control (+)**

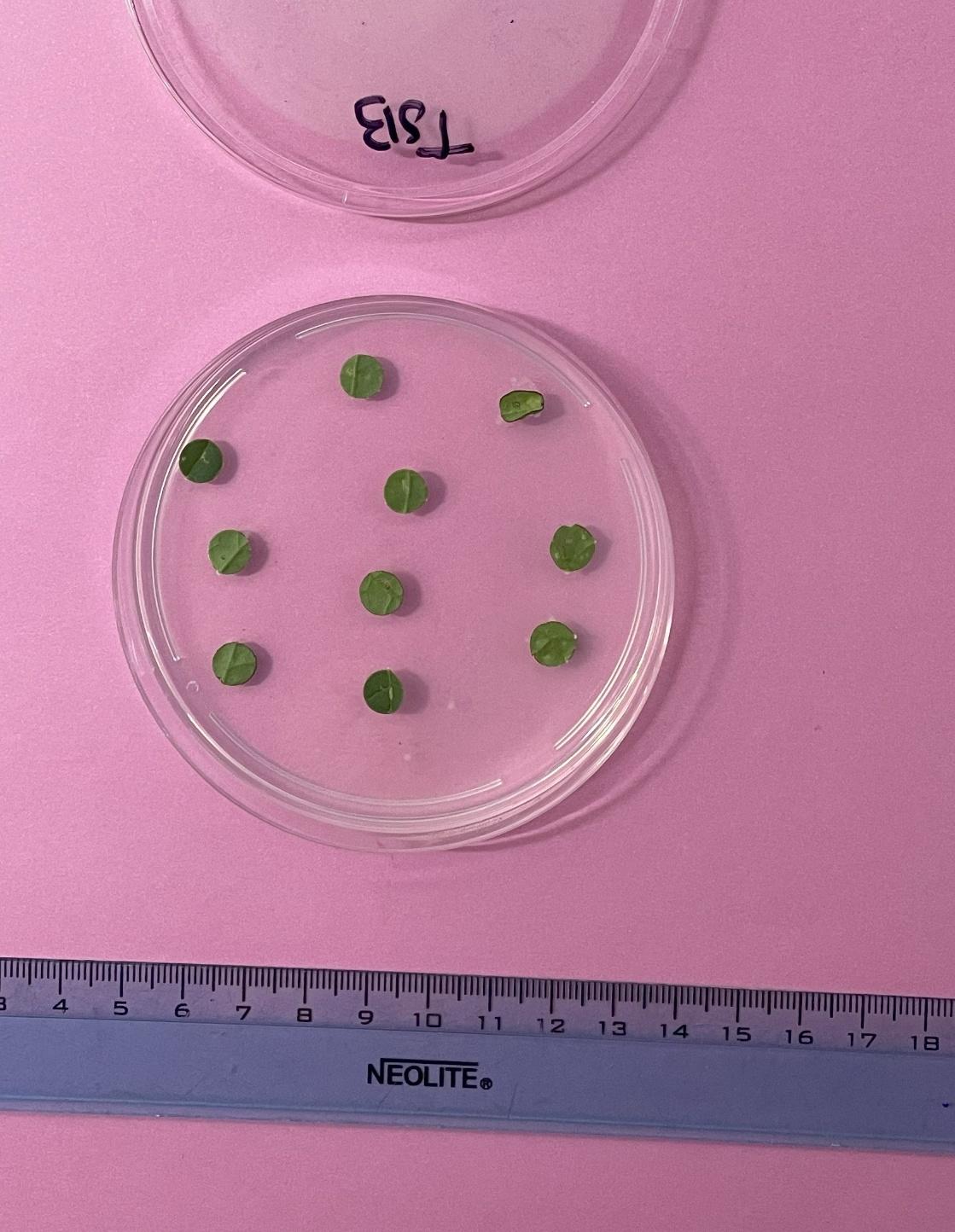

**TSB – control (-)**

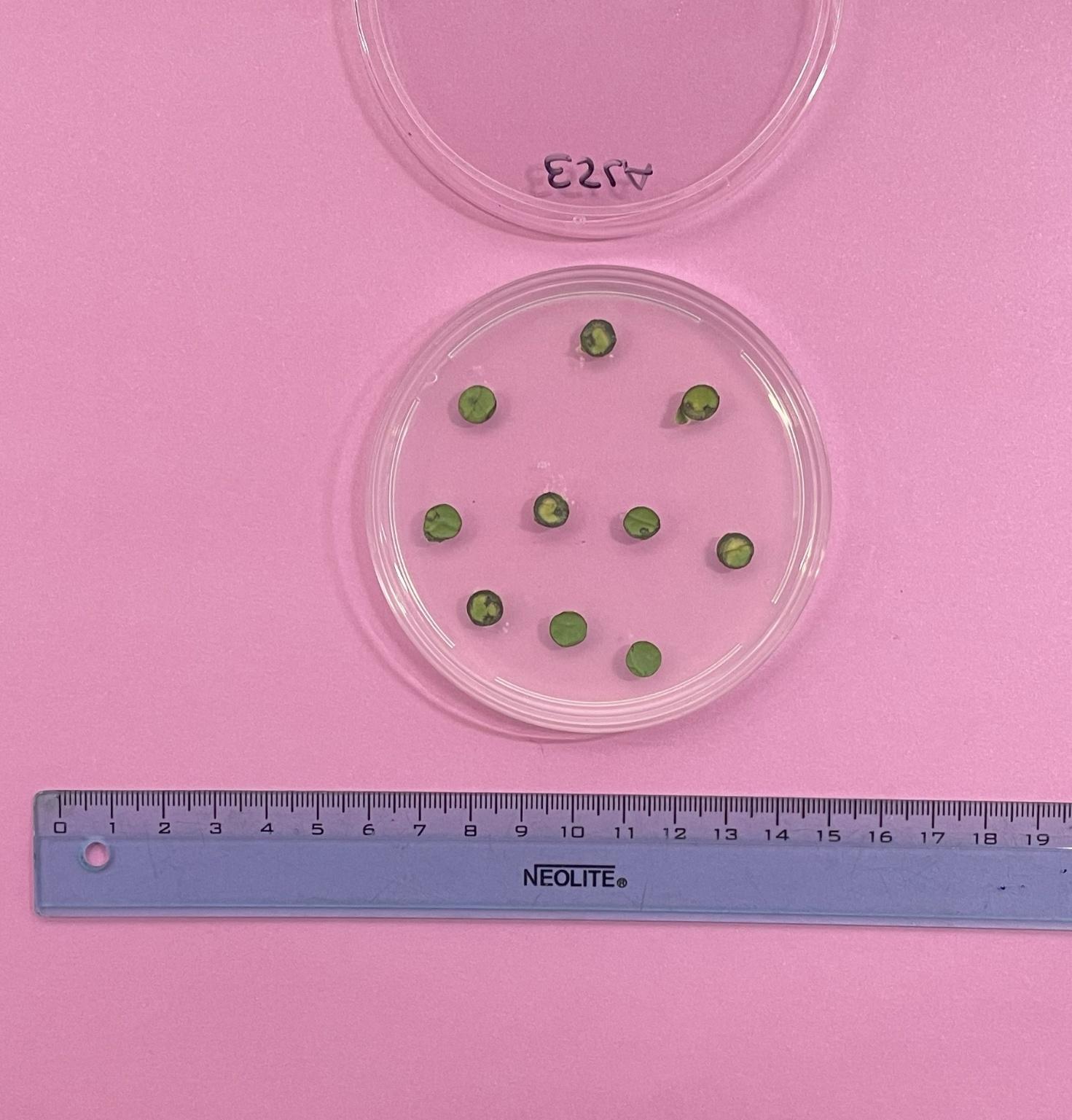

**AS23**

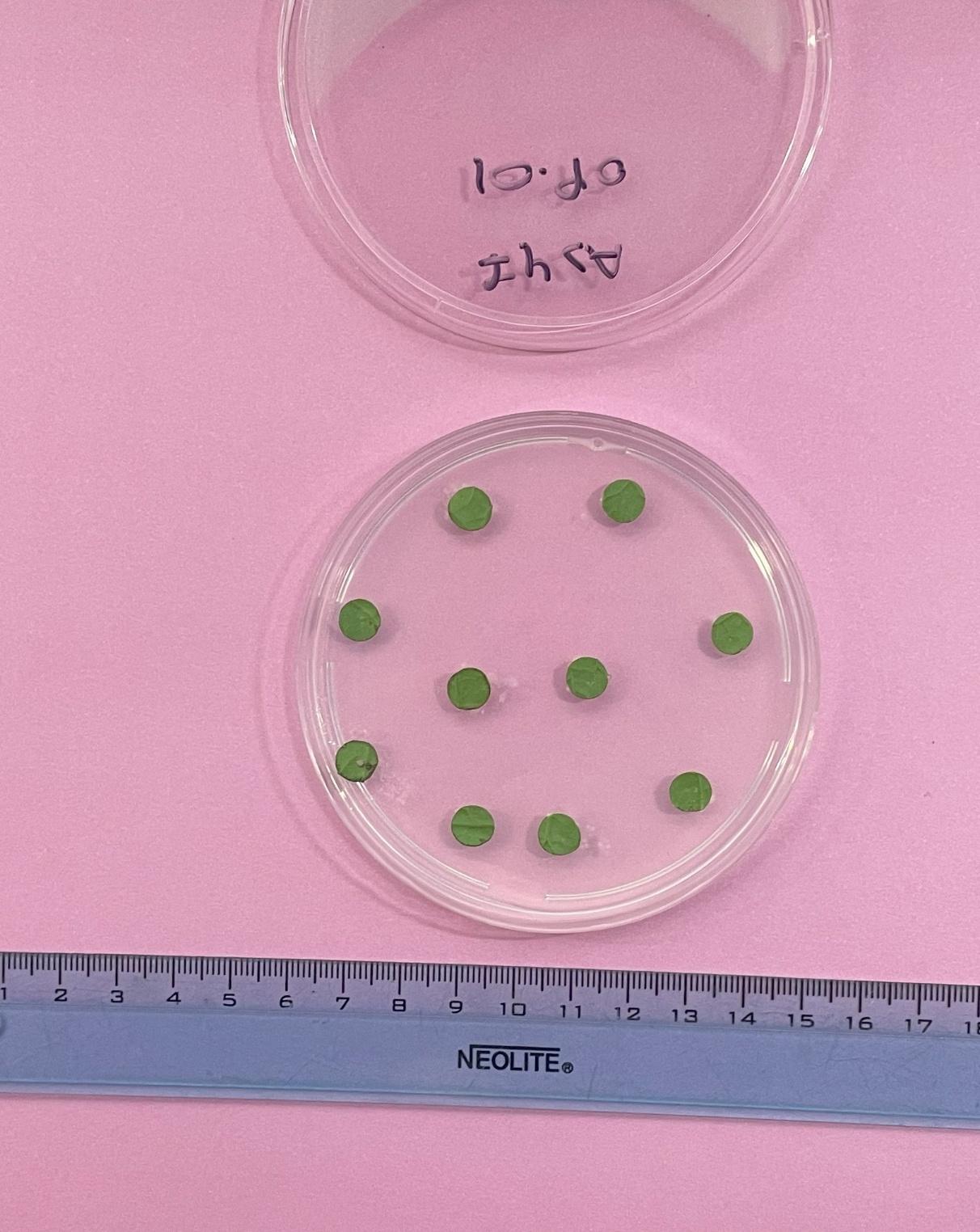

**AS47**

**Fig S13.** Example of a pathogenicity assay based on the presence or absence of symptoms. The assay included *Pseudomonas syringae* pv. *tomato* as a positive control and TSB medium as a negative control.

**PGPR assay**

**Bioestimulant activity**

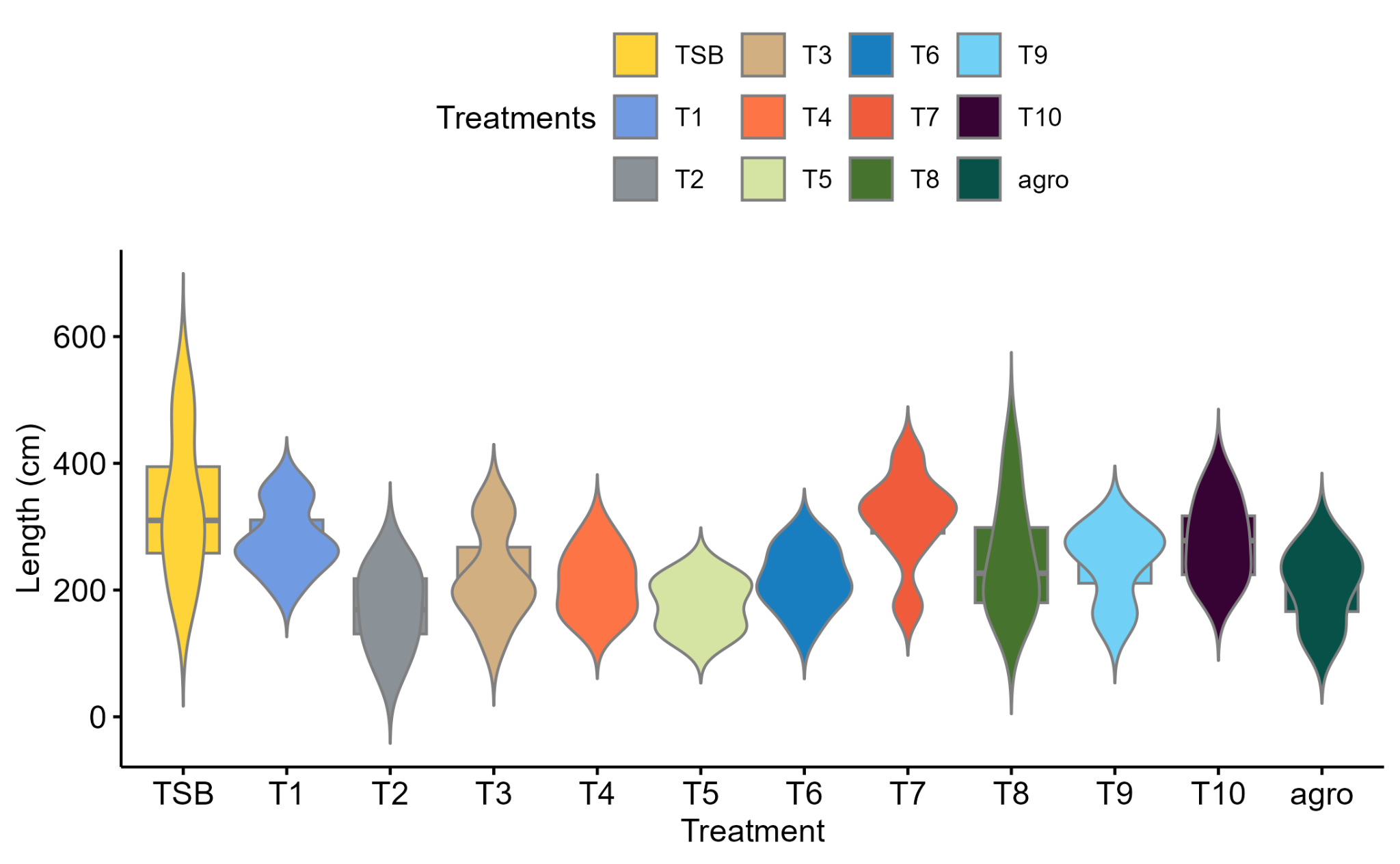

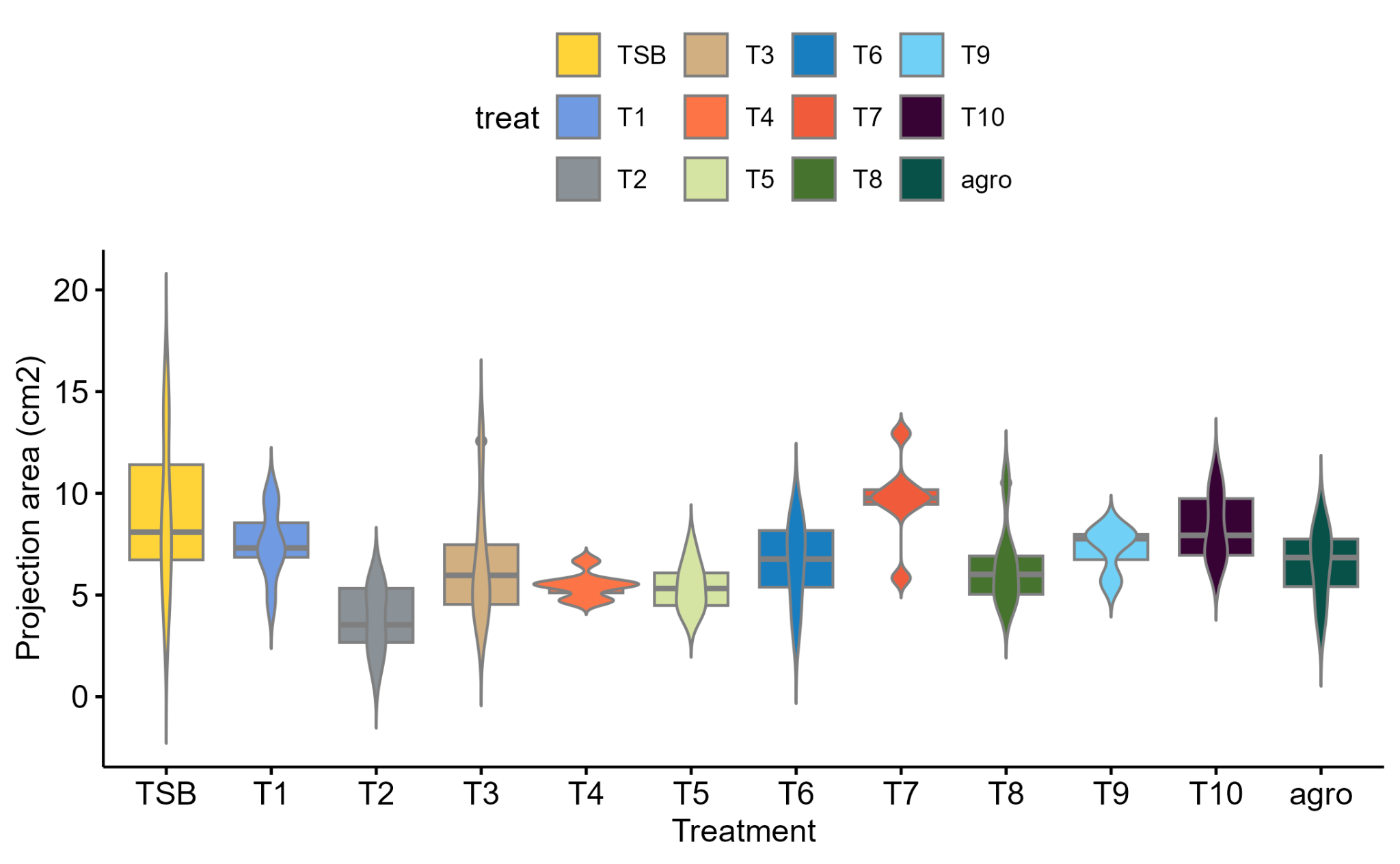

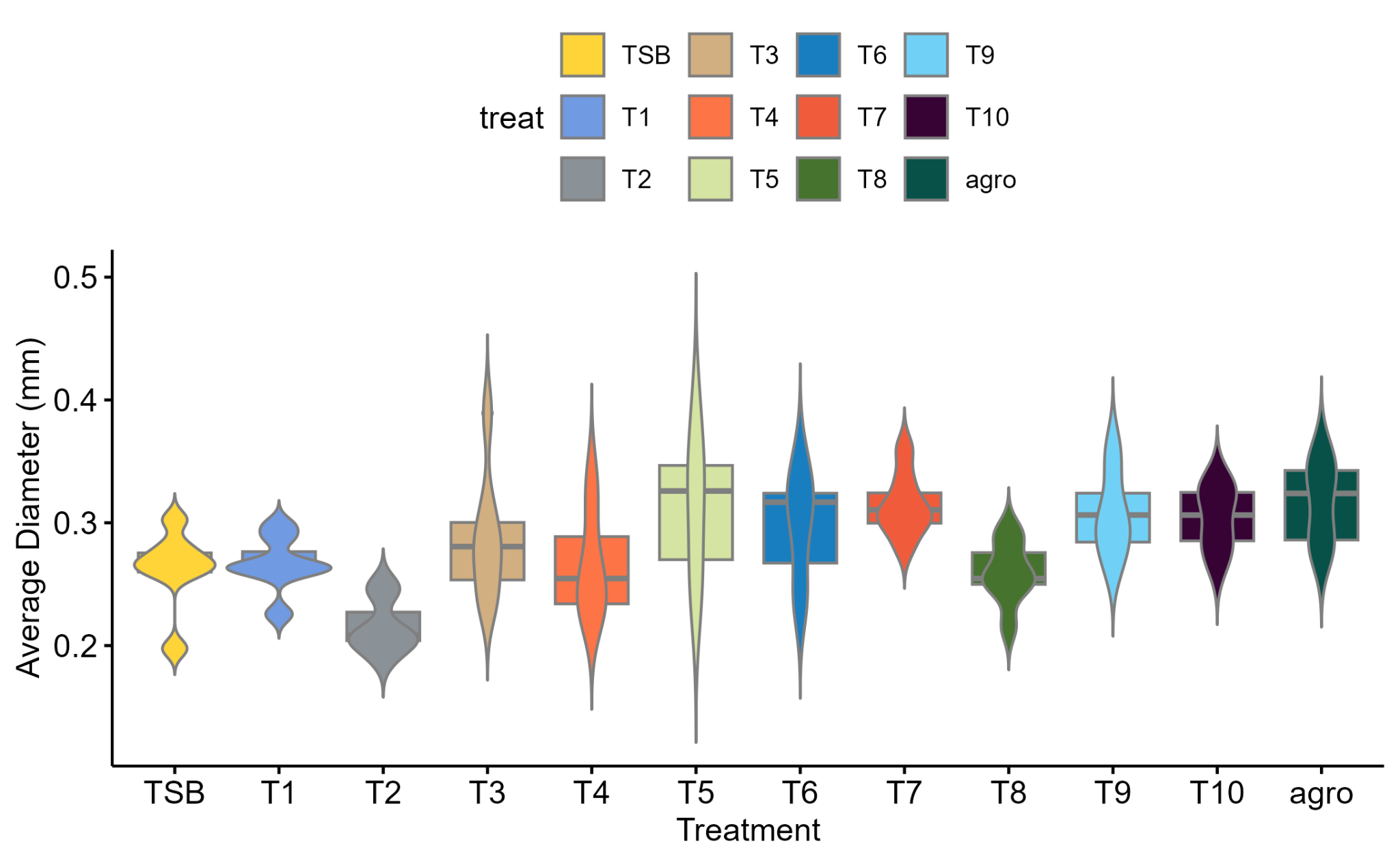

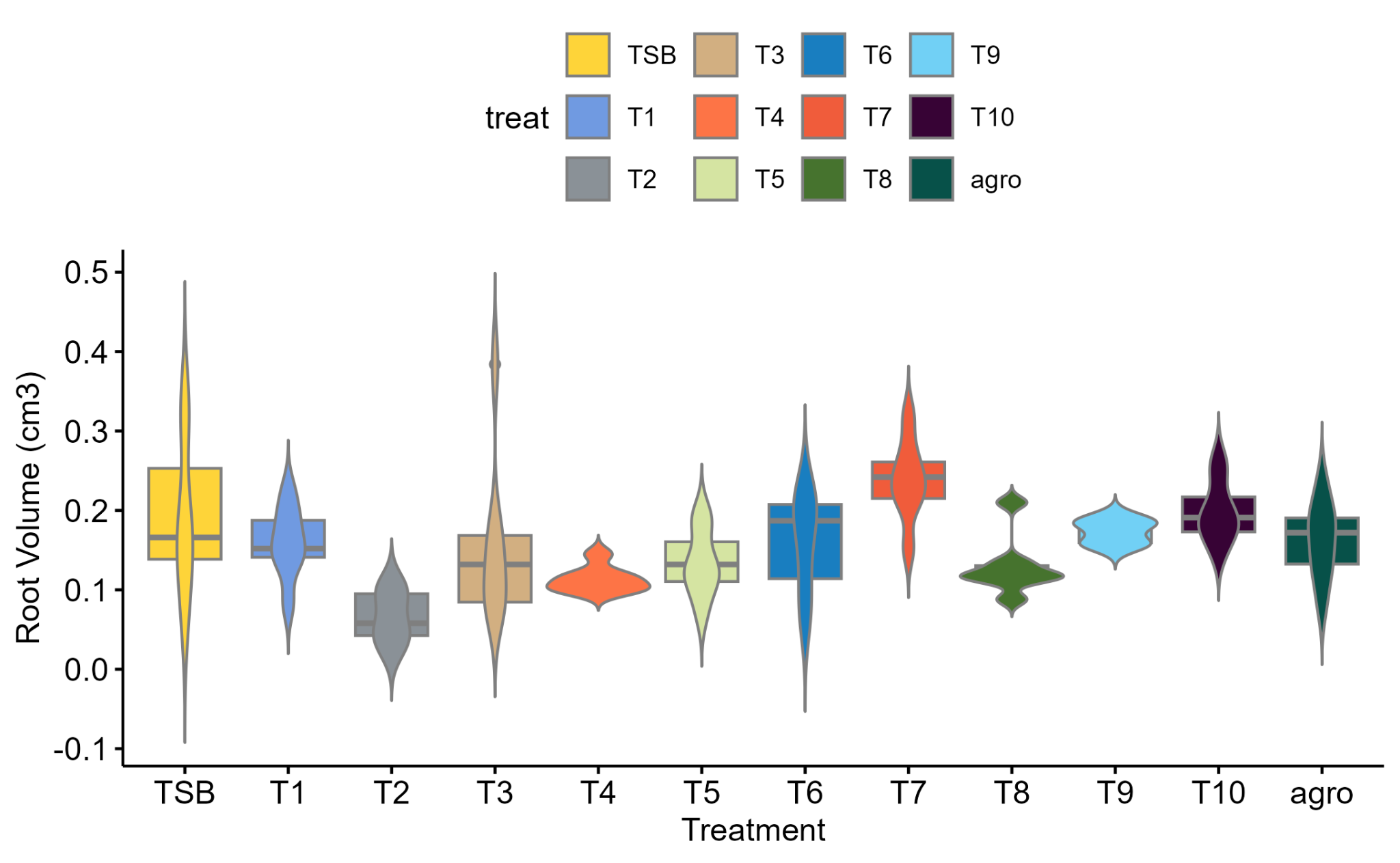

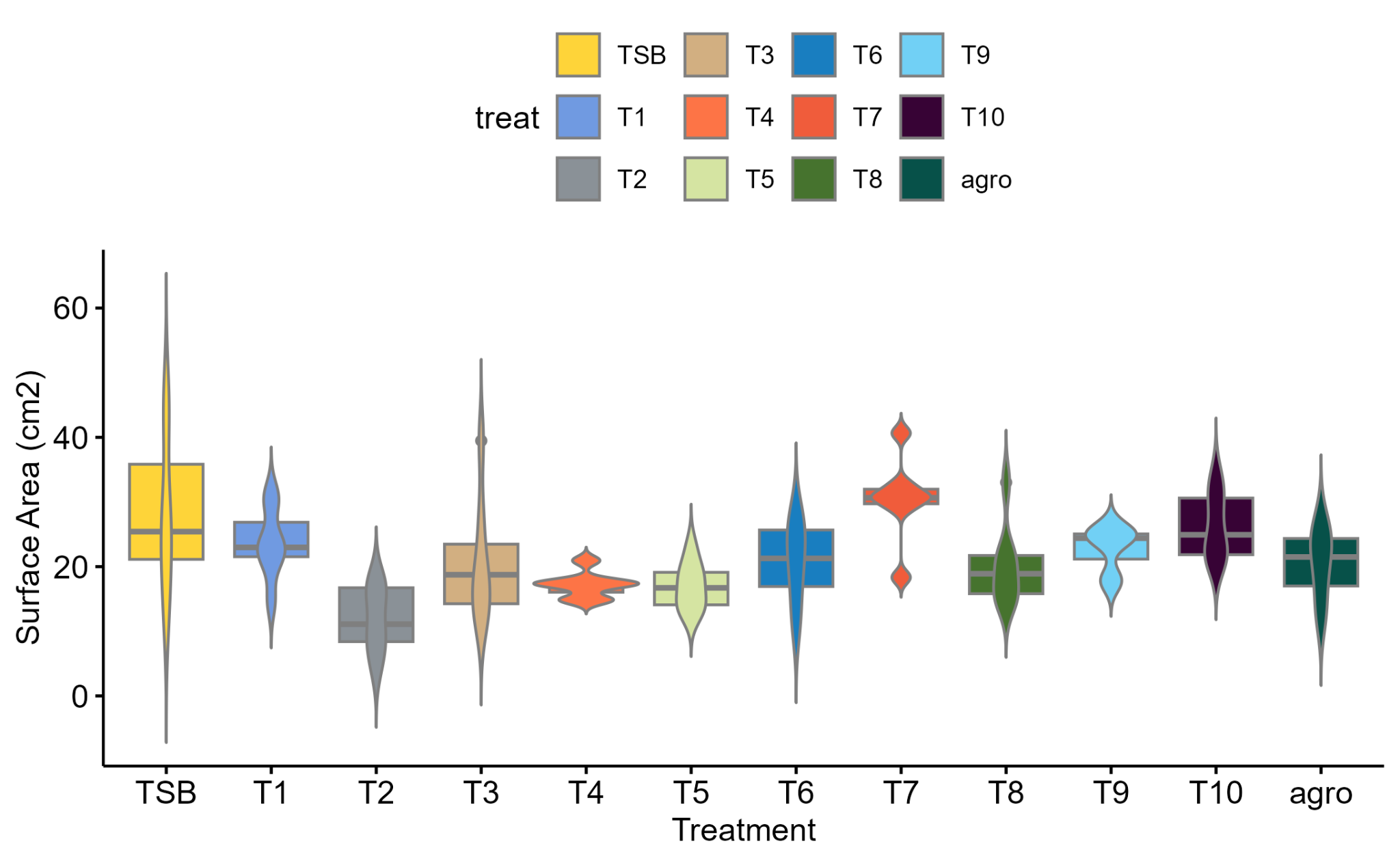

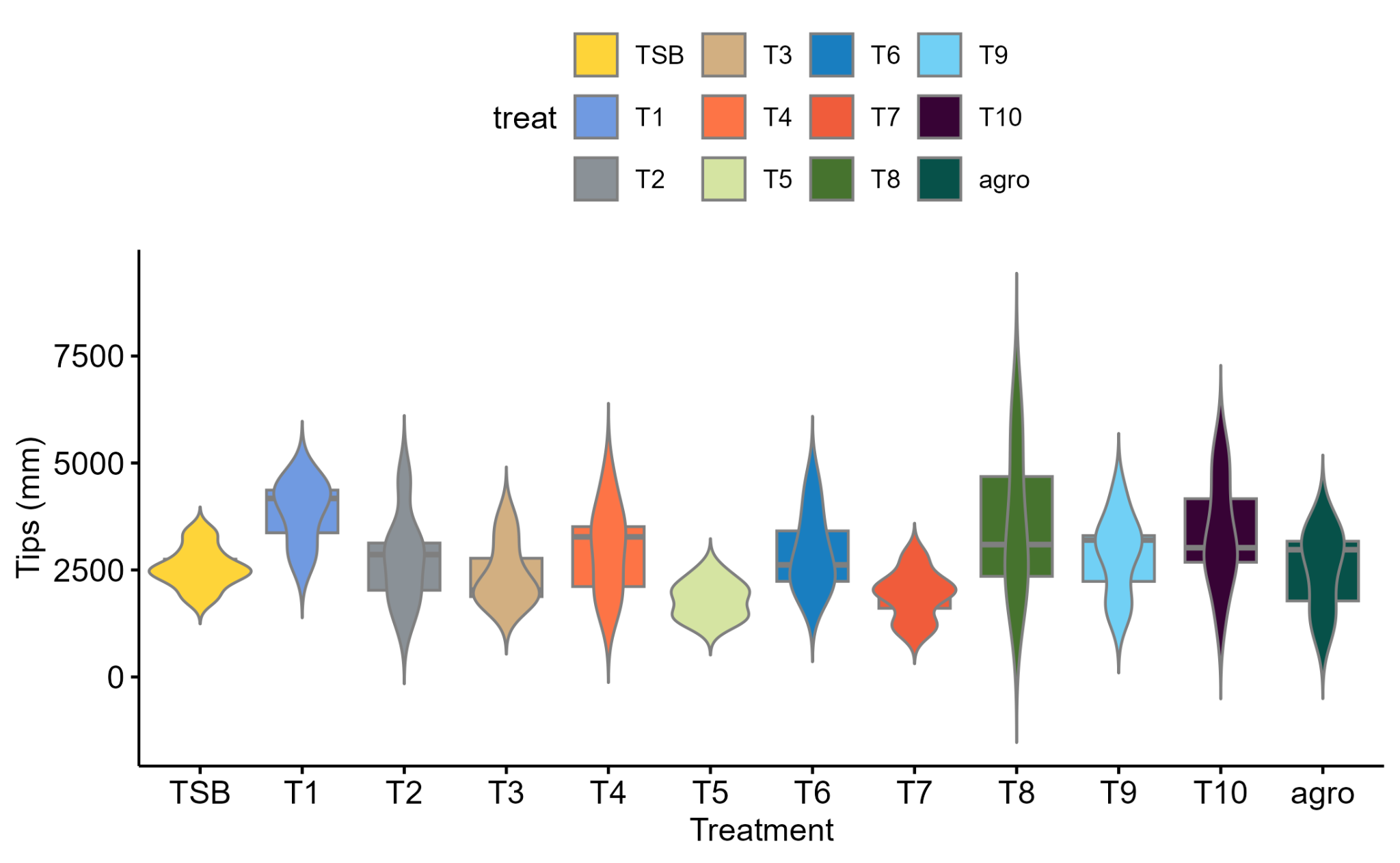

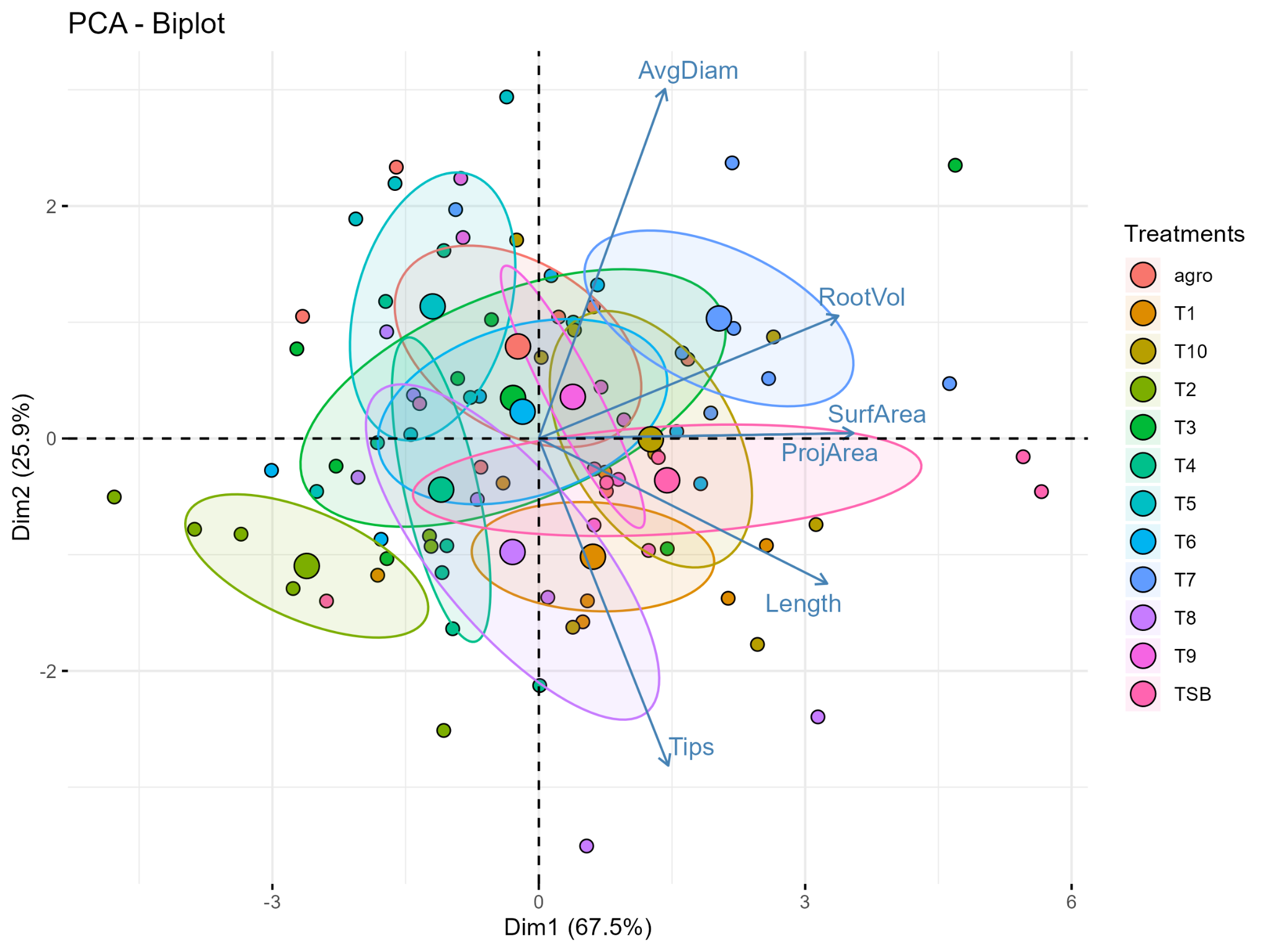

pca biplot

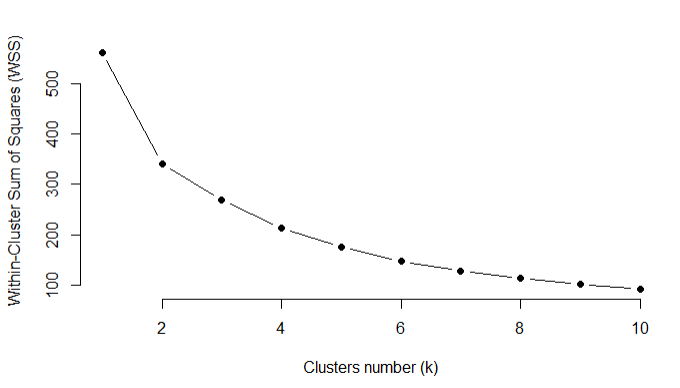

wss kmeans k value=4 elbow

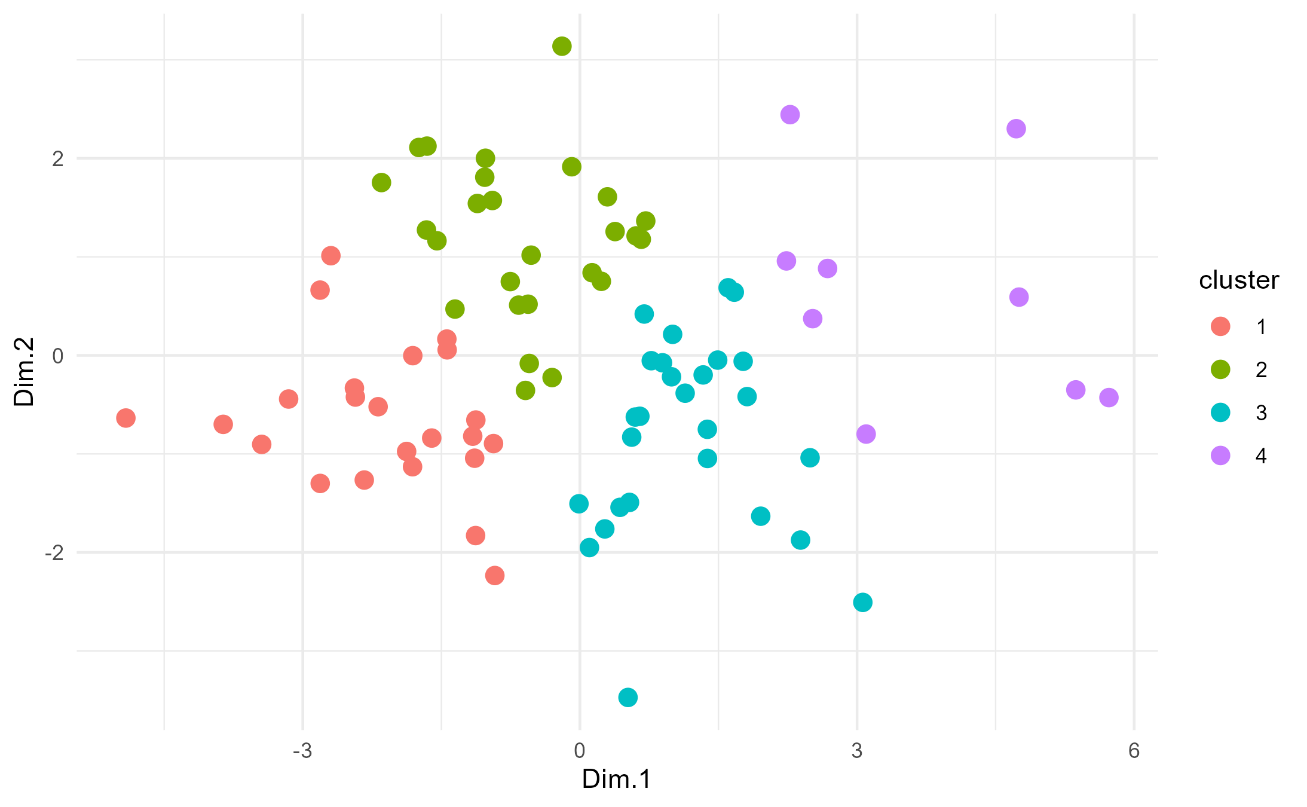

Kmeans clustering. Strains with similar root measurements are colored according to K-means clustering analysis. The optimal number of clusters was determined using the elbow method.

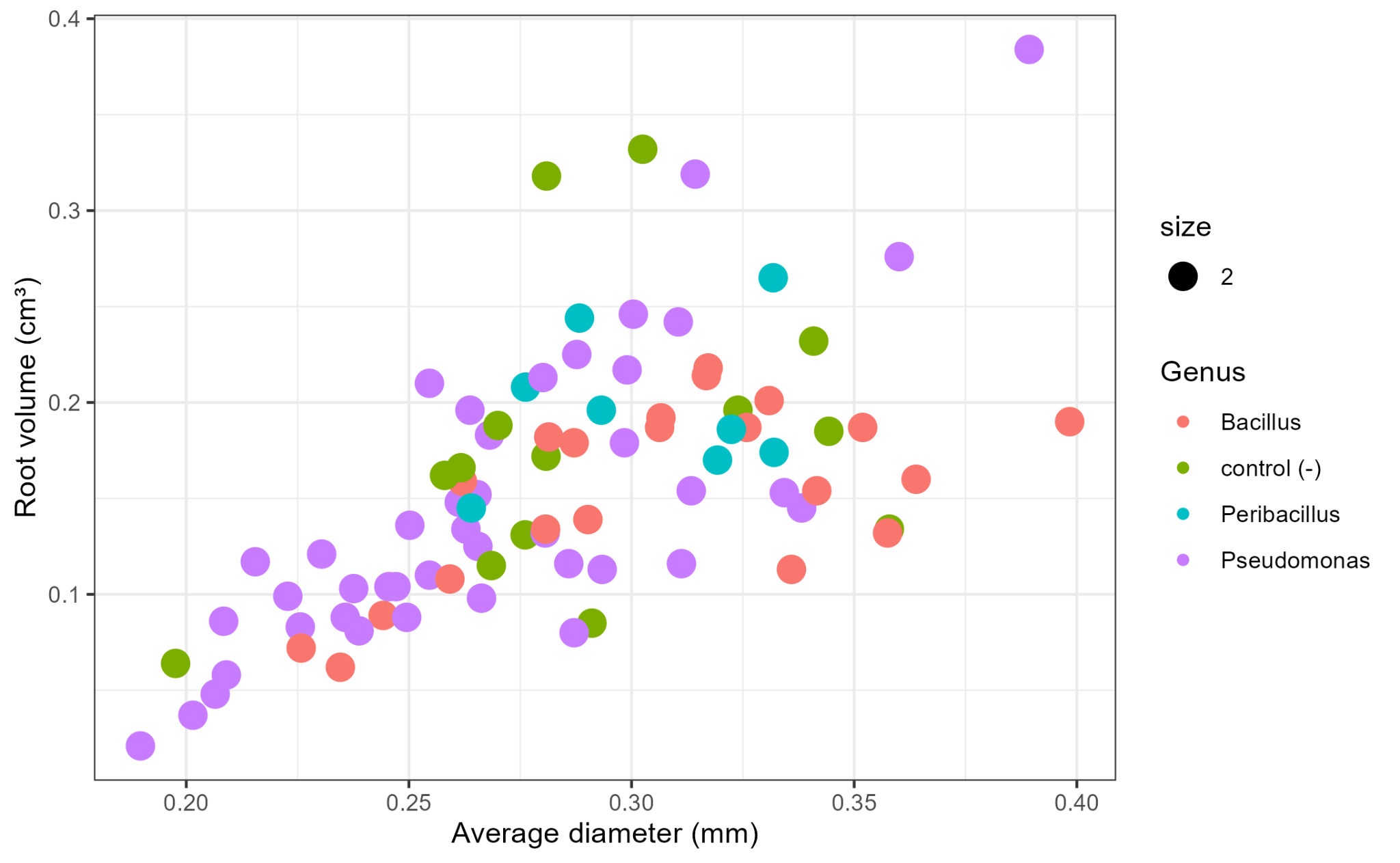

Scatterplot of all treatments and no bacterial controls, positive and negative, based on average diameter and root volume. Color was based on the Genus

| Treatment | Biostimulant (H2O) | | | | | |
| --- | --- | --- | --- | --- | --- | --- |
|  | Length root | Projection area | Surface area | Average diameter | Root volume | Tips number |
| 1 |  |  |  |  |  |  |
| 2 | *** | * | * |  |  |  |
| 3 | * |  |  |  |  |  |
| 4 | * |  |  |  |  |  |
| 5 | *** |  |  |  |  |  |
| 6 | * |  |  |  |  |  |
| 7 |  |  |  | . |  |  |
| 8 |  |  |  |  |  |  |
| 9 |  |  |  |  |  |  |
| 10 |  |  |  |  |  |  |
| control (-) | ** |  |  | . |  |  |
| control (-) |  |  |  |  |  |  |

**Pathogen resistance**

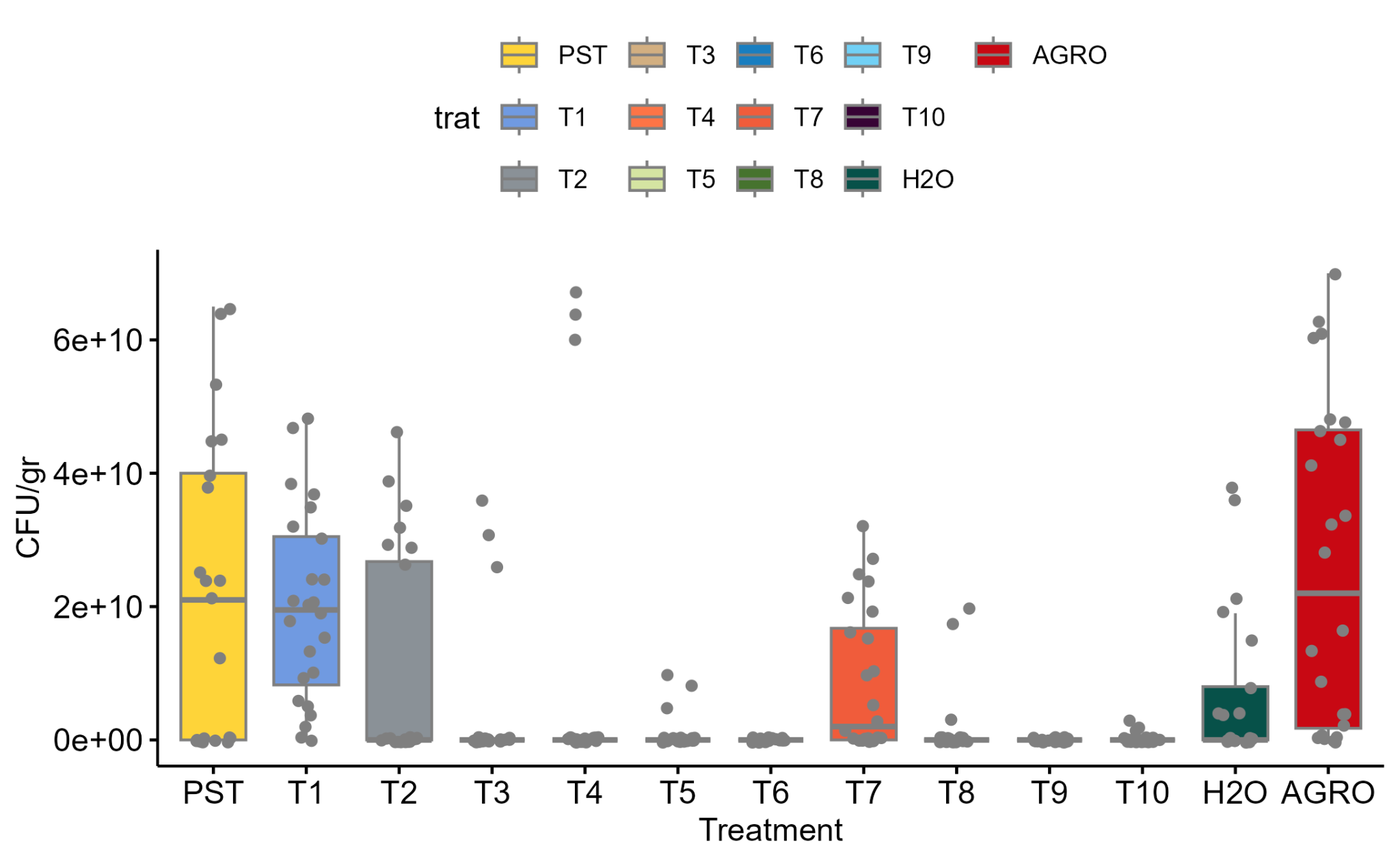

**Figure S14.** Boxplot showing the response in colony-forming unit (CFU) per gram of leaf tissue infected with *Pseudomonas syringae* pv. *tomato* (*Pst*) DC 3000 to bacterial treatments (T). Treatments include: TSB, a medium without *Pst* infection; PST, TSB medium with *Pst* infection; and AGRO, *Agrobacterium tumefaciens*. Statistical differences were assessed using one-way ANOVA (p>0.05) and the Tukey test.

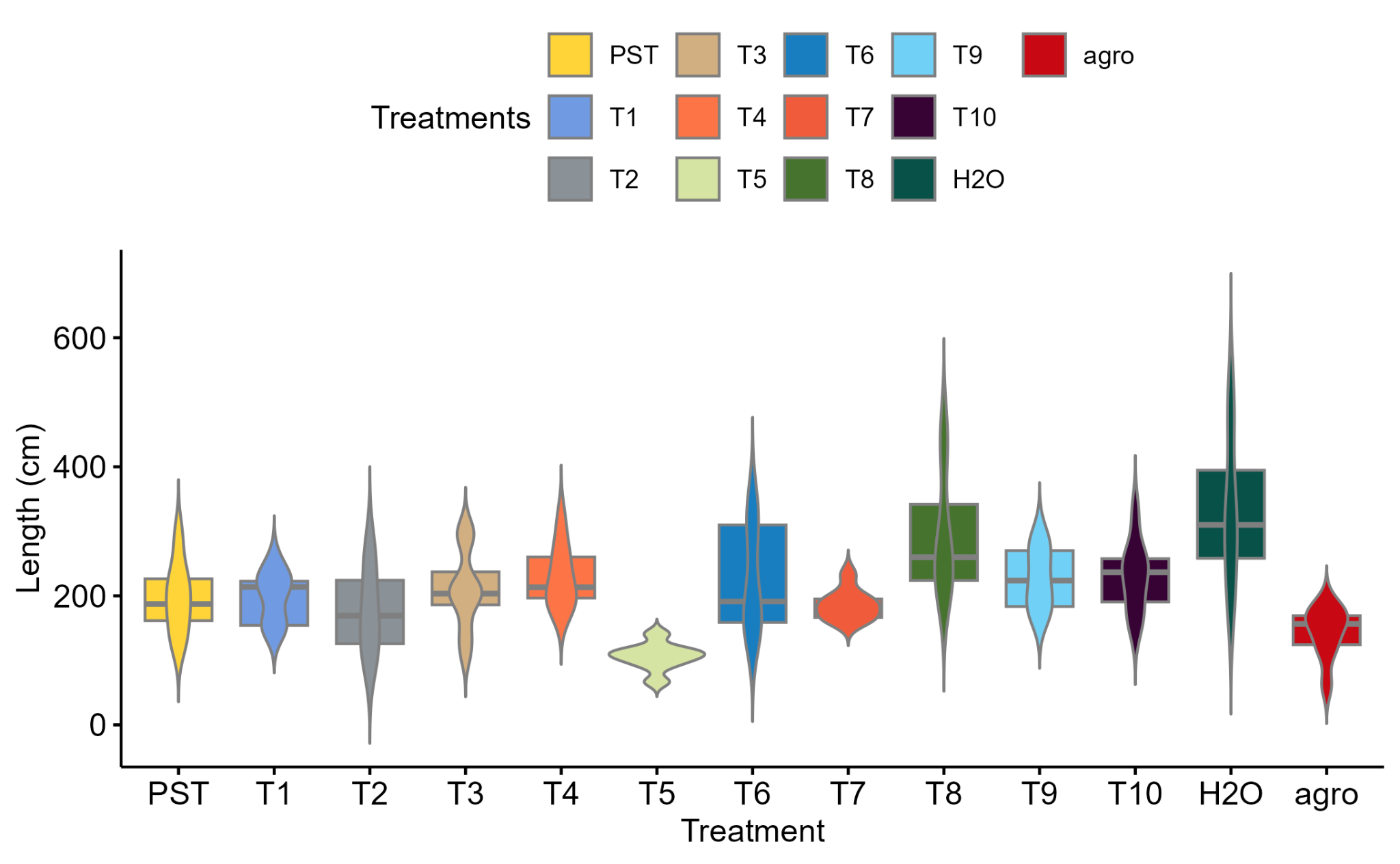

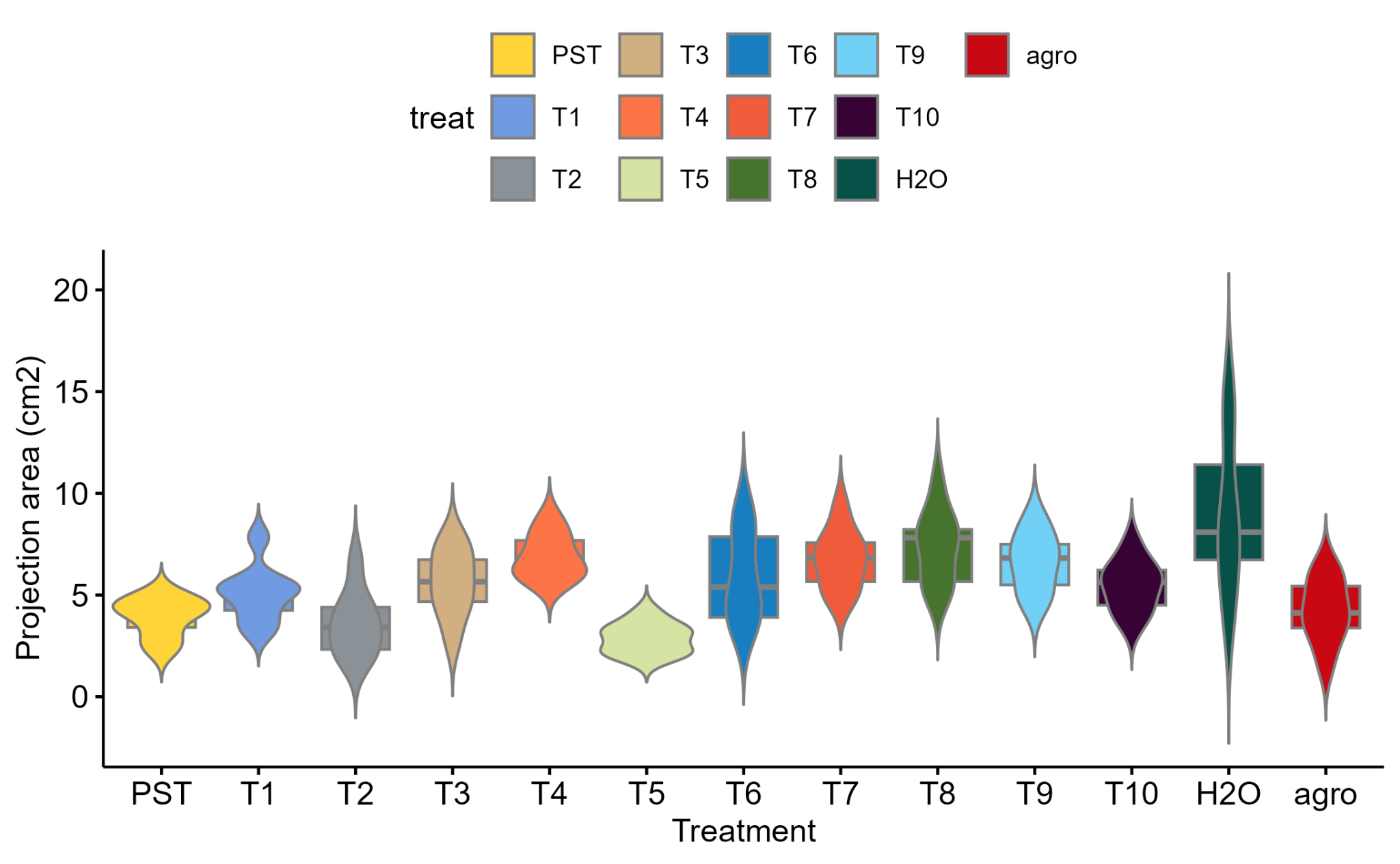

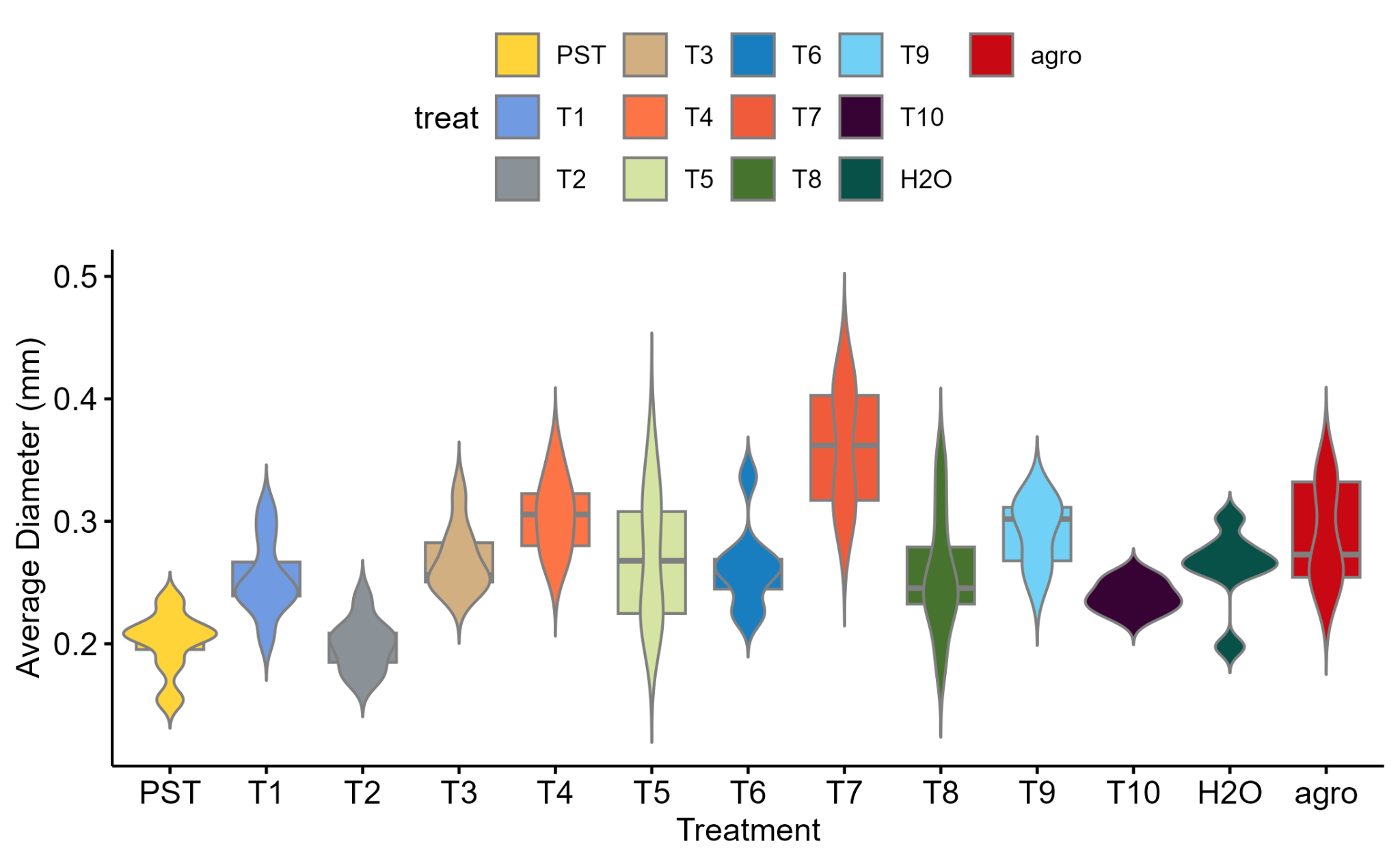

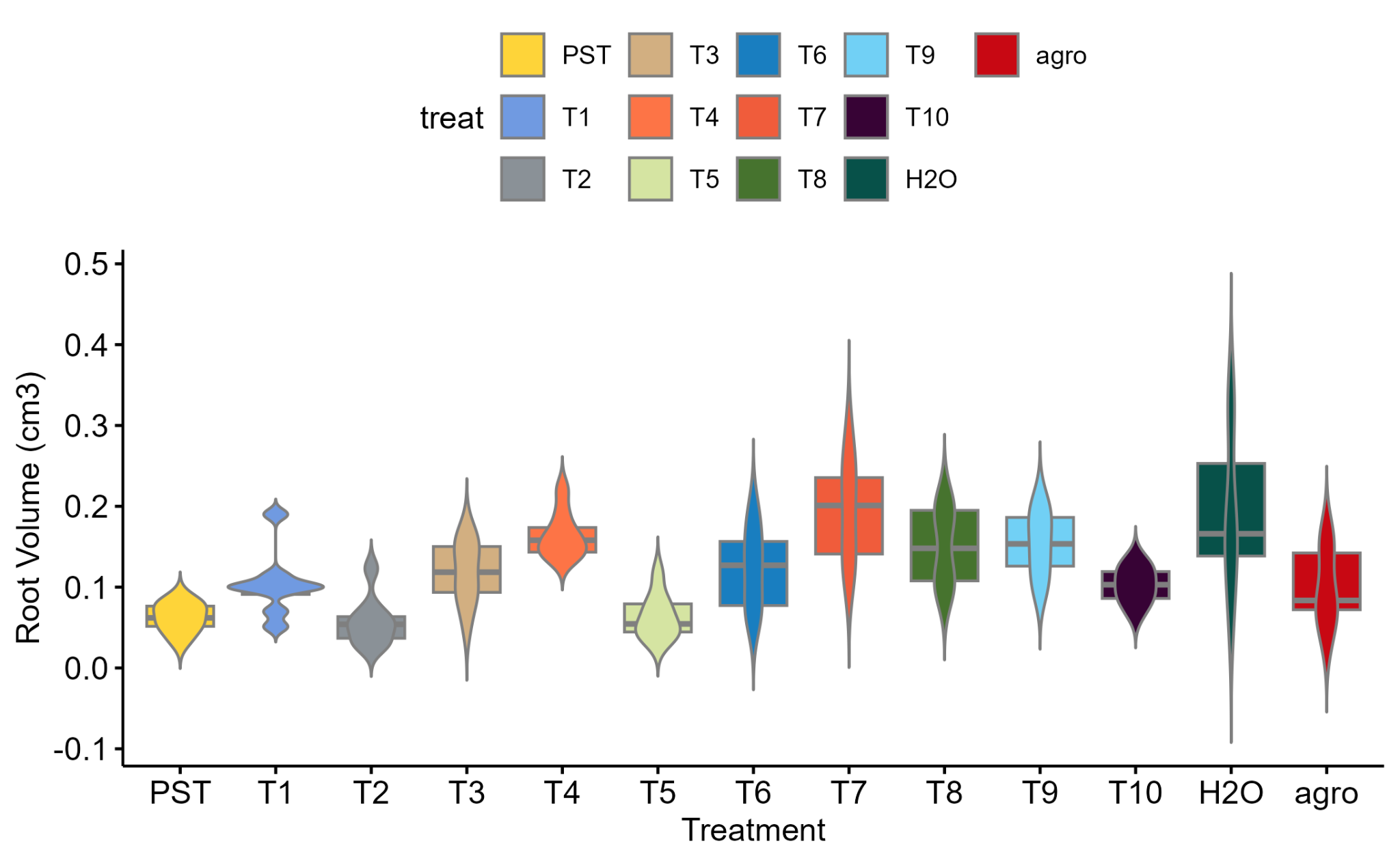

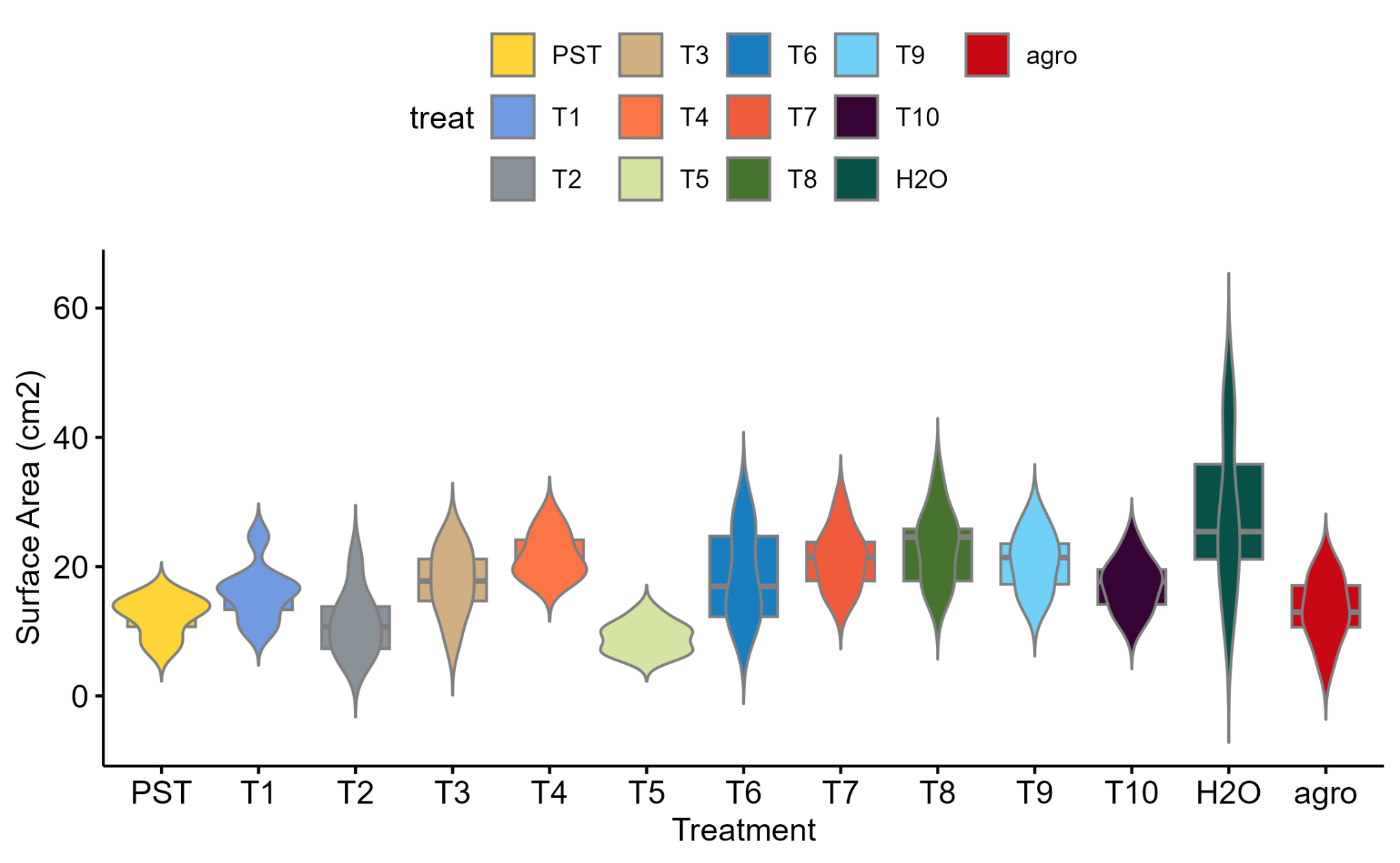

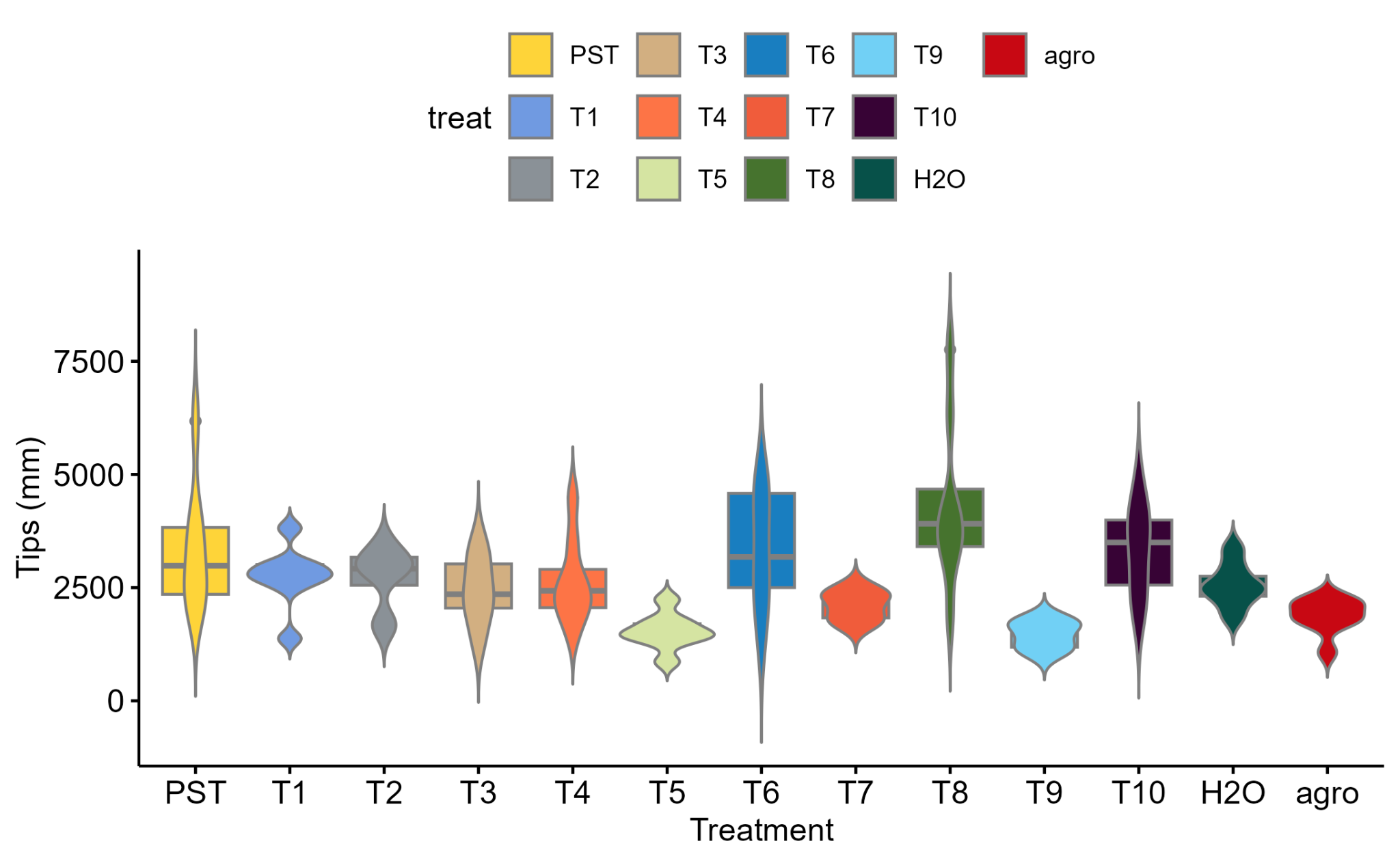

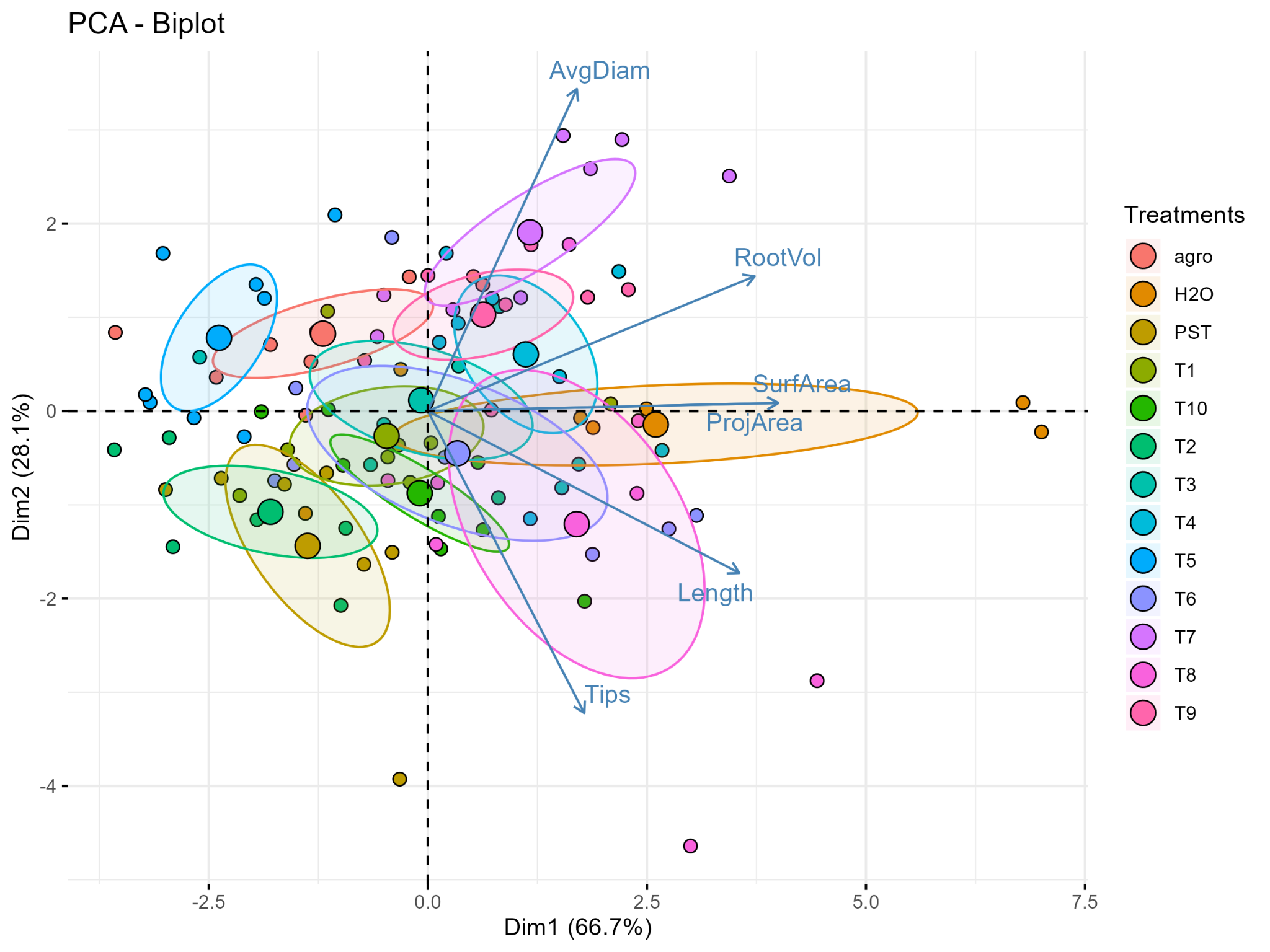

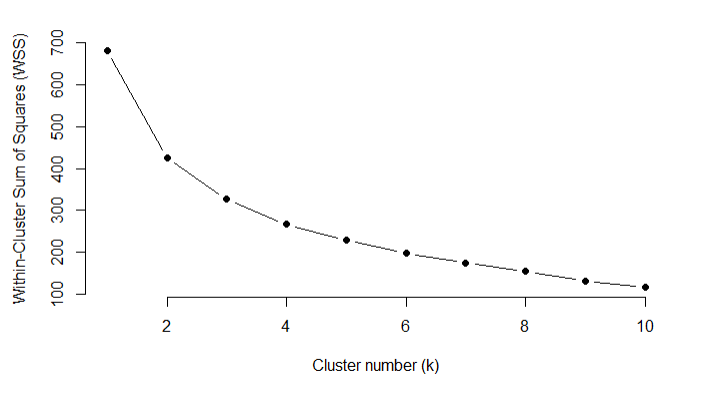

wss kmeans k value=4 elbow

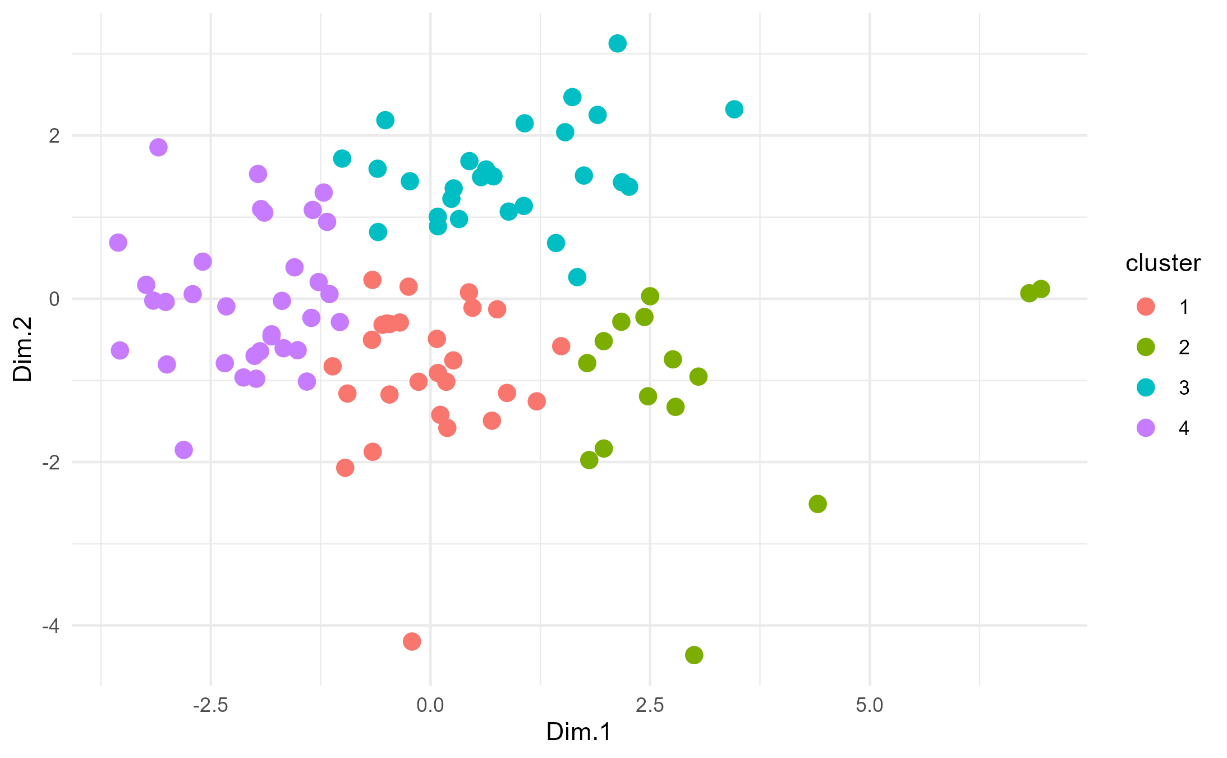

Kmeans clustering. Strains with similar root measurements are colored according to K-means clustering analysis. The optimal number of clusters was determined using the elbow method.

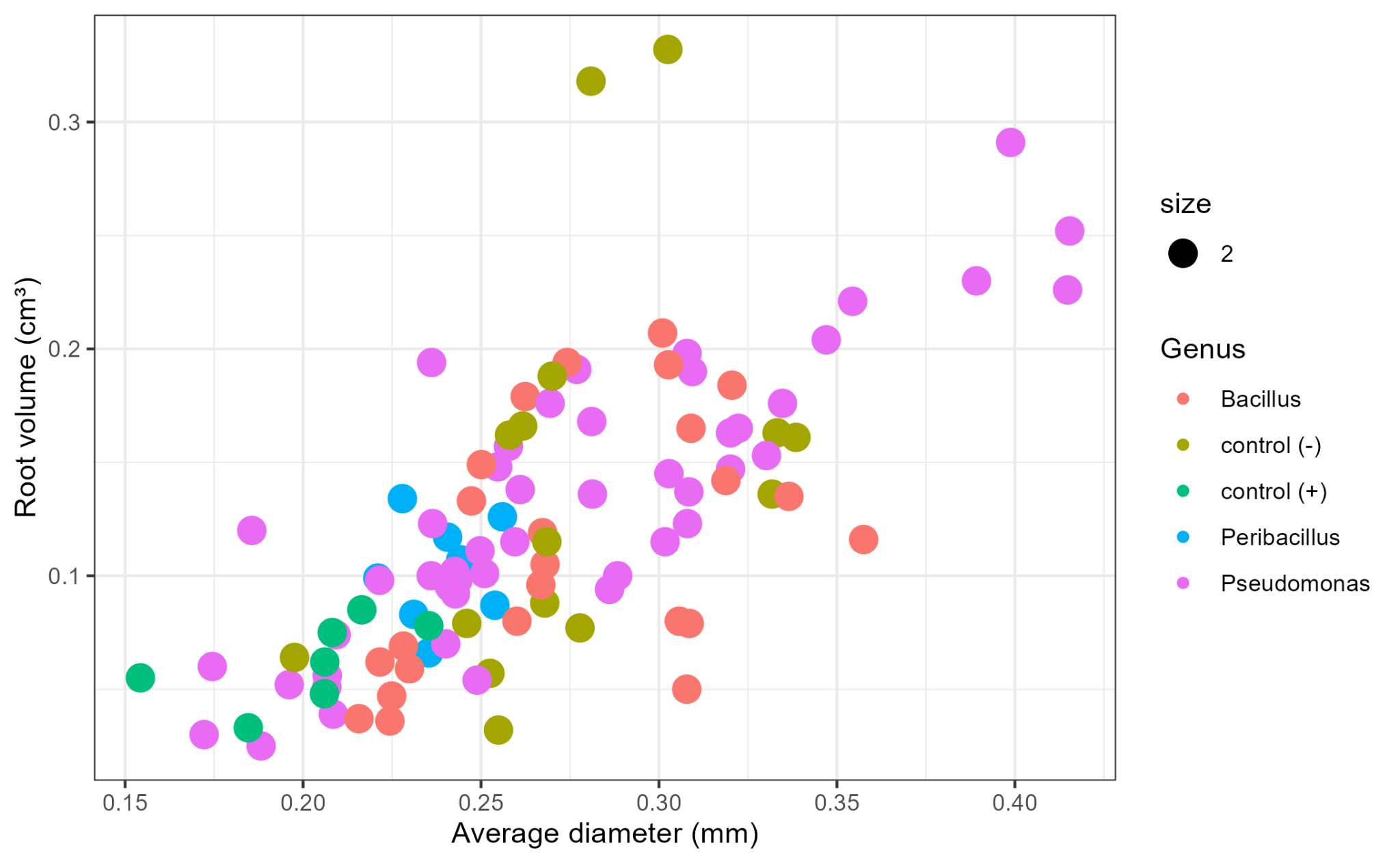

**Fig S15.** Scatterplot of all treatments and no bacterial controls, positive and negative, based on average diameter and root volume. Color was based on the Genus

**Table S2**. Root measurements related to *Pseudomonas syringae* pv. *tomato* (pst) immunity response

| Treatment | Pathogen resistance (pst) | | | | | | |
| --- | --- | --- | --- | --- | --- | --- | --- |
|  | CFU/gr | Length root | Projection area | Surface area | Average diameter | Root volume | Tips number |
| 1 |  |  |  |  | . |  |  |
| 2 |  |  |  |  |  |  |  |
| 3 | * |  |  |  | ** |  |  |
| 4 |  | ** | ** | ** | *** | ** |  |
| 5 | ** |  |  |  | ** |  | ** |
| 6 | *** |  |  |  | * |  |  |
| 7 |  |  | ** | ** | *** | ** |  |
| 8 | ** |  | ** | ** | * | ** |  |
| 9 | *** |  | * | * | *** | ** | ** |
| 10 | ** |  |  |  |  | * |  |
| control (-) |  |  |  |  | *** |  | * |
| control (-) |  |  |  |  | * | * |  |
| control (+) |  |  |  |  |  |  |  |

**Table S1** Profile of PGPR for bacterial strains isolated from soils associated with healthy (S) and asymptomatic (As) tomato plants, their isolation ID, and results from functional assays

| ID | Tomato phenotype | ACC  deaminase | Indolacetic acid (IAA) | Osmotic stress | Nitrogen fixation | Phosphate solubilization | ROS |
| --- | --- | --- | --- | --- | --- | --- | --- |
| AS01 | Asymptomatic | 0 | 0 | 1 | 0 | 0 | 0 |
| AS02 | Asymptomatic | 0 | 0 | 1 | 0 | 0 | 0 |
| AS03 | Asymptomatic | 0 | 0 | 1 | 0 | 0 | 0 |
| AS04 | Asymptomatic | 0 | 1 | 1 | 0 | 0 | 0 |
| AS05 | Asymptomatic | 0 | 0 | 0 | 0 | 0 | 0 |
| AS06 | Asymptomatic | 0 | 0 | 0 | 0 | 0 | 0 |
| AS07 | Asymptomatic | 0 | 0 | 1 | 0 | 0 | 0 |
| AS08 | Asymptomatic | 0 | 0 | 1 | 0 | 0 | 0 |
| AS09 | Asymptomatic | 0 | 0 | 0 | 0 | 0 | 0 |
| AS10 | Asymptomatic | 0 | 0 | 0 | 0 | 0 | 0 |
| AS11 | Asymptomatic | 1 | 0 | 0 | 0 | 0 | 0 |
| AS12 | Asymptomatic | 1 | 0 | 1 | 0 | 0 | 0 |
| AS13 | Asymptomatic | 1 | 0 | 1 | 0 | 0 | 1 |
| AS14 | Asymptomatic | 0 | 0 | 1 | 0 | 0 | 0 |
| AS15 | Asymptomatic | 0 | 0 | 1 | 0 | 0 | 0 |
| AS16 | Asymptomatic | 0 | 0 | 1 | 0 | 0 | 0 |
| AS17 | Asymptomatic | 0 | 0 | 1 | 0 | 0 | 0 |
| AS18 | Asymptomatic | 0 | 0 | 1 | 0 | 0 | 0 |
| AS19 | Asymptomatic | 0 | 0 | 1 | 0 | 0 | 0 |
| AS20 | Asymptomatic | 0 | 0 | 0 | 0 | 0 | 0 |
| AS21 | Asymptomatic | 0 | 0 | 0 | 0 | 0 | 0 |
| AS22 | Asymptomatic | 0 | 0 | 0 | 0 | 0 | 0 |
| AS23 | Asymptomatic | 0 | 1 | 1 | 0 | 0 | 0 |
| AS24 | Asymptomatic | 0 | 0 | 0 | 0 | 0 | 0 |
| AS25 | Asymptomatic | 0 | 0 | 1 | 0 | 0 | 0 |
| AS26 | Asymptomatic | 0 | 0 | 1 | 0 | 0 | 0 |
| AS27 | Asymptomatic | 0 | 0 | 1 | 0 | 0 | 0 |
| AS28 | Asymptomatic | 1 | 0 | 1 | 0 | 0 | 0 |
| AS29 | Asymptomatic | 0 | 0 | 1 | 0 | 0 | 0 |
| AS30 | Asymptomatic | 0 | 0 | 0 | 0 | 0 | 0 |
| AS31 | Asymptomatic | 0 | 1 | 1 | 0 | 0 | 0 |
| AS32 | Asymptomatic | 0 | 0 | 0 | 0 | 0 | 0 |
| AS33 | Asymptomatic | 0 | 0 | 0 | 0 | 0 | 0 |
| AS34 | Asymptomatic | 0 | 0 | 1 | 0 | 0 | 0 |
| AS35 | Asymptomatic | 0 | 0 | 0 | 0 | 0 | 0 |
| AS36 | Asymptomatic | 0 | 0 | 1 | 0 | 0 | 0 |
| AS37 | Asymptomatic | 0 | 0 | 1 | 0 | 0 | 0 |
| AS38 | Asymptomatic | 0 | 0 | 0 | 0 | 0 | 0 |
| AS39 | Asymptomatic | 0 | 1 | 1 | 0 | 0 | 1 |
| AS40 | Asymptomatic | 0 | 0 | 1 | 0 | 0 | 0 |
| AS41 | Asymptomatic | 0 | 0 | 1 | 0 | 0 | 0 |
| AS42 | Asymptomatic | 0 | 0 | 1 | 0 | 0 | 0 |
| AS43 | Asymptomatic | 0 | 1 | 1 | 0 | 0 | 1 |
| AS44 | Asymptomatic | 1 | 0 | 0 | 0 | 0 | 0 |
| AS45 | Asymptomatic | 0 | 0 | 0 | 0 | 0 | 0 |
| AS46 | Asymptomatic | 1 | 1 | 1 | 0 | 0 | 1 |
| AS47 | Asymptomatic | 0 | 1 | 1 | 0 | 0 | 1 |
| AS48 | Asymptomatic | 0 | 0 | 1 | 0 | 0 | 0 |
| AS49 | Asymptomatic | 0 | 0 | 1 | 0 | 0 | 0 |
| AS50 | Asymptomatic | 0 | 0 | 1 | 0 | 0 | 0 |
| AS51 | Asymptomatic | 0 | 0 | 1 | 0 | 0 | 0 |
| AS52 | Asymptomatic | 0 | 1 | 0 | 0 | 0 | 0 |
| AS53 | Asymptomatic | 0 | 0 | 1 | 0 | 0 | 0 |
| AS54 | Asymptomatic | 0 | 1 | 1 | 0 | 0 | 1 |
| AS55 | Asymptomatic | 0 | 0 | 1 | 0 | 0 | 0 |
| AS56 | Asymptomatic | 0 | 0 | 1 | 0 | 0 | 0 |
| AS57 | Asymptomatic | 0 | 0 | 0 | 0 | 0 | 0 |
| AS58 | Asymptomatic | 0 | 0 | 0 | 0 | 0 | 0 |
| AS59 | Asymptomatic | 1 | 0 | 1 | 0 | 0 | 1 |
| AS60 | Asymptomatic | 1 | 0 | 1 | 0 | 0 | 1 |
| AS61 | Asymptomatic | 1 | 0 | 0 | 0 | 1 | 0 |
| AS62 | Asymptomatic | 0 | 0 | 0 | 0 | 0 | 0 |
| AS63 | Asymptomatic | 0 | 0 | 0 | 0 | 0 | 0 |
| AS64 | Asymptomatic | 0 | 0 | 1 | 0 | 0 | 0 |
| AS65 | Asymptomatic | 0 | 0 | 0 | 0 | 0 | 0 |
| AS66 | Asymptomatic | 0 | 1 | 1 | 0 | 0 | 1 |
| AS67 | Asymptomatic | 1 | 0 | 1 | 0 | 0 | 1 |
| AS68 | Asymptomatic | 0 | 0 | 0 | 0 | 0 | 0 |
| AS69 | Asymptomatic | 0 | 0 | 1 | 0 | 0 | 0 |
| AS70 | Asymptomatic | 0 | 0 | 0 | 0 | 0 | 0 |
| AS71 | Asymptomatic | 0 | 0 | 1 | 0 | 0 | 0 |
| AS72 | Asymptomatic | 0 | 0 | 0 | 0 | 0 | 0 |
| AS73 | Asymptomatic | 0 | 1 | 1 | 0 | 0 | 0 |
| AS74 | Asymptomatic | 0 | 0 | 1 | 0 | 0 | 0 |
| AS75 | Asymptomatic | 0 | 0 | 0 | 0 | 0 | 0 |
| AS76 | Asymptomatic | 0 | 0 | 0 | 0 | 0 | 0 |
| AS77 | Asymptomatic | 1 | 0 | 1 | 0 | 0 | 1 |
| AS78 | Asymptomatic | 0 | 0 | 1 | 0 | 0 | 0 |
| AS79 | Asymptomatic | 0 | 0 | 0 | 0 | 0 | 0 |
| AS80 | Asymptomatic | 0 | 0 | 1 | 0 | 0 | 0 |
| AS81 | Asymptomatic | 0 | 1 | 1 | 0 | 0 | 1 |
| AS82 | Asymptomatic | 0 | 0 | 1 | 0 | 0 | 0 |
| AS83 | Asymptomatic | 0 | 0 | 0 | 0 | 0 | 0 |
| AS84 | Asymptomatic | 0 | 0 | 0 | 0 | 0 | 0 |
| AS85 | Asymptomatic | 0 | 1 | 1 | 0 | 0 | 1 |
| AS86 | Asymptomatic | 0 | 0 | 0 | 0 | 0 | 0 |
| AS87 | Asymptomatic | 0 | 0 | 0 | 1 | 1 | 1 |
| AS88 | Asymptomatic | 0 | 1 | 0 | 0 | 0 | 1 |
| AS89 | Asymptomatic | 0 | 0 | 1 | 1 | 1 | 1 |
| AS90 | Asymptomatic | 0 | 0 | 1 | 0 | 0 | 0 |
| AS91 | Asymptomatic | 0 | 0 | 1 | 0 | 0 | 0 |
| AS92 | Asymptomatic | 0 | 0 | 0 | 0 | 0 | 0 |
| AS93 | Asymptomatic | 0 | 0 | 1 | 0 | 0 | 0 |
| AS94 | Asymptomatic | 1 | 0 | 1 | 0 | 0 | 0 |
| AS95 | Asymptomatic | 0 | 0 | 1 | 0 | 0 | 0 |
| AS96 | Asymptomatic | 0 | 0 | 1 | 0 | 0 | 0 |
| AS97 | Asymptomatic | 0 | 0 | 0 | 0 | 0 | 0 |
| AS98 | Asymptomatic | 0 | 0 | 0 | 0 | 0 | 0 |
| AS99 | Asymptomatic | 0 | 0 | 1 | 0 | 0 | 0 |
| AS100 | Asymptomatic | 0 | 0 | 0 | 0 | 0 | 0 |
| AS101 | Asymptomatic | 0 | 1 | 0 | 0 | 0 | 0 |
| AS102 | Asymptomatic | 0 | 0 | 1 | 0 | 0 | 0 |
| AS103 | Asymptomatic | 0 | 0 | 0 | 0 | 0 | 0 |
| AS104 | Asymptomatic | 0 | 0 | 1 | 0 | 0 | 1 |
| AS105 | Asymptomatic | 0 | 0 | 1 | 0 | 0 | 0 |
| AS106 | Asymptomatic | 0 | 0 | 1 | 0 | 0 | 0 |
| AS107 | Asymptomatic | 0 | 0 | 1 | 0 | 0 | 0 |
| AS108 | Asymptomatic | 0 | 0 | 1 | 0 | 0 | 0 |
| AS109 | Asymptomatic | 0 | 1 | 1 | 0 | 0 | 1 |
| AS110 | Asymptomatic | 0 | 0 | 0 | 0 | 0 | 0 |
| AS111 | Asymptomatic | 0 | 0 | 0 | 0 | 0 | 0 |
| AS112 | Asymptomatic | 0 | 1 | 1 | 0 | 0 | 0 |
| AS113 | Asymptomatic | 0 | 0 | 1 | 0 | 0 | 0 |
| AS114 | Asymptomatic | 0 | 0 | 1 | 0 | 0 | 0 |
| AS115 | Asymptomatic | 1 | 0 | 1 | 0 | 0 | 1 |
| AS116 | Asymptomatic | 1 | 0 | 1 | 0 | 0 | 1 |
| AS117 | Asymptomatic | 0 | 0 | 0 | 0 | 0 | 0 |
| AS118 | Asymptomatic | 0 | 0 | 1 | 0 | 0 | 0 |
| AS119 | Asymptomatic | 0 | 0 | 1 | 0 | 0 | 0 |
| AS120 | Asymptomatic | 0 | 0 | 1 | 0 | 0 | 0 |
| AS121 | Asymptomatic | 0 | 0 | 0 | 0 | 0 | 0 |
| AS122 | Asymptomatic | 0 | 0 | 1 | 0 | 0 | 0 |
| AS123 | Asymptomatic | 0 | 0 | 1 | 0 | 0 | 0 |
| AS124 | Asymptomatic | 0 | 0 | 1 | 0 | 0 | 0 |
| AS125 | Asymptomatic | 0 | 0 | 0 | 0 | 0 | 0 |
| AS126 | Asymptomatic | 0 | 0 | 1 | 0 | 0 | 0 |
| AS127 | Asymptomatic | 0 | 0 | 0 | 0 | 0 | 0 |
| AS128 | Asymptomatic | 0 | 0 | 1 | 0 | 0 | 0 |
| AS129 | Asymptomatic | 0 | 0 | 1 | 0 | 0 | 0 |
| S01 | Healthy | 0 | 0 | 1 | 0 | 0 | 0 |
| S02 | Healthy | 0 | 0 | 1 | 0 | 0 | 0 |
| S03 | Healthy | 0 | 0 | 1 | 0 | 0 | 0 |
| S04 | Healthy | 0 | 0 | 0 | 0 | 0 | 0 |
| S05 | Healthy | 1 | 0 | 1 | 0 | 0 | 0 |
| S06 | Healthy | 0 | 0 | 1 | 0 | 1 | 0 |
| S07 | Healthy | 0 | 0 | 1 | 0 | 0 | 0 |
| S08 | Healthy | 0 | 0 | 1 | 0 | 0 | 0 |
| S09 | Healthy | 0 | 1 | 1 | 0 | 0 | 0 |
| S10 | Healthy | 0 | 0 | 1 | 0 | 0 | 0 |
| S11 | Healthy | 0 | 0 | 0 | 0 | 0 | 0 |
| S12 | Healthy | 0 | 0 | 0 | 0 | 0 | 0 |
| S13 | Healthy | 0 | 0 | 1 | 0 | 0 | 0 |
| S14 | Healthy | 0 | 0 | 1 | 0 | 0 | 0 |
| S15 | Healthy | 0 | 0 | 1 | 0 | 0 | 0 |
| S16 | Healthy | 0 | 0 | 1 | 0 | 0 | 0 |
| S17 | Healthy | 0 | 0 | 1 | 0 | 0 | 0 |
| S18 | Healthy | 1 | 0 | 1 | 1 | 1 | 0 |
| S19 | Healthy | 0 | 0 | 1 | 0 | 0 | 0 |
| S20 | Healthy | 0 | 0 | 1 | 0 | 0 | 0 |
| S21 | Healthy | 0 | 0 | 1 | 0 | 0 | 0 |
| S22 | Healthy | 0 | 0 | 1 | 1 | 1 | 1 |
| S23 | Healthy | 0 | 0 | 1 | 0 | 0 | 0 |
| S24 | Healthy | 0 | 0 | 0 | 0 | 0 | 0 |
| S25 | Healthy | 0 | 0 | 1 | 0 | 0 | 0 |
| S26 | Healthy | 0 | 0 | 0 | 0 | 0 | 0 |
| S27 | Healthy | 0 | 0 | 1 | 0 | 0 | 0 |
| S28 | Healthy | 0 | 0 | 1 | 0 | 0 | 0 |
| S29 | Healthy | 0 | 0 | 1 | 0 | 0 | 0 |
| S30 | Healthy | 0 | 0 | 1 | 0 | 0 | 0 |
| S31 | Healthy | 0 | 0 | 1 | 0 | 0 | 0 |
| S32 | Healthy | 1 | 0 | 1 | 0 | 0 | 0 |
| S33 | Healthy | 1 | 0 | 1 | 0 | 0 | 0 |
| S34 | Healthy | 0 | 0 | 1 | 0 | 0 | 0 |
| S35 | Healthy | 0 | 0 | 0 | 0 | 0 | 0 |
| S36 | Healthy | 0 | 1 | 1 | 0 | 0 | 0 |
| S37 | Healthy | 0 | 0 | 0 | 0 | 0 | 0 |
| S38 | Healthy | 0 | 0 | 1 | 0 | 0 | 0 |
| S39 | Healthy | 0 | 0 | 1 | 0 | 0 | 0 |
| S40 | Healthy | 0 | 0 | 1 | 0 | 0 | 0 |
| S41 | Healthy | 0 | 0 | 1 | 0 | 0 | 0 |
| S42 | Healthy | 0 | 0 | 1 | 0 | 0 | 0 |
| S43 | Healthy | 0 | 0 | 1 | 0 | 0 | 0 |
| S44 | Healthy | 0 | 0 | 0 | 0 | 0 | 0 |
| S45 | Healthy | 0 | 0 | 1 | 0 | 0 | 0 |
| S46 | Healthy | 0 | 1 | 1 | 0 | 0 | 0 |
| S47 | Healthy | 1 | 0 | 1 | 0 | 0 | 0 |
| S48 | Healthy | 0 | 0 | 0 | 0 | 0 | 0 |
| S49 | Healthy | 0 | 0 | 0 | 0 | 0 | 0 |
| S50 | Healthy | 0 | 0 | 1 | 0 | 0 | 0 |
| S51 | Healthy | 0 | 1 | 1 | 0 | 0 | 0 |
| S52 | Healthy | 0 | 0 | 1 | 0 | 0 | 0 |
| S53 | Healthy | 0 | 0 | 0 | 0 | 0 | 0 |
| S54 | Healthy | 0 | 0 | 1 | 0 | 0 | 0 |
| S55 | Healthy | 0 | 0 | 1 | 0 | 0 | 0 |
| S56 | Healthy | 0 | 0 | 1 | 0 | 0 | 0 |
| S57 | Healthy | 0 | 0 | 1 | 0 | 0 | 0 |
| S58 | Healthy | 0 | 0 | 1 | 0 | 0 | 0 |
| S59 | Healthy | 0 | 0 | 1 | 0 | 0 | 0 |
| S60 | Healthy | 0 | 0 | 0 | 0 | 0 | 0 |
| S61 | Healthy | 0 | 0 | 1 | 0 | 0 | 0 |
| S62 | Healthy | 0 | 0 | 0 | 0 | 0 | 0 |
| S63 | Healthy | 0 | 0 | 1 | 0 | 0 | 0 |
| S64 | Healthy | 0 | 0 | 1 | 0 | 0 | 0 |
| S65 | Healthy | 0 | 0 | 0 | 1 | 1 | 0 |
| S66 | Healthy | 1 | 1 | 0 | 0 | 0 | 0 |
| S67 | Healthy | 0 | 0 | 0 | 0 | 0 | 0 |
| S68 | Healthy | 0 | 0 | 0 | 0 | 0 | 0 |
| S69 | Healthy | 0 | 0 | 1 | 0 | 0 | 0 |
| S70 | Healthy | 0 | 0 | 1 | 0 | 0 | 0 |
| S71 | Healthy | 0 | 0 | 1 | 0 | 0 | 0 |
| S72 | Healthy | 1 | 0 | 1 | 1 | 1 | 0 |
| S73 | Healthy | 0 | 1 | 1 | 0 | 0 | 0 |
| S74 | Healthy | 0 | 0 | 1 | 0 | 0 | 0 |
| S75 | Healthy | 0 | 0 | 1 | 0 | 0 | 0 |
| S76 | Healthy | 0 | 0 | 1 | 0 | 0 | 0 |
| S77 | Healthy | 0 | 0 | 1 | 0 | 0 | 0 |
| S78 | Healthy | 1 | 0 | 1 | 0 | 0 | 0 |
| S79 | Healthy | 0 | 0 | 1 | 0 | 0 | 0 |
| S80 | Healthy | 0 | 0 | 1 | 0 | 0 | 0 |
| S81 | Healthy | 0 | 0 | 1 | 0 | 0 | 0 |
| S82 | Healthy | 0 | 0 | 1 | 0 | 0 | 0 |
| S83 | Healthy | 0 | 0 | 1 | 0 | 0 | 0 |
| S84 | Healthy | 0 | 0 | 1 | 0 | 0 | 0 |
| S85 | Healthy | 0 | 0 | 0 | 0 | 0 | 0 |
| S86 | Healthy | 0 | 0 | 1 | 0 | 0 | 0 |
| S87 | Healthy | 1 | 0 | 0 | 0 | 0 | 0 |
| S88 | Healthy | 0 | 0 | 1 | 0 | 0 | 0 |
| S89 | Healthy | 0 | 0 | 1 | 0 | 0 | 0 |
| S90 | Healthy | 0 | 0 | 1 | 0 | 0 | 0 |
| S91 | Healthy | 0 | 0 | 0 | 0 | 0 | 0 |
| S92 | Healthy | 0 | 0 | 1 | 0 | 0 | 0 |
| S93 | Healthy | 0 | 0 | 1 | 0 | 0 | 0 |
| S94 | Healthy | 1 | 0 | 0 | 0 | 1 | 0 |
